# The reactivity of an unusual amidase may explain colibactin’s DNA cross-linking activity

**DOI:** 10.1101/567248

**Authors:** Yindi Jiang, Alessia Stornetta, Peter W. Villalta, Matthew R. Wilson, Paul D. Boudreau, Li Zha, Silvia Balbo, Emily P. Balskus

## Abstract

Certain commensal and pathogenic bacteria produce colibactin, a small molecule genotoxin that causes interstrand cross-links in host cell DNA. Though colibactin has been found to alkylate DNA, the molecular basis for cross-link formation is unclear. Here, we report that the colibactin biosynthetic enzyme ClbL is an amide bond-forming enzyme that links aminoketone and β-keto thioester substrates *in vitro* and *in vivo*. The substrate specificity of ClbL strongly supports a role for this enzyme in terminating the colibactin NRPS-PKS assembly line. This transformation would incorporate two electrophilic cyclopropane warheads into the final natural product scaffold. Overall, this work provides a biosynthetic explanation for colibactin’s DNA crosslinking activity and paves the way for further study of its chemical structure.

## INTRODUCTION

Colibactin is a genotoxin made by human gut commensal and extraintestinal pathogenic *Escherichia coli* strains and other Proteobacteria.^1^ Colibactin-producing *E. coli* (*pks*^+^ *E. coli*) cause DNA double strand breaks,^1–2^ affect progression of colitis-associated colorectal cancer (CRC) in mouse models,^3–4^ and are more frequently detected in patients with colorectal cancer.^5–7^ Recent studies have indicated colibactin’s genotoxicity likely arises from a direct interaction with DNA, as we and others have reported the accumulation of interstrand cross-links in human cell lines incubated with *pks*^+^ *E. coli*.^8–9^ However, the underlying chemical and enzymatic basis for this activity remains unclear.

Colibactin is produced by a 54-kb nonribosomal peptide synthetase-polyketide synthase (NRPS-PKS) hybrid assembly line (the *pks* island),^1^ and its biosynthesis involves a self-protection mechanism in which an *N*-acyl-D-asparagine scaffold (prodrug motif) is initially assembled and elaborated to produce an inactive metabolite (precolibactin) (**Figure 1A**).^10–12^ Precolibactin is cleaved by the periplasmic peptidase ClbP to release the active colibactin genotoxin. This active species has not yet been identified, isolated, or structurally characterized, presumably due to its instability. We and others have isolated and structurally characterized several stable candidate precolibactins (**1-5**) from *pks*^+^ *E. coli* strains lacking ClbP (**Figure 1B**).^13–20^ Notably, metabolites **2-5** contain a cyclopropane ring found in some DNA-alkylating agents. Formation of an α,β-unsaturated imine after cleaving the prodrug motif in precolibactin enhances the electrophilicity of the cyclopropane toward DNA bases, which led to the identification of colibactin-DNA adducts.^9, 21^ However, it is unclear how this structural motif could lead to generation of an interstrand cross-link.

**Figure 1.**
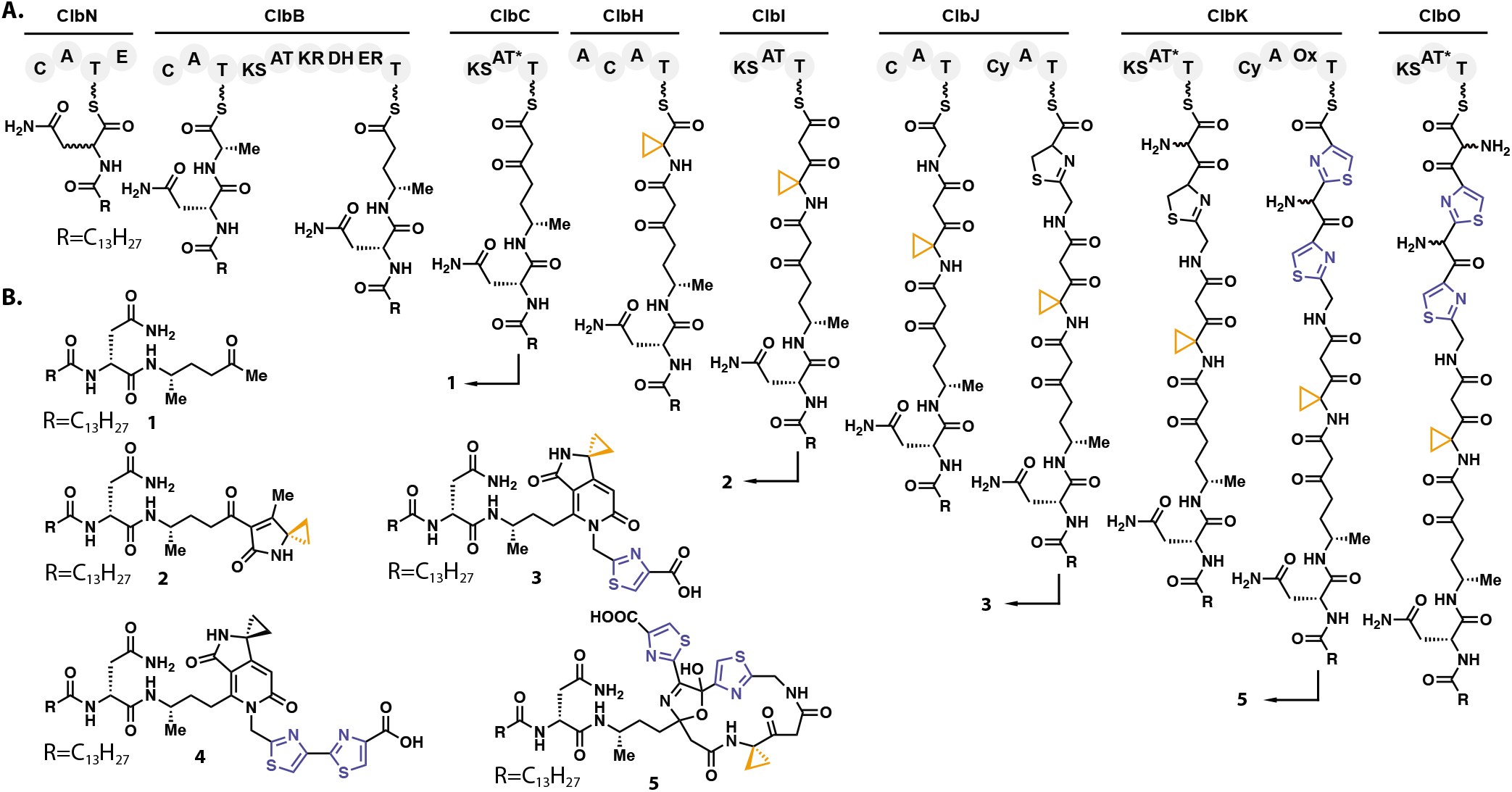
The *pks* enzymatic assembly line and the biosynthesis of characterized candidate precolibactins. (**A**) Hypothetical biosynthetic pathway for candidate precolibactins **1-3** and **5**. C, condensation; A, adenylation; T, thiolation; E, epimerization; KS, ketosynthase; AT, acyltransferase; KR, ketoreductase; DH, dehydratase; ER, enoyl reductase; AT*, atypical acyltransferase; Cy, cyclization; Ox, oxidase. Cyclopropane rings are highlighted in orange. Thiazoles are highlighted in blue. (**B**) Chemical structures of isolated and characterized candidate precolibactins **1-5**.

To address this question, we focused on understanding previously uncharacterized components of the colibactin biosynthetic enzymes. Efforts to elucidate the biosynthetic origins of isolated candidate precolibactins have led to putative functional assignments for all of the enzymes in the biosynthetic pathway except for the putative amidase ClbL. This enzyme is essential for the genotoxicity of *pks*^+^ *E. coli*^1^ but is not required for biosynthesis of any isolated candidate precolibactins reported to date. A recent finding that *pks*^+^ *E. coli* strains with inactive ClbL were incapable of cross-linking DNA suggested that ClbL participates in producing the active cross-linking agent.^22^ However, the molecular basis for ClbL’s contribution to colibactin’s cross-linking activity is unknown.

Here, we elucidate the function of ClbL in colibactin biosynthesis. By identifying, isolating, structurally characterizing a ClbL-dependent candidate precolibactin from bacterial cultures, and reconstituting its biosynthesis *in vitro*, we show that ClbL is an amide bond-forming enzyme. ClbL couples β-keto thioesters and aminoketones, structural motifs known to be generated by the colibactin assembly line. Based on this observation, we hypothesize that ClbL assembles a pseudo-dimeric precolibactin precursor possessing two electrophilic cyclopropane warheads which upon ClbP activation can crosslink DNA. This proposal is supported by the detection of two new colibactin-derived DNA adducts in *pks*^+^ *E. coli* treated DNA samples. Overall, this work reveals the activity of ClbL and provides a molecular explanation for colibactin’s DNA cross-linking activity that incorporates all essential colibactin biosynthetic enzymes.

## Results

ClbL belongs to the amidase signature (AS) family of enzymes, which contain a Ser-*cis*Ser-Lys catalytic triad (**Figure S1**).^23^ These enzymes catalyze a wide variety of hydrolytic reactions, including the breakdown of the neurotransmitter anandamide by fatty acid amide hydrolase (FAAH) in humans and hydrolysis of glutamine by Glu-tRNA^Gln^ amidotransferase subunit A (GatA) in bacteria.^24–25^ Phylogenetic analysis of representative amidases indicated that ClbL is part of a distinct clade, potentially implying functional differences (**Figure S2**). Alignment of ClbL with structurally characterized amidases revealed the three conserved, essential catalytic residues (K80, S155, and S179) (**Figure S1**). To test whether these active site amino acids are essential for ClbL function and genotoxicity, plasmids expressing versions of ClbL with these residues individually mutated to alanine were constructed and used to complement a strain of *pks*^+^ *E. coli* lacking *clbL* (BAC*pksΔclbL*). These mutant strains failed to cause cell cycle arrest in HeLa cells (**Figure 2A**), suggesting that these three residues are crucial for the function of ClbL and that ClbL’s catalytic activity is critical for genotoxicity.

**Figure 2.**
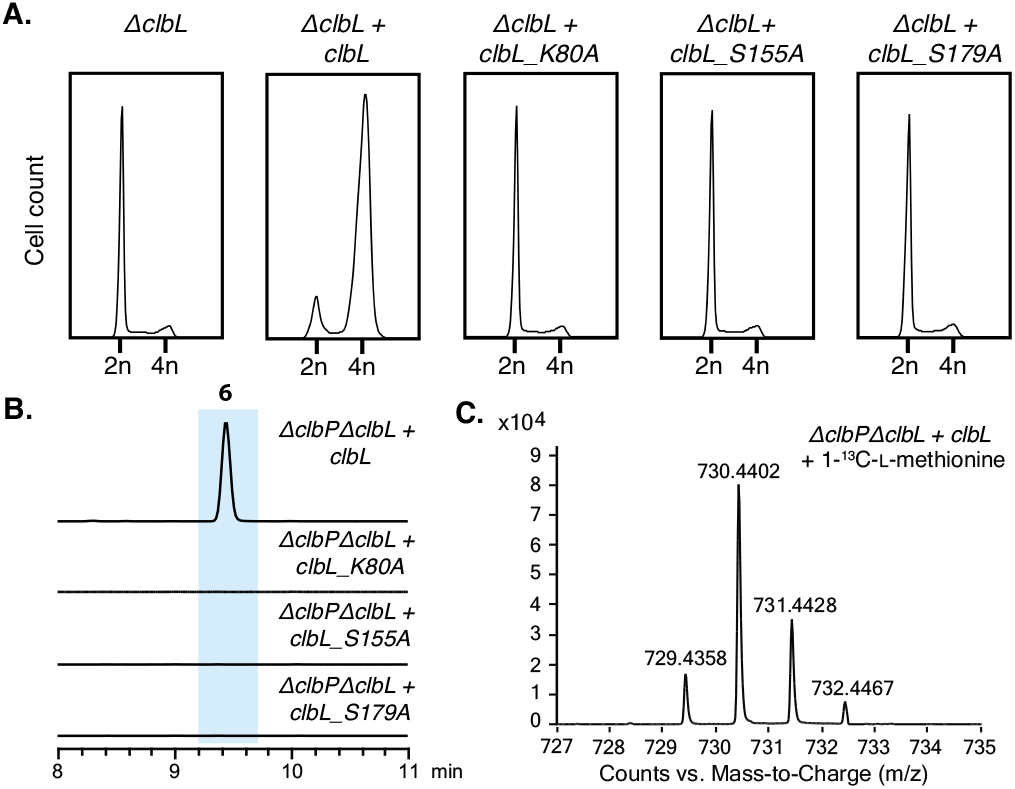
Discovery of a ClbL-dependent candidate precolibactin. (**A**) Cell cycle analysis of HeLa cells infected with wild-type or mutant ClbL expressing *E. coli* strains. (**B**) Extracted ion chromatograms (EICs) of metabolite **6** (*m*/*z* 729.4334) in the extracts of *E. coli* expressing wild-type or mutant ClbL. (**C**) LC-MS analysis of feeding of 1-^13^C-L-methionine to ClbL expressing *E. coli* strains.

To identify candidate precolibactins linked to ClbL, we overexpressed wild-type ClbL or the ClbL active site mutants in *E. coli* DH10B BAC*pksΔclbP/ΔclbL*. Methanol extracts from these cultured strains were compared to identify mass features affected by the activity of ClbL. As amidases are typically amide bond-cleaving enzymes, we expected the substrate for ClbL to accumulate in the mutant strains. Surprisingly, we were unable to detect any up-regulated metabolites. Consistent with the previous findings,^16^ ClbL mutant strains produced detectable levels of **2** and other known candidate precolibactins (**Figure S3**), confirming that these metabolites are not ClbL-dependent. We also observed a novel metabolite (**6**, *m*/*z* 729.4334) in strains expressing wild-type ClbL that was completely abolished in strains expressing ClbL mutants (**Figure 2B, Table S4**). To confirm metabolite **6** was *pks*-associated, we fed wild-type ClbL expressing strains with L-[1-^13^C]Met, the amino acid precursor of the cyclopropane ring.^15,16^ Observation of a +1 mass shift demonstrated that methionine is incorporated into **6**. Moreover, MS/MS fragmentation analysis indicated that **6** contains the prodrug scaffold (*m*/*z* 324.2437)^18^ and a colibactin-characteristic fragment ion (*m*/*z* 231.1119)^17, 26^ (**Figure S4**). These results suggested that metabolite **6** is a novel ClbL-dependent candidate precolibactin.

We then isolated and structurally characterized this new metabolite, obtaining approximately 0.7 mg of purified 6 from 140 L of an *E. coli* DH10B BAC*pksΔclbP* strain overexpressing ClbL. The high-resolution mass of this compound ([M+H]^+^ = 729.4334) yielded a molecular formula of C_41_H_56_N_6_O_6_. One-dimensional (1D) and two-dimensional (2D) NMR along with MS/MS fragmentation analysis allowed us to elucidate its chemical structure (**Figure 3A, Figure S5-S8, Table S5**).

**Figure 3.**
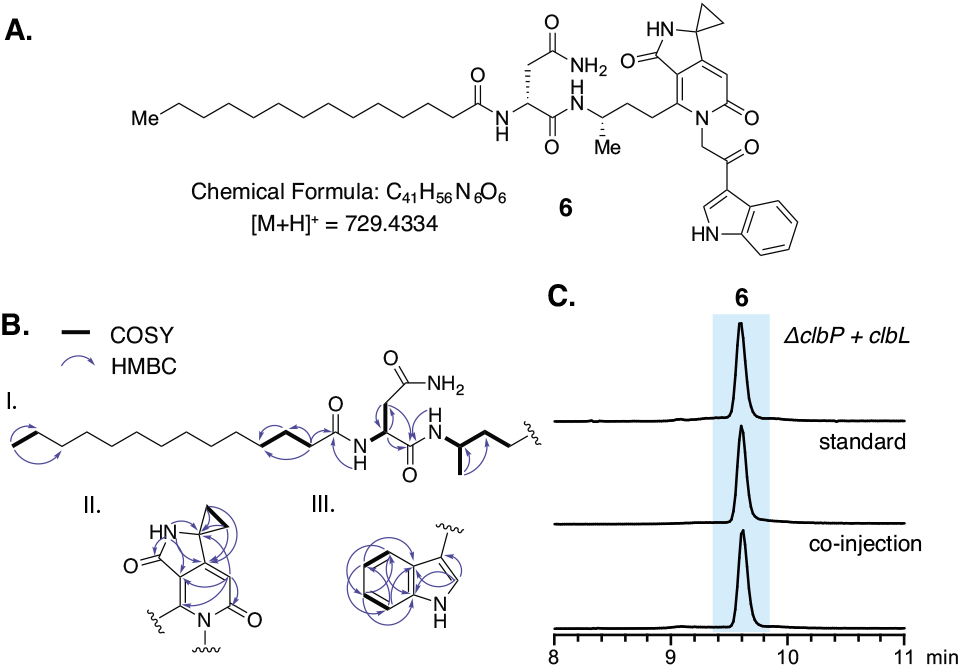
Structure determination of candidate precolibactin 6. (**A**) Chemical structure of metabolite **6**. (**B**) Key COSY and HMBC correlations that support this assignment. (**C**) EICs of metabolite **6** (*m*/*z* 729.4334) from the extracts of bacterial cultures, synthetic standard, and co-injection of synthetic standard and bacterial culture extracts.

Similarities between the NMR spectra of known precolibactins and metabolite **6** enabled us to assign fragments I and II easily (**Figure 3B**). Unexpectedly, COSY and HMBC correlations indicated the presence of an indole ring (**Figure 3B**). We were unable to assign the precise locations of a methylene and carbonyl within **6** using NMR, but MS/MS fragmentation analysis revealed a loss of indole (*m*/*z* 117.0607) from the parent ion, suggesting that the ketone is directly attached to the indole ring (**Figure S9**). To further support this structural assignment, we chemically synthesized a standard of the proposed metabolite **6** structure. This standard possessed the same exact mass, MS/MS fragmentation pattern, retention time, ^1^H spectrum, and UV spectrum as **6** isolated from bacterial strains (**Figure 3C, Figure S10-S12**), confirming our structural assignment.

The chemical structure of **6** guided subsequent efforts to elucidate the function of ClbL. Metabolite **6** does not contain either a free amine or a carboxylic acid, which are the functional groups liberated by the activity of characterized amidases. Based on the known steps of colibactin biosynthesis, we envisioned two possible pathways for the formation of **6** (**Figure 4A**). In pathway A, the intermediate generated by ClbJ-NPRS1^17,18^ could be intercepted and offloaded via nucleophilic attack by the known *E. coli* metabolite indole.^27^ In pathway B, 2-amino-1-(1*H*-indol-3-yl)ethan-1-one (**7**) could act as a nucleophile to offload the β-keto thioester intermediate generated by the upstream biosynthetic enzyme ClbI.^28^

**Figure 4.**
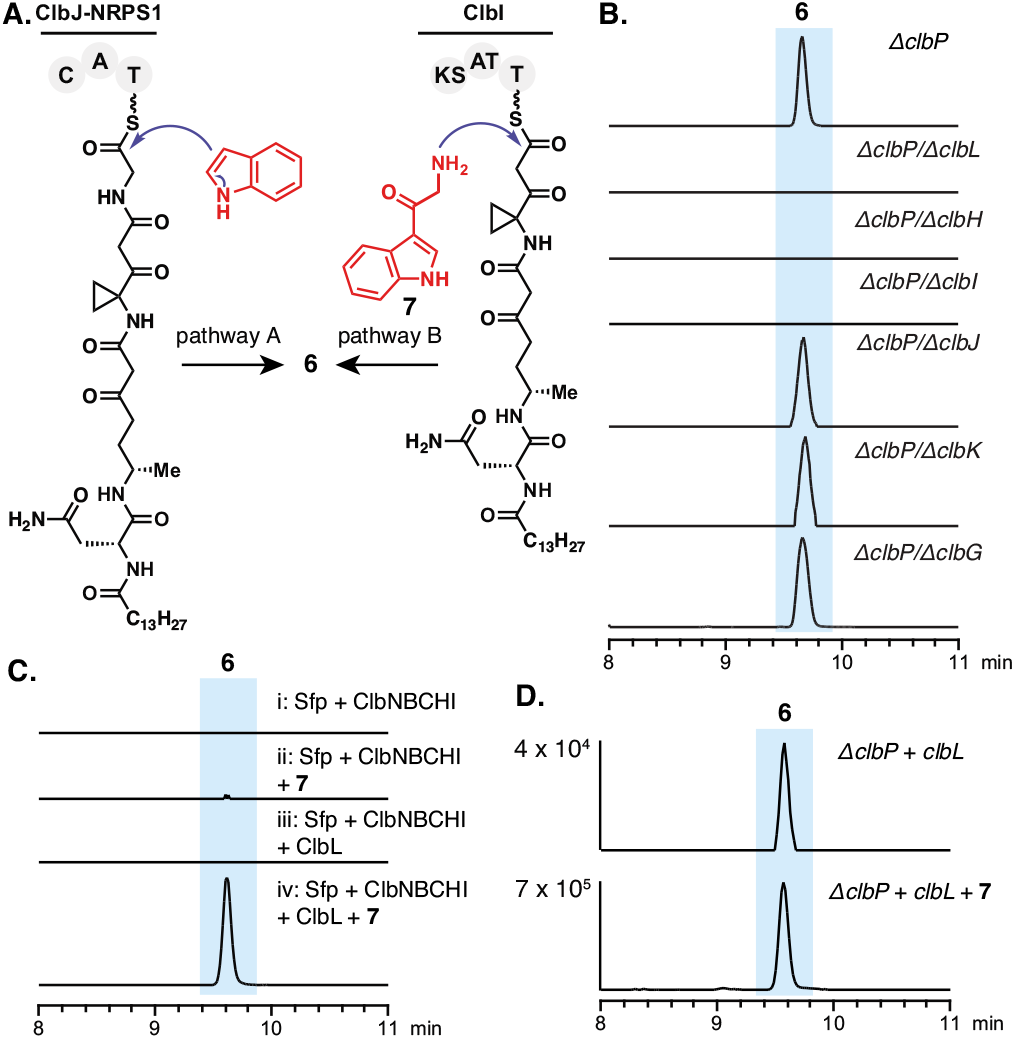
Investigating the formation of candidate precolibactin 6. (**A**) Two proposed pathways could generate **6**. (**B**) LC-MS analysis of extracts from double mutants of *E. coli* DH10B BAC*pks*. (**C**) LC-MS analysis of *in vitro* reconstitution assays. (**D**) LC-MS analysis of cell pellet extracts from *E. coli* DH10B BAC*pksΔclbP* + pTrc-*clbL* supplemented with **7**. Panels **B-D** show the EICs of **6** (*m*/*z* 729.4334).

To test the involvement of pathway A, we made multiple double mutants in *E. coli* DH10B BAC*pksΔclbP* (**Figure 4B**). Detection of **6** in the absence of ClbJ and ClbP indicated that ClbJ was not required for its biosynthesis. To confirm this result, we fed [1,2-^13^C_2_]Gly to a glycine auxotrophic BAC*pksΔclbP* strain. We observed a +2 mass shift in the known glycine-derived precolibactin **3**, but not in **6** (**Figure S13**), confirming that pathway A is not the source of **6**.

We next explored the involvement of pathway B by reconstituting the biosynthesis of metabolite **6** *in vitro*. We cloned, overexpressed, and purified C-terminal His6-tagged ClbL (**Figure S14**). An assay mixture containing Sfp, ClbN, ClbB, ClbC, ClbH, ClbI, ClbL, myristoyl-CoA, malonyl-CoA, L-Asn, L-Ala, NADPH, ATP, SAM, and **7** generated **6** (**Figure 4C**, trace iv). To further confirm the involvement of **7** in this process, **7** was fed to an *E. coli* DH10B BAC*pksΔclbP* strain overexpressing ClbL. An approximately 20-fold increase of the production of **6** was observed (ion count of 4 x 10^4^ vs. 7 x 10^5^, **Figure 4D**), indicating pathway B is operative *in vivo*. Notably, these results suggested that ClbL catalyzes amide bond formation rather than amide bond hydrolysis as was originally predicted by bioinformatics.

To characterize the activity of ClbL *in vitro*, we synthesized ethyl thioester **8** as a surrogate for the ClbI-tethered biosynthetic intermediate (**Figure 1A and 5A**). Incubating ClbL with **7** and **8** (**Figure 5A**) revealed a single new product peak accumulating during an HPLC time course experiment (**Figure 5B**). We confirmed this product was amide **9** by comparison to a synthetic standard (**Figure S15**).

**Figure 5.**
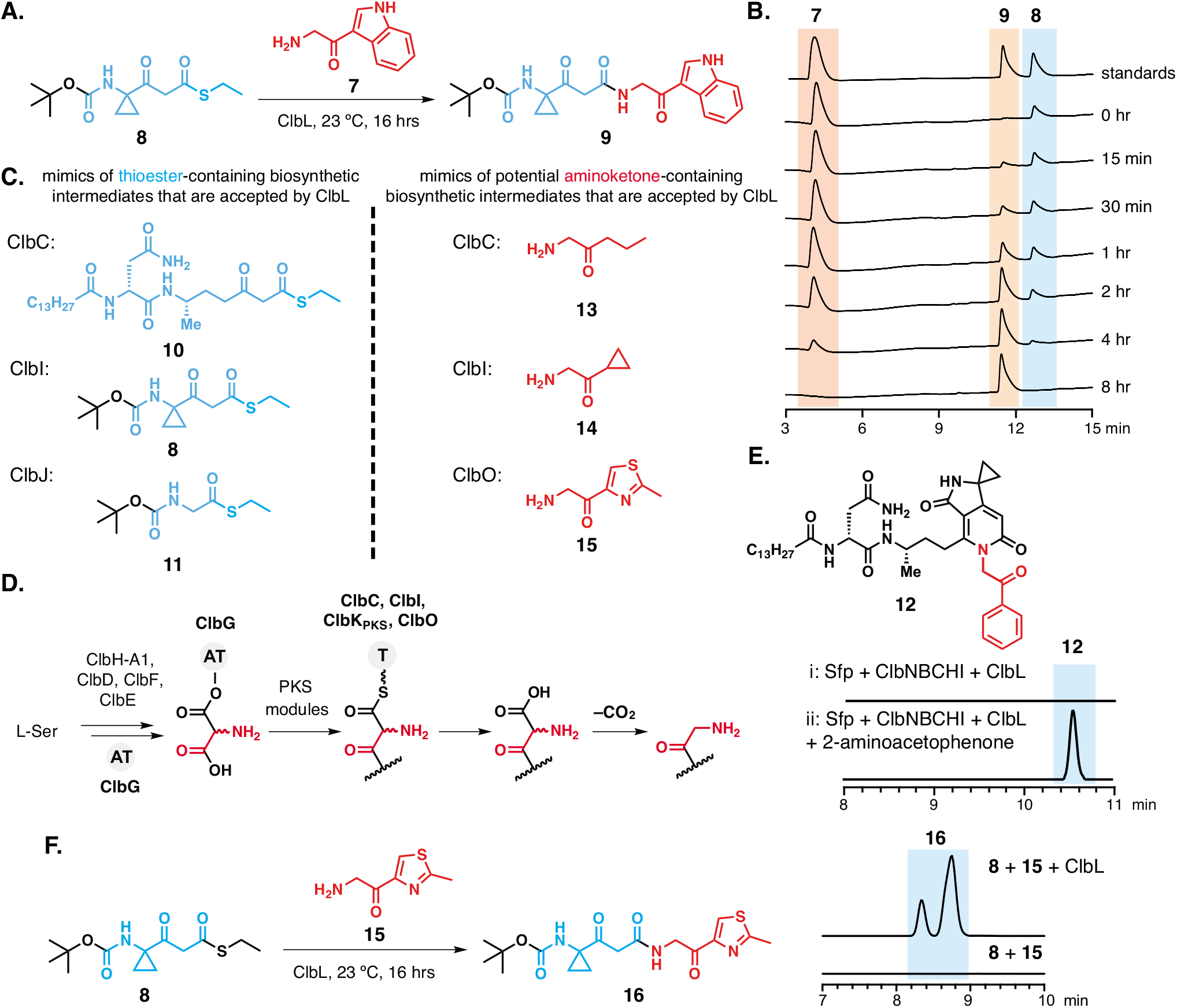
*In vitro* characterization of ClbL reveals its role in amide formation. (**A**) Reaction scheme of an *in vitro* assay to form **9**. (**B**) HPLC time course for the formation of **9** by ClbL (monitored at 250 nm). (**C**) Thioesters and α-aminoketones accepted by ClbL. (**D**) The *pks* assembly line may produce α-aminoketone nucleophiles. (**E**) EICs of **12** (*m*/*z* 690.4225) formed in an *in vitro* reconstitution assay. (**F**) ClbL catalyzes the formation of **16** *in vitro*. The two peaks likely correspond to two tautomers of the β-ketoamide product.

To evaluate ClbL’s preference for potential thioester substrates, we designed and synthesized ethyl thioester mimics of different assembly line-tethered colibactin biosynthetic intermediates (**Figure S16**). ClbL and **7** were incubated with these mimics individually, among which thioester **10** (a mimic for the ClbC-bound intermediate) and **11** (a mimic for the ClbJ-bound intermediate) could be recognized by ClbL (**Figure 5C, Figure S17**). To assess ClbL’s ability to discriminate among these substrates, we conducted a competition experiment. Mixing **8** and **11** in equal molar amounts with ClbL and **7** resulted in accumulation of more of the product arising from **8** compared to the product arising from **11** (**Figure S17**). This suggests ClbL preferentially recognizes the ClbI-bound intermediates. Furthermore, reexamination of our earlier *in vitro* reconstitution of the biosynthesis of **6** did not reveal the product of amide bond formation between **7** and the ClbC-tethered thioester intermediate. Together, these data indicate that ClbL preferentially uses the ClbI-bound thioester as an electrophile for amide bond formation.

We next sought to investigate the relevance of indole-containing nucleophile **7** to colibactin biosynthesis. This metabolite has only previously been found in termitomycamide B, which was isolated from the giant mushroom *Termitomyces titanicus* (**Figure S18**).^29^ Intriguingly, **7** contains an α-aminoketone, a structural motif that could originate from decarboxylation of the essential colibactin building block aminomalonate (**Figure 5D**). To examine the involvement of aminomalonyl-ACP biosynthetic enzymes in the production of **7**, we used an *E. coli* DH10B BAC*pksΔclbPΔclbG* strain, in which the transfer of aminomalonate to assembly line enzymes is abolished. Production of **6** by this strain indicates this building block is not required (**Figure 4B**). We further showed that **7** does not originate from any of the *pks* island enzymes as it was detected in *E. coli* DH10B (**Figure S19**). **7** was also absent from growth medium, indicating it is produced in *E. coli* endogenously. Therefore, we questioned whether the incorporation of **7** was relevant for the biosynthesis of the active colibactin genotoxin. Feeding **7** to *pks*^+^ *E. coli* during infection of host cells did not boost genotoxicity (**Figure S20**), suggesting that **7** is likely not a relevant substrate for ClbL in colibactin biosynthesis. We therefore hypothesize that metabolite **6** arises from the promiscuous activity of ClbL and is not an on-pathway intermediate.

In considering potential alternative substrates for ClbL, we recognized that the α-aminoketone motif in 7 may be generated by colibactin biosynthetic enzymes and incorporated into ClbL’s native substrate. Indeed, assays with alternative amine substrates showed that the α-aminoketone of **7** is crucial for recognition by ClbL. No products were observed when tryptamine replaced **7** in assay mixture for *in vitro* reconstitution of the biosynthesis of **6**. In contrast, we identified a new product **12** (*m*/*z* 690.4225) in assay mixtures containing 2-aminoacetophenone in place of **7** (**Figure 5E**). This result suggested the α-aminoketone and not the indole ring that is important for substrate recognition by ClbL.

To identify the true nucleophilic coupling partner for ClbL, we examined its reactivity toward potential α-aminoketone substrates derived from the colibactin assembly line. We previously identified four PKS enzymes in the assembly line (ClbC, ClbI, ClbK, and ClbO) that could accept aminomalonate *in vitro*^18^ Isolation of candidate precolibactin **5** also showed that the PKS module of ClbK accepts this building block *in vivo*, generating an α-aminoketone that could be further oxidized to an imine.^19^ Once offloaded from PKS modules, terminal aminomalonates should readily undergo decarboxylation to give rise to α-aminoketones resembling **7** (**Figure 5D**). To examine whether ClbL can use assembly line derived α-aminoketones for amide bond formation, we obtained mimics of these potential substrates (**13-15**) and incubated them with ClbL and preferred thioester **8** (**Figure S21**). While ClbL can use all three mimics, it preferentially recognizes **15**, which mimics the α-aminoketone derived from ClbO, the final module of the colibactin assembly line (**Figure 5C** and **5F**, **Figure S22**). Together, these results suggest that ClbL’s role in colibactin biosynthesis is to form an amide bond between a ClbI-bound thioester and the α-aminoketone derived from ClbO (**Figure 6**).

**Figure 6.**
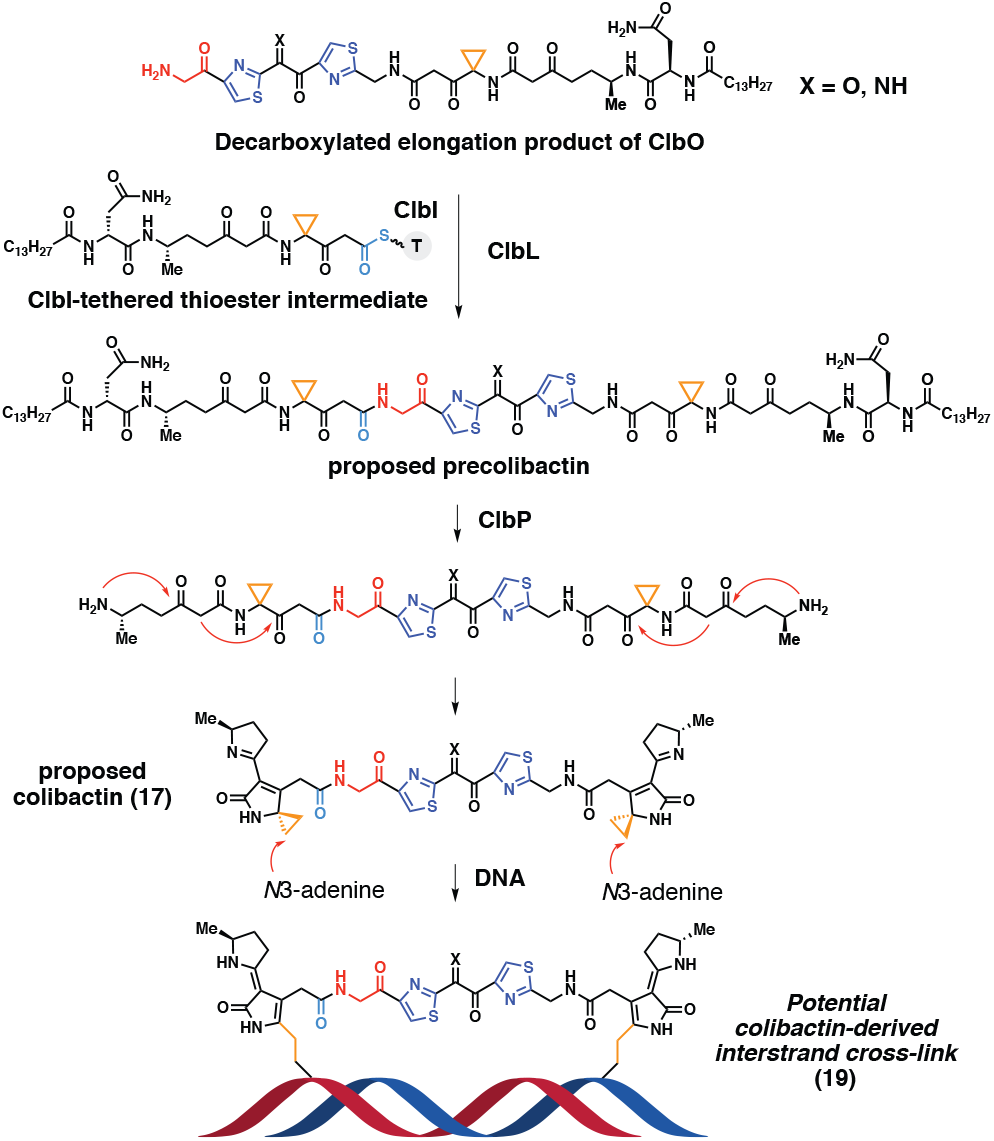
A model for ClbL-catalyzed formation of the active colibactin genotoxin). Key structural features colored as in Figures 1, 4, and 5.

This proposed assembly line logic has important implications for colibactin’s biological activity. Amide bond formation by ClbL could link the precursors of two electrophilic cyclopropane warheads together. Cleavage of the prodrug motif by ClbP would generate a species capable of generating interstrand cross-links (**17**) (**Figure 6**). This metabolite could be the active colibactin genotoxin. This scenario is consistent with previous observations that *pks*^+^ *E. coli* promote interstrand cross-link formation in linearized plasmid DNA and human cell lines,^8,9^ as well as a recent finding that a synthetic ‘colibactin mimic’ containing two cyclopropane rings cross-links DNA *in vitro*, while a monomer cannot.^30^ This proposal is consistent with our current understanding of colibactin-mediated DNA damage. We previously identified and characterized a pair of diasteromeric colibactin-derived DNA adducts (**18**) that likely arise from degradation of a larger lesion.^9^ Depurination of the proposed colibactin-DNA cross-link **19** would give adduct **20**, which could undergo oxidative cleavage to give monoadducts **21** and **18** (**Figure 7A**).

**Figure 7.**
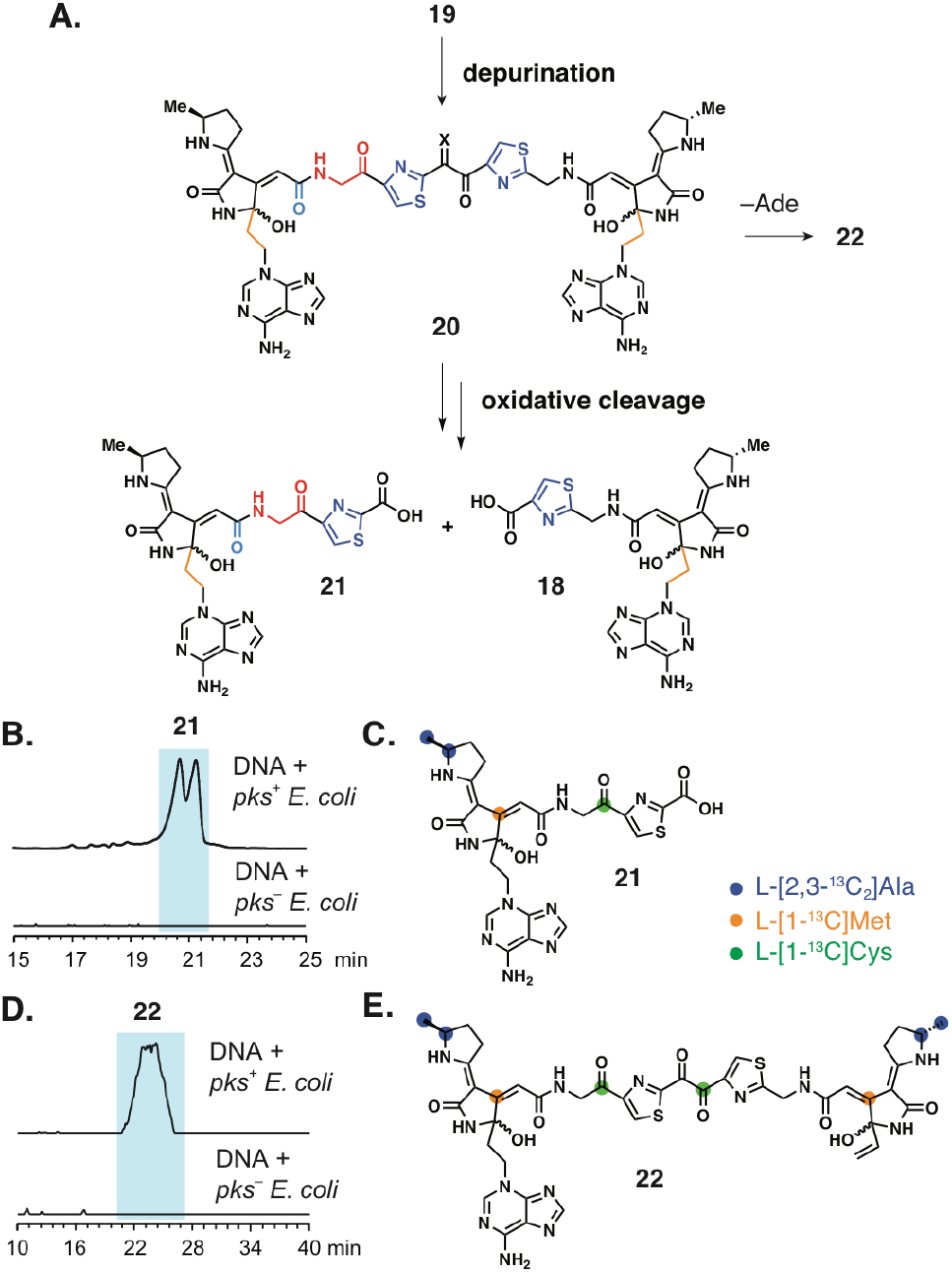
Detection of two new colibactin-derived DNA adducts. (**A**) Predicted structures of colibactin-derived DNA adduct **20** and its decomposition products **21** and **18**. (**B**) EICs of **21** (*m*/*z* 568.1721) in DNA samples treated with *pks*^+^ *E. coli*. The two peaks likely arise from multiple diastereomers. (**C**) Adduct **21** incorporates one alanine, one methionine, and one cysteine. (**D**) EICs of **22** (*m*/*z* 938.2855) in *pks*^+^ *E. coli* treated DNA samples. (**E**) Adduct **22** incorporates two alanines, two methionines, and two cysteines.

To test this proposal, we used untargeted DNA adductomics^9^ to identify two colibactin-derived DNA adducts in linearized plasmid DNA treated with *pks*^+^ *E. coli*. The first DNA adduct (*m*/*z* 568.1721) had an exact mass corresponding to proposed decomposition product **21** (**Figure 7B**). MS/MS fragmentation analysis and isotope labeling studies also supported this proposed structure (**Figure 7C, Figure S23-25**). While the cross-linked adduct **20** was not detected possibly due to its low abundance and instability, we were able to find a large DNA adduct **22** (*m*/*z* 938.2855), which could derive from diketone **20** (**Figure 7D**). To confirm this adduct is *pks*-associated, we supplemented assay mixtures with either L-[1-^13^C]Met or L-[1-^13^C]Cys. The observation of +2 mass shifts in **22** suggested that it derives from two methionines and two cysteines (**Figure 7E, Figure S26**), supporting our proposal that ClbL generates a precolibactin containing two electrophilic warheads and two thiazoles. Further characterization of colibactin-derived adducts will be necessary to clarify the identity of the active cross-linking agent.

## Conclusions

ClbL belongs to the AS enzyme family, which participates in numerous biological processes, including the biosynthesis of the natural product actinonin in *Streptomyces* sp. ATCC 14903.^31^ In addition to catalyzing amide bond hydrolysis, some amidases exhibit acyltransferase activity. For example, the amidase from *Rhodococcus sp*. R312 transfers the acyl group of an amide to hydroxylamine at a 33-fold faster rate than the hydrolysis of the same amide, an activity that could be explained by the stronger nucleophilicity of hydroxylamine toward the acyl-enzyme intermediate compared to water.^32^ Our discovery of ClbL’s activity expands the known roles of amidases to include interfacing with enzymatic assembly lines. It is intriguing that ClbL preferentially recognizes α-aminoketones over the corresponding primary amines. Future crystallographic studies may shed further light on this specificity.

ClbL’s *in vitro* activity leads us to propose that this enzyme links two intermediates generated by the colibactin NRPS-PKS assembly line, a ClbI-bound thioester and a ClbO-derived α-aminoketone (**Figure 6**). This transformation likely plays a critical role in generating a DNA cross-linking agent that could represent the active colibactin genotoxin. This proposal is supported by ClbL’s essential role in genotoxicity and cross-link formation, as well as our detection of adduct **22** in linearized plasmid DNA treated with *pks*^+^ *E. coli*. We cannot exclude the possibility that ClbL accepts additional substrates. For example, we were unable to test whether ClbL could use a mimic of the putative ClbO-bound aminomalonate thioester intermediate as an electrophile due to its intrinsic instability. However, precedented colibactin biosynthetic logic and its DNA interstrand cross-linking activity allow us to establish two minimal criteria for the structure of colibactin: 1) it contains two electrophilic cyclopropane rings and 2) its construction requires all of the essential colibactin biosynthetic enzymes. Application of these criteria to the potential products of ClbL greatly narrows the potential structures for the active colibactin genotoxin(s) (**Figure S27**). These criteria also help in assessing the biological relevance of other reported candidate precolibactins. For example, Qian and Zhang recently disclosed a candidate precolibactin they proposed is the precursor to colibactin (**Figure S28**).^20^ Though the corresponding ClbP cleavage product appeared to damage DNA *in vitro* and in cells in a Cu^2+^-dependent manner, it did not form cross-links, consistent with the presence of only one cyclopropane. Moreover, the production of this compound does not require ClbL, making it unlikely to be a major contributor to genotoxicity.

In summary, these studies provide a biochemical rationale for the formation of interstrand DNA cross-links by the gut bacterial genotoxin colibactin. As such cross-links represent the most toxic form of DNA damage in human cells, it is possible this activity could explain the effects of *pks*^+^ *E. coli* on tumor development. Future efforts to isolate and characterize the final pre-colibactin, colibactin, and colibactin-DNA adducts will provide us with the additional molecular information needed to decipher how colibactin influences CRC development in patients.

## ACKNOWLEDGMENT

We thank Alex Sieg for help with cloning, Lihan Zhang for help with NMR experiments, and Matthew Volpe for comments on the manuscript. We acknowledge financial support from National Institutes of Health (R01CA208834-02) (E. P. B). Salary support for P.W.V. was provided by the U.S. National Institutes of Health and National Cancer Institute [Grant R50-CA211256]. Mass spectrometry for DNA adduct analysis was carried out in the Analytical Biochemistry Shared Resource of the Masonic Cancer Center, supported in part by the U.S. National Institutes of Health and National Cancer Institute [Cancer Center Support Grant CA-77598]. M.R.W. acknowledges support from the American Cancer Society-New England Division Postdoctoral Fellowship PF-16-122-01-CDD.

