## Supplementary Material for "The reactivity of an unusual amidase may explain colibactin’s DNA cross-linking activity"

### Table of Contents

**1. Supplementary methods** ........ **S2**
**2. Supplementary tables** ........ **S11**
**3. Supplementary figures** ........ **S15**
**4. Synthetic procedures** ........ **S39**
**5. NMR spectra** ........ **S45**
**6. Supplementary references** ........ **S59**

### 1. Supplementary methods

**General materials and methods.** All chemicals and solvents were purchased from Sigma Aldrich (St. Louis, MO) except where noted. Oligonucleotide primers were synthesized by Sigma Aldrich (St. Louis, MO). DNA polymerases, Gibson Assembly Master Mix, restriction enzymes, T4 ligase, and electrocompetent *E. coli* NEB® 10-beta cells were purchased from New England BioLabs (Ipswich, MA). Thermocycling was carried out in a C1000 Gradient Cycler (Bio-Rad Laboratories, Hercules, CA). Gel extraction of DNA fragments and restriction endonuclease clean up were performed using a Zymoclean Gel DNA Recovery Kit from Zymo Research (Irvine, CA) or an Illustra GFX PCR DNA and Gel Band Purification Kit from GE Healthcare (Chicago, IL). DNA concentrations were determined using a NanoDrop 2000 UV-Vis Spectrophotometer (Thermo Fisher Scientific, Waltham, MA). Luria-Bertani Lennox (LB) medium was purchased from Alfa Aesar (Tewksbury, MA). Recombinant plasmid DNA was purified with a QIAprep Spin Miniprep Kit from Qiagen (Hilden, Germany). DNA sequencing was performed by Eton Bioscience (San Diego, CA).

Strains of DH10B *E. coli* harboring pBeloBAC11-*pks* or pBeloBAC11-*pks*Δ*clbP* were obtained from the Bonnet lab, Laboratoire de Bactériologie Clinique, Centre Hospitalier de Clermont-Ferrand, Clermont-Ferrand F-63003, France. The glycine auxotrophic strain (*E. coli* JW2535-1) was obtained from the Coli Genetic Stock Center (CGSC, Yale University, New Haven, CT). Isotope-labeled amino acids fed to bacterial cultures were purchased from Cambridge Isotope Laboratories (Tewksbury, MA). Optical densities of *E. coli* cultures were determined with a DU 730 Life Sciences UV-Vis spectrophotometer from Beckman Coulter (Brea, CA) by measuring absorbance at 600 nm.

HeLa cells were obtained from the American Type Culture Collection (ATCC, Manassas, VA). Dulbecco's phosphate buffered saline (DPBS), Dulbecco's modified eagle medium (DMEM), 100× Antibiotic-Antimycotic, fetal bovine serum (FBS), 1M HEPES, and RNase A were purchased from Thermo Fisher Scientific.

Proton nuclear magnetic resonance ($^1$H NMR) and two-dimensional (2D NMR) spectra of candidate precolibactin **6** were obtained on a Bruker AVANCE II 600 MHz spectrometer equipped with a TXO cryoprobe (Bruker-BioSpin Corporation, Billerica, MA) in the East Quad NMR facility at Harvard Medical School (Boston, MA). Other NMR spectra were collected in the Magnetic Resonance Laboratory in the Department of Chemistry and Chemical Biology at Harvard University (Cambridge, MA). Chemical shifts (δ) are reported in parts per million (ppm) downfield from tetramethylsilane using the solvent resonance as an internal standard for $^1$H ($CDCl_3$ = 7.26 ppm, DMSO-$d_6$ = 2.50 ppm) and $^{13}$C ($CDCl_3$ = 77.25 ppm, DMSO-$d_6$ = 39.52 ppm). Data are reported as follows: chemical shift, multiplicity (br = broad, s = singlet, d = doublet, t = triplet, q = quartet, m = multiplet), coupling constants in Hertz (Hz), and integration. NMR solvents were purchased from Cambridge Isotope Laboratories (Tewksbury, MA). NMR spectra were visualized using MestReNova, version 12.0.1-20560 (Mestrelab Research, Escondido, CA). Optical rotation data were obtained using a 1 mL cell with a 0.5 dm path length on a Jasco P-2000 polarimeter equipped with a sodium (589 nm, D) lamp.

During protein purification, affinity chromatography and SDS-PAGE analysis were conducted using HisPur nickel-nitrilotriacetic acid-agarose (Ni-NTA) resin (Thermo Fisher Scientific) and 4-15% Mini-PROTEAN TGX Precast SDS-PAGE gels with 2× Laemmli samples buffer (Bio-Rad). Protein concentrations were determined by measuring absorbance at 280 nm with a NanoDrop 2000 UV-Vis Spectrophotometer (Thermo Fisher Scientific).

Analytical, semi-preparative, and preparative HPLC was performed on a Dionex UltiMate 3000 instrument (Thermo Fisher Scientific). LC/MS and LC/MS/MS analyses were carried out on an Agilent 6530 Accurate-Mass quad time-of-flight mass spectrometer system equipped with an electrospray ionization source (Agilent Technologies, Palo Alto, CA). Liquid chromatography was performed on an Agilent 1260 Infinity HPLC using a Hypersil Gold™ aQ C18 polar endcapped column (50 × 3.0 mm, 3 µm particle size, Thermo Fisher Scientific). The injection volume was 10 µL and the flow rate was 0.35 mL/min using 0.1% formic acid in acetonitrile as mobile phase A and 0.1% formic acid in water as mobile phase B.

**Multiple sequence alignment of ClbL.** Amino acid sequences of ClbL and seven structurally characterized amidases were obtained from the NCBI public database. Multiple sequence alignments were generated using ClustalW in MEGA7[1] and visualized in Jalview.[2]

**Phylogenetic analysis of ClbL.** Amino acid sequences of amidase signature family enzymes from different species were obtained from the NCBI public database and aligned by ClustalW in MEGA7.[1] The phylogenetic tree was constructed in MEGA7 using the Maximum Likelihood method with LG model in a bootstrap test of 1,000 replicates and visualized in iTOL (https://itol.embl.de).[3]

**Inactivation of *clbL*, *clbH*, *clbI*, *clbJ*, and *clbK* using λ Red recombinase-mediated gene disruption.** Inactivation of clbG was carried out as reported.[4] The target genes *clbL*, *clbH*, *clbI*, *clbJ*, and *clbK* were disrupted by the PCR-targeting λ Red recombinase-mediated gene disruption system.[5] The disruption cassette, which included the apramycin resistance gene *aac(3)IV* flanked by 39 bp arms homologous to the target gene, was generated by PCR amplification using primers shown in **Supplementary Table 1**. PCR reactions (20 µL) contained 10 µL of Q5 High Fidelity 2× Master Mix, 1 ng of plasmid DNA template pIJ773, and 500 pmoles of each primer. Thermocycling was carried out in a C1000 gradient cycler using the conditions shown in **Supplementary Table 3**. The disruption cassettes were purified using a Zymoclean Gel DNA Recovery Kit.

For λ Red recombinase-mediated gene disruption, the purified disruption cassette was electroporated into *E. coli* BW25113 harboring the λ Red recombinase expression plasmid pkD46 and either BAC*pks* or BAC*pks*Δ*clbP*. Mutants were selected on LB plates containing 50 µg/mL apramycin and 25 µg/mL chloramphenicol. The mutant BACs were isolated from these colonies and gene disruption was verified by PCR. DH10B *E. coli* were transformed by electroporation with these mutant BACs and selected on LB plates containing 50 µg/mL apramycin and 25 µg/mL chloramphenicol.

**Design and introduction of *clbL, clbL-K80A, clbL-S155A, and clbL-S179A* overexpression plasmids into *E. coli* DH10B BAC*pks* knockout strains.** Linearized pTrcHisA vector was PCR amplified from pTrcHisA vector (Thermo Fisher Scientific) using primers shown in **Supplementary Table 2**. Full-length *clbL* was PCR amplified from *E. coli* CFT073 genomic DNA (purchased from ATCC) using primers shown in **Supplementary Table 2**. PCR reactions (20 μL) contained 10 μL Q5 High-Fidelity 2× Master Mix, 1 ng of DNA template, and 500 pmoles of each primer. Thermocycling was carried out in a C1000 gradient cycler using the conditions shown in **Supplementary Table 3**.

Gibson assembly reactions (10 μL) contained 100 ng linearized pTrcHisA vector, a 3-fold of molar excess purified PCR products of *clbL*, and 5 μL of 2× Gibson Assembly Master Mix. The mixtures were incubated at 50 ºC for 15 min and used to transform 50 μL of electrocompetent *E. coli* NEB® 10-beta cells following the manufacturer's protocol. The identity of the assembled plasmids was confirmed by sequencing.

Site-directed mutagenesis of pTrcHisA-*clbL* was performed using 25 μL of Phusion High-Fidelity PCR Master Mix, 50 ng of template, and 500 pmoles of each primer in a total volume of 50 μL. Thermocycling was carried out in a C1000 gradient cycler using the parameters outlined in **Supplementary Table 3**. The primers used are listed in **Supplementary Table 2**. Digestion of the template in each PCR reaction was performed by the addition of 1 μL of DpnI and incubation at 37 ºC for 1 h, followed by the addition of another 1 μL of DpnI and incubation at 37 ºC for 1 h. 2.5 μL of the digestion reaction was used to transform 50 μL of electrocompetent *E. coli* NEB® 10-beta cells following the manufacturer's protocol. The success of mutagenesis was confirmed by sequencing of purified plasmid DNA.

The pTrcHisA-*clbL, -clbL-K80A, -clbL_S155A,* and *–clbL-S179A* plasmids were electroporated into electrocompetent *E. coli* DH10B BAC*pks*Δ*clbP* or *E. coli* DH10B BAC*pks*Δ*clbP*Δ*clbL* and stored at –80 ºC as frozen 20% (v/v) glycerol/LB stocks.

**HeLa cell infection and cell cycle analysis**. HeLa cells in a 6-well plate were washed twice with DMEM medium supplemented with 10% FBS and infected with bacterial cell suspension in DMEM medium supplemented with 10% FBS and 25 mM HEPES with a multiplicity of infection (MOI) of 1,000 for 1 h. HeLa cells were then washed twice with 1× DPBS and incubated in DMEM medium supplemented with 10% FBS, 1× Antibiotic-Antimycotic, and 200 μg/mL gentamicin.

One day post bacterial infection, HeLa cells were collected, centrifuged at 4 °C (400 × *g* for 2 min), washed with 1× DPBS, and resuspended in 70% ice-cold ethanol for fixation overnight at 4 °C. For DNA content analysis, cells were centrifuged (800 × *g* for 5 min), washed with 1× DPBS, and stained with 1× DPBS supplemented with 20 μg/mL propidium iodide and 200 μg/mL RNase A at room temperature for at least 30 min. Cell cycle was monitored on the BD LSR II flow cytometer (BD Biosciences, Franklin Lakes, NJ) with 10,000 events/determination and analyzed with Flowjo software (Tree Star Inc., Ashland, OR). All cell cycle experiments were replicated three times independently and a cell cycle profile from one representative experiment was shown.

**Metabolite analyses of *E. coli* BAC*pks* knockout strain harboring overexpression plasmids.** A starter culture of an *E. coli* BAC*pks* knockout strain harboring an appropriate overexpression plasmid (5 mL) was inoculated from frozen cell stock and grown overnight at 37 ºC in LB medium supplemented with 50 μg/mL kanamycin, 25 μg/mL chloramphenicol, and 100 μg/mL ampicillin. The starter culture was used to inoculate 50 mL of fresh LB medium containing 50 μg/mL kanamycin, 25 μg/mL chloramphenicol, and 100 μg/mL ampicillin with a normalized number of cells, such that an $OD_{600}$ of 1 of the overnight culture gave a 1:100 volume of inoculum. In specified experiments, the culture was supplemented with 1 mM **7**, 1 mg/mL [1-$^{13}$C]-L-methionine, or 1 mg/mL [1,2-$^{13}C_2$]-glycine at the time of inoculation. The culture was incubated at 37 °C with 200 rpm shaking for 24 h, transferred into 50 mL Falcon tubes, and centrifuged at 4 ºC (4,000 rpm × 10 min). The cell pellets were flash frozen in liquid nitrogen and lyophilized overnight. The resulting dried biomass was then extracted with LC-MS-grade MeOH (Honeywell Research Chemicals, 1 mL) by vortexing. Samples were centrifuged at 4 ºC (13,000 rpm × 20 min), and the supernatant was analyzed by LC/MS.

For the comparative metabolomics experiment, the following liquid chromatography gradient was applied: 0-1.5 min, 98% B isocratic; 1.5-45 min, 98-0% B; 45-48 min, 0% B isocratic; 48-50 min, 0-98% B; 50-60 min, 98% isocratic. The *m/z* values reported correspond to monoisotopic peaks. The chromatographic datasets were aligned by retention time and mass, and the aligned data was statistically analyzed using the XCMS online software (https://xcmsonline.scripps.edu).[6] The results were then extracted from XCMS-processed data using the parameters $p < 0.05$, maximum intensity > 10, 000, and fold change > 5.

For the other experiments, the following liquid chromatography gradient was applied: 0-1 min, 60% B isocratic; 1-9 min, 60-25% B; 9-11 min, 25-15% B; 11-12 min, 15-1% B; 12-15 min, 1% B isocratic; 15-17 min, 1-60% B; 17-21 min, 60% B isocratic. The *m/z* values reported correspond to monoisotopic peaks.

All experiments were performed in positive ionization mode with a mass range of *m/z* 100 to 1700 and a scan speed of 1 scan/sec. The capillary voltage was set to 3500 V. The source parameters were set with a gas temperature of 275 ºC, a flow rate of 8 L/min, and nebulizer at 35 psig. LC/MS/MS analysis of **6** used a collision energy of 30 eV. LC-MS and LC/MS/MS data were acquired with MassHunter Workstation Data Acquisition (Agilent Technologies) and analyzed using MassHunter Qualitative Analysis software (Agilent Technologies). A mass window of ± 10 ppm was used to extract the $[M+H]^+$ ion.

**Isolation and structural characterization of candidate precolibactin 6.** Starter cultures of an *E. coli* DH10B BAC*pks*Δ*clbP* + pTrcHisA-*clbL* strain (3 × 50 mL) were inoculated from a frozen cell stock and grown overnight at 37 ºC in LB medium supplemented with 50 μg/mL kanamycin and 100 μg/mL ampicillin. The starter cultures were used to inoculate 50 × 2 L and 800 × 50 mL (total of 140 L) of fresh LB medium containing 50 μg/mL kanamycin and 100 μg/mL ampicillin with a normalized number of cells, such that an $OD_{600}$ of 1 of the overnight culture gave a 1:100 volume of inoculum. The cultures were incubated at 37 ºC with 200 rpm shaking for 24 h. After the fermentation period, the cells were harvested by centrifugation at 4 ºC (6,000 × *g* for 15 min), flash frozen in liquid nitrogen, and lyophilized overnight to yield 105 g of dried biomass. The biomass was finely ground using a mortar and pestle and extracted with MeOH (3 × 1.5 L; 30 min of stirring at room temperature for each extraction). The resulting MeOH extract was concentrated in vacuo to give 17.2 g of a yellow/brown solid.

The crude organic extract was subjected to two rounds of solid phase purification using a SepPak C18 cartridge (10 g) and eluted using a MeOH/$H_2O$ step gradient (100 mL each step). The elution steps were 25% MeOH/$H_2O$, 50% MeOH/$H_2O$, 75% MeOH/$H_2O$, and 100% MeOH. Based on LC-MS analysis, **6** eluted in the 75% MeOH fraction. The crude product was purified by Sephadex LH-20 column chromatography (100% MeOH isocratic elution). The **6**-containing fraction was collected and further purified by reverse-phase HPLC using a Hypersil GoldTM aQ C18 polar endcapped column (250 × 10 mm, 5 μm particle size, Thermo Fisher Scientific). The following gradient was applied: 0-15 min, 35-32% B; 15-20 min, 32-25% B; 20-20.5 min, 25-0% B; 20.5-23 min, 0% B isocratic; 23-24 min, 0-35% B; 24-30 min, 35% B isocratic (solvent A: acetonitrile + 0.1% formic acid; solvent B: water + 0.1% formic acid; flow rate: 3 mL/min). The product-containing fraction was concentrated *in vacuo* to afford approximately 0.7 mg of **6**. DMSO-$d_6$ was added to the white solid and NMR spectra were obtained in a Shigemi tube matched to DMSO-$d_6$. $^1$H NMR (600 MHz, DMSO-$d_6$): δ (ppm) = 12.17 (br s, 1H), 8.62 (s, 1H), 8.42 (s, 1H), 8.09 (d, $J$ = 7.8 Hz, 1H), 7.83 (d, $J$ = 8.2 Hz, 1H), 7.63 (d, $J$ = 7.8 Hz, 1H), 7.51 (d, $J$ = 8.1 Hz, 1H), 7.24 - 7.17 (m, 3H), 6.79 (s, 1H), 6.07 (s, 1H), 5.53 - 5.40 (m, 2H), 4.44 (q, $J$ = 7.7 Hz, 1H), 3.81 - 3.76 (m, 1H), 3.19 - 3.07 (m, 2H), 2.42 (dd, $J$ = 15.2, 6.3 Hz, 1H), 2.33 (dd, $J$ = 15.2, 7.4 Hz, 1H), 2.03 - 2.00 (m, 2H), 1.66 - 1.57 (m, 2H), 1.49 - 1.34 (m, 2H), 1.41 - 1.38 (m, 2H), 1.35 - 1.32 (m, 2H), 1.27 - 1.19 (m, 20H), 1.00 (d, $J$ = 6.6 Hz, 3H), 0.85 (t, $J$ = 7.0 Hz, 3H). HRMS (ESI): calcd for $C_{41}H_{57}N_6O_6^+$ $[M+H]^+$, 729.4334; found, 729.4344.

**Generation of a glycine auxotrophic *E. coli* strain harboring colibactin biosynthesis genes.** To generate a glycine auxotrophic *E. coli* strain harboring colibactin biosynthesis genes, an *E. coli* JW2535-1 strain was electroporated with BAC*pks*Δ*clbP*Δ*clbL* and pTrcHisA-*clbL* and selected on a LB agar plate containing 100 μg/mL ampicillin, 50 μg/mL kanamycin, and 25 μg/mL chloramphenicol.

**Cloning, overexpression, and purification of ClbL.** The cloning, overexpression, and purification of Sfp, ClbN, ClbB, ClbC, ClbH, and ClbI were reported previously[4, 7]. The gene *clbL* was PCR amplified from *E. coli* CTF073 genomic DNA using the primers shown in **Supplementary Table 2**. The PCR reaction contained 25 μL of Phusion High-Fidelity PCR Master Mix, 2 ng of DNA template, and 500 pmoles of each primer. Thermocycling was carried out in a C1000 gradient cycler using the parameters outlined in **Supplementary Table 3**.

PCR reactions were analyzed by agarose gel electrophoresis with ethidium bromide staining, pooled, and purified. Amplified fragments were digested with NdeI and XhoI for 2.5 h at 37 °C. The digest contained 4 μL of water, 3 μL of 10× NEB Buffer, 20 μL of PCR product, and 1.5 μL of each restriction enzyme (20,000 U/μL). Restriction digests were purified directly using agarose gel electrophoresis. Gel fragments were further purified using an Illustra GFX PCR DNA and Gel Band Purification Kit. The digests were ligated into linearized pET-28a expression vectors using T4 DNA ligase to encode a C-terminal $His_6$-tagged construct.

Ligation reactions contained 3 μL of water, 1 μL of 10× T4 Ligase Buffer, 1 μL of digested vector, 3 μL of digested insert DNA, and 2 μL of T4 DNA Ligase (400 U/μL). Reactions were incubated at 16 °C for approximately 16 h, after which 5 μL of each ligation was used to transform a single tube of chemically competent *E. coli* TOP10 cells (Thermo Fisher Scientific). The identities of the resulting constructs were confirmed by sequencing of purified plasmid DNA. The construct was transformed into chemically competent *E. coli* C43(DE3) cells (Lucigen, Middleton, WI) and stored at -80 °C as frozen 1:1 LB/glycerol stocks.

A 25 mL starter culture of *E. coli* C43(DE3) was inoculated from a frozen stock and grown overnight at 37 °C in LB medium supplemented with 50 μg/mL kanamycin. Overnight cultures were diluted in 1:100 into 2 L of LB medium containing 50 μg/mL kanamycin. Cultures were incubated at 37 °C with shaking at 175 rpm, moved to 25 °C at $OD_{600}$ = 0.2-0.3, induced with 500 μM IPTG at OD600 = 0.5-0.6, and incubated at 25 °C overnight with shaking.

Cells from 2 L of culture were harvested by centrifugation (6,500 rpm × 15 min) and resuspended in 40 mL of lysis buffer (50 mM Tris-HCl pH 8.3, 250 mM NaCl, 10 mM $MgCl_2$, and 10% v/v glycerol). The cells were lysed by passage through cell disruptor (Avestin EmulsiFlex-C3) twice at 7,500 psi, and the lysate was clarified by centrifugation at 4 °C (13,000 rpm × 30 min). The supernatant was incubated with 2 mL of Ni-NTA resin equilibrated with elution buffer (50 mM Tris-HCl pH 8.3, 500 mM NaCl, 10 mM $MgCl_2$, 10% v/v glycerol, 10 mM imidazole) on a nutator for 2 h at 4 °C. The mixture was centrifuged (3,500 rpm × 10 min) and the unbound fraction discarded. The Ni-NTA resin was re-suspended in 1.5 mL of elution buffer containing 10 mM imidazole, loaded onto a glass column, and washed with ~15 mL of elution buffer containing 25 mM imidazole. Protein was eluted from the column using a stepwise imidazole gradient in elution buffer (50 mM, 75 mM, 100 mM, 125 mM, 150 mM, 200 mM), collecting 2 mL fractions. The resin was rinsed with elution buffer containing 250 mM imidazole until all resin-bound protein was eluted. SDS–PAGE analysis (4–15% Tris-HCl gel) was employed to confirm the presence and purity of protein in fractions. Fractions containing the desired protein were combined and dialyzed twice against 2 L of storage buffer (50 mM Tris–HCl pH 8.3, 200 mM NaCl, 10 mM $MgCl_2$, 10% v/v glycerol). Solutions containing protein were concentrated with a Spin-X concentrator (Corning) to 0.8–1.0 mL and centrifuged at 4 °C (13,200 rpm × 10 min) to remove particulate. Protein was further purified by gel filtration chromatography on a GE HighLoad 26/600 Superdex 200 pg column. This procedure yielded 0.5–2 mg/L of purified ClbL.

**In vitro reconstitution of 6 and 12**. The assay for reconstituting the formation of metabolites **6** and **12** contained 50 mM Tris-HCl (pH 8.3), 200 mM NaCl, 10 mM $MgCl_2$, 1 mM TCEP, 1.7% DMSO, 125 μM CoA, 500 μM myristoyl-CoA, 500 μM malonyl-CoA, 2 mM NADPH, 0.3 mM L-Asn, 0.3 mM L-Ala, 0.3 mM SAM, 2.3 mM ATP, 3 μM Sfp, 5 μM ClbN, 5 μM ClbB, 5 μM ClbC, 5 μM ClbH, 10 μM ClbI, 6 μM ClbL, and 1 mM **7** (for the reconstitution of **6**) or 2-aminoacetophenone (for the reconstitution of **12**) in a total volume of 25 μL. ClbN, ClbB, ClbC, ClbH, and ClbI were incubated with Sfp and CoA at 22 °C for 1.5 h for phosphopantetheinylation. The complete reaction was initiated by the addition of myristoyl-CoA, malonyl-CoA, NADPH, L-Asn, L-Ala, SAM, ATP, ClbL, and **7** or 2-aminoacetophenone.

The assay mixtures were incubated at 22 °C for 16 h, quenched by the addition of 125 μL ice-cold methanol, incubated on ice for 30 min, and then centrifuged at 4 °C (13,200 rpm × 15 min). The supernatant was saved for LC-MS and LC-MS/MS analysis. The following gradient was applied: 0-1 min, 60% B isocratic; 1-9 min, 60-25% B; 9-11 min, 25-15% B; 11-12 min, 15-1% B; 12-15 min, 1% B isocratic; 15-17 min, 1-60% B; 17-21 min, 60% B isocratic. The *m/z* values reported correspond to monoisotopic peaks.

**In vitro assay of ClbL.** The assay for examining the substrate scope of ClbL contained 50 mM Tris-HCl (pH 8.3), 200 mM NaCl, 10 mM $MgCl_2$, 6 μM ClbL, 1 mM indicated thioester substrate, and 1 mM indicated aminoketone substrate in a total volume of 25 μL. Synthesis of indicated thioester substrates was described in **Synthetic procedures**. The aminoketone substrates **13**-**15** were purchased from Enamine Ltd. The assay mixtures were incubated at 22 °C for 16 h, quenched by the addition of 125 μL ice-cold methanol, incubated on ice for 30 min, and centrifuged at 4 °C (13,200 rpm × 15 min). The supernatant was saved for HPLC, LC-MS, and LC-MS/MS analysis. The following liquid chromatography gradient was applied: 0-1 min, 95% B isocratic; 1-12 min, 95-5% B; 12-15 min, 5% B isocratic; 15-17 min, 5-95% B; 17-21 min, 95% B isocratic. The *m/z* values reported correspond to monoisotopic peaks. LC/MS/MS analysis of **9** used a collision energy of 10 eV.

**Preparation of cross-links of linearized plasmid DNA.** The 5.4 kb pcDNA3 plasmid was digested with BamHI-HF overnight. The linearized plasmid was purified using the DNA Clean & Concentrator kit (Zymo Research). In a 15 mL Falcon tube, $7.2 \times 10^7$ bacteria from an overnight culture of *pks*$^-$ or *pks*$^+$ *E. coli* was added to 2.4 mL DMEM medium supplemented with 60 μL 1 M HEPES and 9.6 μg of linearized pcDNA3. In some experiments, the culture was added 1 mg/mL [2,3-$^{13}C_2$]-L-alanine, [1-$^{13}$C]-L-methionine, or [1-$^{13}$C]-L-cysteine at the time of inoculation. The resulting mixture was incubated at 37 °C for 4 h and bacteria was pelleted by centrifugation at 4 °C (13,200 rpm × 5 min). The cross-linked DNA was purified from the supernatant using the DNA Clean & Concentrator kit (Zymo Research) and used for DNA adductomics.

**DNA enzymatic digestion.** The amounts of purified cross-linked DNA were estimated by dissolving the samples in water and measuring the concentrations by a NanoDrop 2000 UV-Vis Spectrophotometer. DNA concentrations ranged from 2-19 μg. The DNA digestion was carried out using a cocktail of enzymes consisting of recombinant DNase I expressed in *Pichia pastoris* (R-DNase, 10000 U·mg$^{-1}$), phosphodiesterase-1 from *Crotalus adamanteus* venom (PDE-1, 0.4 U·mg$^{-1}$), and recombinant alkaline phosphatase highly active expressed in *Pichia pastoris* (R-ALP, 7000 U·mg$^{-1}$), all purchased from MilliporeSigma (Burlington, MA, USA). All enzymes were purified prior to use by passage of the enzyme solution through an Amicon Ultra (Burlington, MA, USA) double filtration membrane (0.5 mL, cutoff 10 kDa). The hydrolysis consisted of two steps: an initial 24 h incubation with DNase at room temperature and a 70 min incubation at 37 °C with DNase, PDE-1, and ALP followed by 24 h incubation at room temperature. The first step used 0.5 U/μg DNA of DNase and the second step used 0.5 U/μg DNA, 0.2 U/μg DNA and 0.02 mU/μg DNA of DNase, ALP and PDE-1, respectively. The enzymes were removed using an Amicon Microcon (Burlington, MA, USA) single filtration membrane (0.5 mL, cutoff 10 kDa).

**Off-line reversed-phase HPLC Purification**. Prior to sample analysis, unmodified deoxynucleosides were removed by off-line reversed-phase HPLC purification, performed using an HPLC (Ultimate 3000, Thermo Fisher Scientific) equipped with a C18 column (2 × 250 mm, Luna C18, 5 µm, 100 Å, Phenomenex, Torrance, CA) maintained at 45 °C, with a flow rate of 0.4 mL/min. A and B mobile phases were $H_2O$ and MeOH, respectively, and the linear gradient increased from 2% to 16.5% B in 11 min followed by an isocratic step at 100% of B (15 min). UV absorption at 190 and 254 nm with 4 Hz detection was used for monitoring the elution of the deoxyribonucleosides. After elution of deoxyadenosine (dA) at 12 min, the eluting liquid was collected for 8 min, completely dried and stored at –20 °C.

**Chromatography for LC-MS**. Mass spectrometric data was acquired with the following conditions. Each dried sample was reconstituted in 20 µL of $H_2O$ and 5 µL of sample, corresponding to DNA amounts ranging from 0.5 - 4.75 µg, were injected onto an UltiMate 3000 RSLCnano UPLC (Thermo Fisher Scientific) system equipped with a 5 µL injection loop. Separation was performed with a capillary column (75 µm ID, 20 cm length, 10 µm orifice) created by hand packing a commercially available fused-silica emitter (New Objective, Woburn, MA) with 5 µm Luna C18 bonded separation media (Phenomenex, Torrance, CA). The flow rate was 1000 nL/min for 5.5 min at 2% $CH_3CN$ in 0.05% formic acid aqueous solution, then decreased to 300 nL/min followed by a linear gradient of 1.23%/min over 39 min for the untargeted screening and over 21 min for the targeted SIM analysis. The column was washed at 95% $CH_3CN$ for 2 min and re-equilibrated at 2% $CH_3CN$ with a flow rate of 1000 nL/min over 2 min. The injection valve was switched at 5.5 min to remove the sample loop from the flow path during the gradient.

**Mass Spectrometry.** All mass spectrometric data was acquired with a Fusion mass spectrometer (Thermo Fisher Scientific). Positive mode electrospray ionization was used under nanospray conditions (300 nL/min) using a Thermo Scientific Nanoflex ion source with a source voltage of 2.2 kV, and the capillary temperature was 300 °C. The S-Lens RF level setting was 60%.

**DNA Adduct Untargeted Screening**. Data-dependent constant neutral loss (CNL)-$MS^3$ analysis was performed with repeated full scan detection followed by $MS^2$ acquisition and constant neutral loss triggering of $MS^3$ fragmentation. Full scan (*m/z* 500 − 2000) detection was performed using the Orbitrap detector at a resolution setting of 120,000, automatic gain control (AGC) target settings of $1 \times 10^6$, and a maximum ion injection time setting of 50 ms. $MS^2$ spectra were acquired with quadrupole isolation of *m/z* 1.5 and HCD fragmentation with stepped collision energy of 15% with +/- 10% (5, 15, 25%) and Orbitrap detection at a resolution setting of 15000, AGC setting of $2 \times 10^5$, and maximum ion injection time of 50 ms. Data-dependent parameters were as follows: triggering range of $1 \times 10^4$ to $1 \times 10^7$, repeat count of 1, exclusion duration of 15 s, and exclusion mass width of ±5 ppm. A mass exclusion list consisting of 56 previously observed unmodified nucleosides-derived ions was used. The "Monoisotopic Peak Determination" feature was enabled and set to "Small molecule" mode. $MS^3$ HCD fragmentation (*m/z* 2.5 isolation width, HCD collision energy of 30%) with Orbitrap detection at a resolution setting of 15000 was triggered upon observation of neutral losses (±5 ppm) of *m/z* 116.0474 (-dR), 232.0947 (-2dR), 151.0494 (-G), 169.0600 (-G, -$H_2O$), 135.0545 (-A), 153.0651 (-A, -$H_2O$), 126.0429 (-T), and 111.0433 (-C) between the parent ion and product ions from the

$MS^2$ spectrum, provided a minimum signal of $5 \times 10^3$ was observed. The following $MS^3$ parameters were used: AGC setting of $2 \times 10^5$, maximum ion injection time of 50 ms. All spectra were acquired with the EASY-IC lock mass (*m/z* 202.0777) enabled. A cycle speed of 3 s was used.

**DNA Adduct Targeted Selected Ion Monitoring (SIM)**. Targeted analysis of the *m/z* 568.1721 and *m/z* 938.2855 adducts generated by growing the plasmids in isotopically-heavy amino acids were analyzed using a quadrupole isolation of wide SIM mass ranges of *m/z* 566.5 – 573.5, *m/z* 936.5 – 943.5, and *m/z* 468.5 – 472.5, for the single charge state of the *m/z* 568.1721 adduct, the single charge state of the *m/z* 938.2855 adduct and its double charge state, respectively. Orbitrap detection was performed at a resolution setting of 120,000 with automatic gain control (AGC) target settings of $2 \times 10^5$, a maximum ion injection time setting of 250 ms, and the EASY-IC (*m/z* 202.0777) lock mass enabled.

### 2. Supplementary tables

**Supplementary Table 1.** Oligonucleotides used for generation of KO resistance cassettes. Regions homologous to the targeted genes are highlighted in red.

| Primer Name | Target | Sequence (5' to 3') |
|---|---|---|
| *clbL*_KO_F | *clbL* | ATG GCG CAG TTT AAC AAG CCG TTG AAT GCG GTG GTG CAG ATT CCG GGG ATC CGT CGA CC |
| *clbL*_KO_R | *clbL* | TAT GCC GCA CGG AAG CTC ATC CAT CGT TTT CGC CAA TGG TGT AGG CTG GAG CTG CTT C |
| *clbH*_KO_F | *clbH* | TTG TAT CGT ATC GCC GGA GAA TAT GGG GAA AAA GCC GCT ATT CCG GGG ATC CGT CGA CC |
| *clbH*_KO_R | *clbH* | CGC CGC ACG CAG TTG TGT CTG ATC TCC TGT GGT CCC TTG TGT AGG CTG GAG CTG CTT C |
| *clbI*_KO_F | *clbI* | GGC GTT TCC CTC AAG CCG ATA CGG TAC AGG CGT TTT GGG ATT CCG GGG ATC CGT CGA CC |
| *clbI*_KO_R | *clbI* | CAT GTC GTT AAC TAG CAC GGC AAG TGC GGA CCC TCC ATC TGT AGG CTG GAG CTG CTT C |
| *clbJ*_KO_F | *clbJ* | TTA CGA CAG TTG GCG CAA TCA GGC GTG TCG CCC AGC CGC ATT CCG GGG ATC CGT CGA CC |
| *clbJ*_KO_R | *clbJ* | ACG CTG CTG TGT TTG ATT CAC CGC CCG TGC ATT GTC CTG TGT AGG CTG GAG CTG CTT C |
| *clbK*_KO_F | *clbK* | CAC AGC GTT GCG GCA TTC GAA ACG GTG CTT CGC ACC GGA ATT CCG GGG ATC CGT CGA CC |
| *clbK*_KO_R | *clbK* | CGC AGC GCT GAC CTG CGA AAC CTC AGC CTG CGC TAA TAC TGT AGG CTG GAG CTG CTT C |

**Supplementary Table 2.** Oligonucleotides used for cloning. Mutation sites are in red. Restriction sites are underlined.

| Primer Name | Target | Sequence (5' to 3') |
|---|---|---|
| GA151 | *pTrc* | GGT TTA TTC CTC CTT ATT TAA TCG ATA CAT TAA TAT ATA |
| GA152 | *pTrc* | CTT CTG CGT TCT GAT TTA ATC TGT A |
| GA161 | *clbL* | AAT AAG GAG GAA TAA ACC ATG AGT GAG CAG AGC TAT C |
| GA162 | *clbL* | CGT TCT GAT TTA ATC TGT ACT AGT ACC CTT CCG GTA C |
| ClbL_K80A_F | *clbL* | CCT TGT ACT GTT GCA GAG TCG TTT GAC |
| ClbL_K80A_R | *clbL* | GTC AAA CGA CTC TGC AAC AGT ACA AGG |
| ClbL_S155A_F | *clbL* | CGT TCA CCC GGC GGA GCG TCC GGA GGA GCG GCG |
| ClbL_S155A_R | *clbL* | CGC CGC TCC TCC GGA CGC TCC GCC GGG TGA ACG |
| ClbL_S179A_F | *clbL* | GAT TTG TTT GGC GCA CTG CGC ATT CCC |
| ClbL_S179A_R | *clbL* | GGG AAT GCG CAG TGC GCC AAA CAA ATC |
| ClbL_cHis_fwd_NdeI | *clbL* | GTC CTA <u>CAT ATG</u> AGT GAG CAG AGC TAT CG |
| ClbL_cHis_rev_XhoI | *clbL* | TAT AAT <u>CTC GAG</u> GTA CCC TTC CGG TAC |

**Supplementary Table 3.** PCR thermocycling conditions for genes listed in **Supplementary Tables 1 and 2**.

| disruption cassette | |
|---|---|
| 98 ºC for 1 min | 1 cycle |
| 98 ºC for 10 sec | 35 cycles |
| 72 ºC for 1.5 min | |
| 72 ºC for 5 min | 1 cycle |

| *pTrc* | | *clbL* cloned into pTrc vector | |
|---|---|---|---|
| 98 ºC for 30 sec | 1 cycle | 98 ºC for 30 sec | 1 cycle |
| 98 ºC for 10 sec | 35 cycles | 98 ºC for 10 sec | 35 cycles |
| 63 ºC for 30 sec | | 61 ºC for 30 sec | |
| 72 ºC for 2.5 min | | 72 ºC for 1 min | |
| 72 ºC for 5 min | 1 cycle | 72 ºC for 5 min | 1 cycle |

| *clbL* site-directed mutagenesis | | *clbL* cloned into pET-28a vector | |
|---|---|---|---|
| 98 ºC for 30 sec | 1 cycle | 98 ºC for 30 sec | 1 cycle |
| 98 ºC for 30 sec | 20 cycles | 98 ºC for 10 sec | 35 cycles |
| 72 ºC for 18 min | | 72 ºC for 3 min | |
| | | 72 ºC for 10 min | 1 cycle |

**Supplementary Table 4.** Down-regulated metabolites from XCMS analysis are sorted by their intensities in wild-type ClbL-expressing strains and ten most abundant metabolites are shown. Candidate precolibactin **6** is highlighted in red.

| | **fold change** | **p-value** | **UP /DOWN** | ***m/z*** | **retention time (min)** | **Intensity: WT ClbL** | **Intensity: mut ClbL** |
|---|---|---|---|---|---|---|---|
| 1 | 13 | 0.00064 | DOWN | 547.3883 | 29.08 | 4.61E+07 | 3.45E+06 |
| 2 | 204472 | 0.00466 | DOWN | 732.4369 | 30.96 | 1.87E+07 | 0.00E+00 |
| 3 | 39 | 0.00237 | DOWN | 1093.7691 | 29.08 | 9.90E+06 | 2.52E+05 |
| 4 | 184 | 0.00010 | DOWN | 572.3835 | 26.55 | 5.17E+06 | 2.81E+04 |
| 5 | 13 | 0.00096 | DOWN | 205.134 | 29.08 | 4.93E+06 | 3.77E+05 |
| 6 | 7 | 0.02713 | DOWN | 454.3656 | 30.25 | 4.15E+06 | 6.22E+05 |
| 7 | 7 | 0.00017 | DOWN | 561.4036 | 30.07 | 3.57E+06 | 5.09E+05 |
| 8 | 6 | 0.00011 | DOWN | 1115.7515 | 29.09 | 5.02E+06 | 8.60E+05 |
| 9 | 1251 | 0.00087 | DOWN | 729.4368 | 29.44 | 2.49E+06 | 1.99E+03 |
| 10 | 2369 | 0.00026 | DOWN | 483.356 | 26.76 | 2.78E+06 | 1.17E+03 |

**Supplementary Table 5.** NMR data of isolated candidate precolibactin **6** recorded in DMSO-$d_6$.

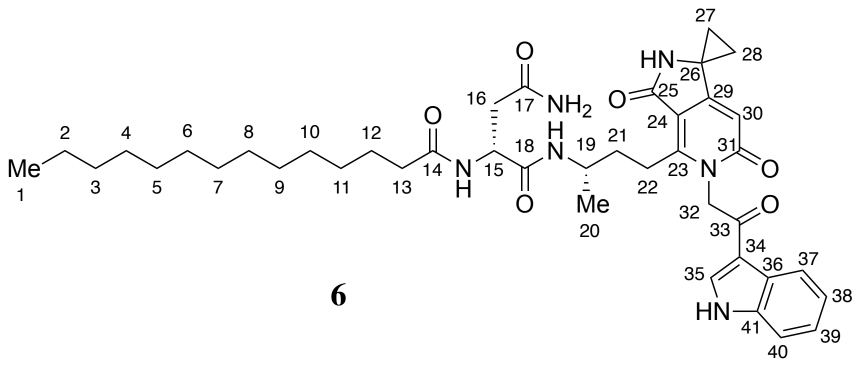


**6**

| Carbon | δC | δH (multiplicity, *J* in Hz) | gCOSY ($^1$H to $^1$H) | gHMBC ($^1$H to $^{13}$C) |
|---|---|---|---|---|
| 1 | 13.1 | 0.85 (t, 7.0) | 2 | 2, 3 |
| 2 | 22.0 | 1.30-1.22 (m) | 1 | – |
| 3 | 31.7 | 1.27-1.19 (m) | – | – |
| 4-11 | 29.42, 29.50 | 1.31-1.19 (m) | 12 | – |
| 12 | 24.2 | 1.49-1.34 (m) | 11, 13 | 11 |
| 13 | 34.3 | 2.03-2.00 (m) | 12 | 11, 12, 14 |
| 14 | 172.3 | – | – | – |
| | | NH, 7.83 (d, 8.2) | 15 | 14 |
| 15 | 49.1 | 4.44 (q, 7.7) | NH, 16 | 16, 18 |
| 16 | 36.5 | 2.42 (dd, 15.2, 6.3), 2.33 (dd, 15.2, 7.4) | 15 | 15, 18 |
| 17 | – | – | – | – |
| | | $NH_2$, 6.79 (s) 7.24-7.17 (m) | – | – |
| 18 | 170.8 | – | – | – |
| | | NH, 7.63 (d, 7.8) | 19 | 18 |
| 19 | 43.7 | 3.81-3.76 (m) | NH, 20 | – |
| 20 | 19.4 | 1.00 (d, 6.6) | 19 | 19, 21 |
| 21 | 34.4 | 1.66-1.57 (m) | 19 | – |
| 22 | 23.6 | 3.19-3.07(m) | 21 | – |
| 23 | 134.4 | – | – | – |
| 24 | 108.9 | – | – | – |
| 25 | 167.4 | – | – | – |
| | | NH, 8.42 (s) | – | 24, 25, 26, 29 |
| 26 | 40.2 | – | – | – |
| 27 | 15.5 | 1.41 - 1.38 (m) 1.35 - 1.32 (m) | 28 | 26, 29 |
| 28 | 15.5 | 1.41 - 1.38 (m) 1.35 - 1.32 (m) | 27 | 26, 29 |

| | | | | |
|---|---|---|---|---|
| 29 | 159.7 | – | – | – |
| 30 | 101.8 | 6.07 (s) | – | 23, 24, 26, 31 |
| 31 | 162.3 | – | – | – |
| 32 | 48.7 | 5.53-5.40 (m) | – | – |
| 33 | – | – | – | – |
| 34 | 114.2 | – | – | – |
| 35 | 133.4 | 8.62 (s) | – | 34, 36, 41 |
| 36 | 125.6 | – | – | – |
| 37 | 120.2 | 8.09 (d, 7.8) | 38 | 39, 41 |
| 38 | 121.0 | 7.23-7.17 (m) | 37 | 36, 40 |
| 39 | 122.0 | 7.23-7.17 (m) | 40 | 37, 41 |
| 40 | 111.3 | 7.51 (d, 8.1) | 39 | 36, 38 |
| 41 | 136.8 | – | – | – |
| | | NH, 12.17 (br s) | – | – |

### 3. Supplementary figures

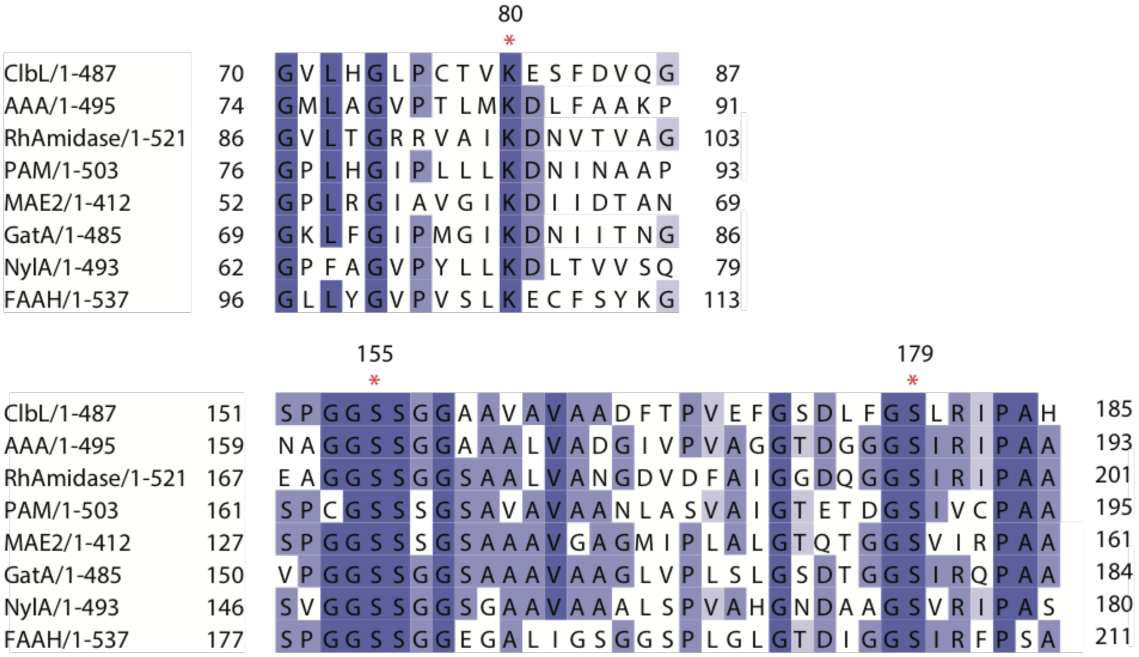


**Supplementary Figure 1.** Alignment of ClbL with structurally characterized amidases, including AAA (*Bacterium* CSBL00001 bacterial aryl acylamidase), RhAmidase (*Rhodococcus sp.* N-771 amidase), PAM (*Stenotrophomonas maltophilia* native peptide amidase), MAE2 (*Bradyrhizobium japonicum* malonamidase E2), GatA (*Staphylococcus aureus* tRNA-dependent amidotransferase subunit A), NylA (*Flavobacterium sp.* 6-aminohexanoate cyclic dimer hydrolase), and FAAH (*Rattus norvegicus* fatty acid amide hydrolase). The predicted catalytic triad is marked with asterisks.

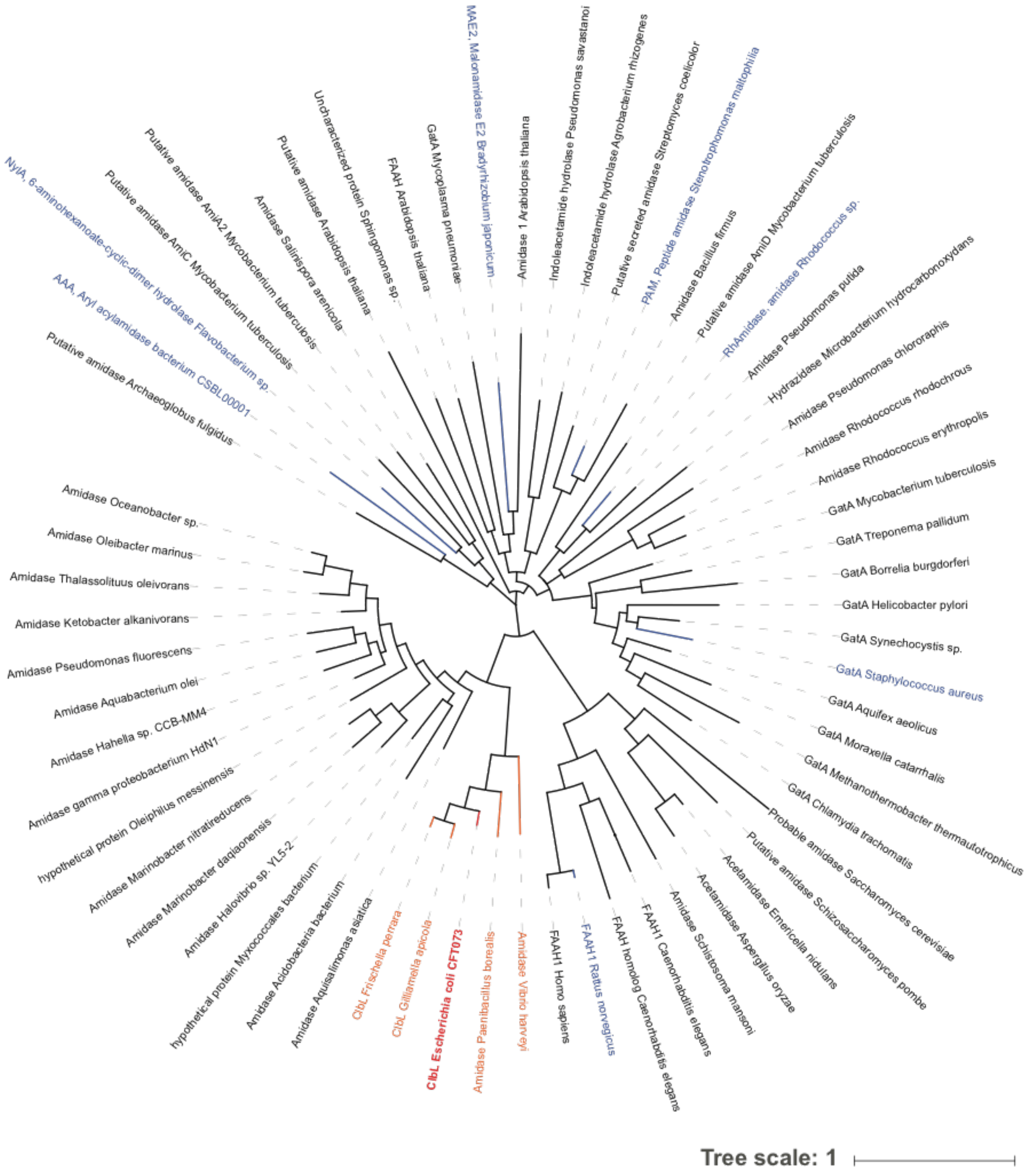


**Supplementary Figure 2.** Phylogenetic analysis of ClbL from *E. coli* (highlighted in red) with members of the amidase signature family enzymes. Structurally characterized amidases are highlighted in blue. Amidases from other species containing the *pks* island are highlighted in orange.

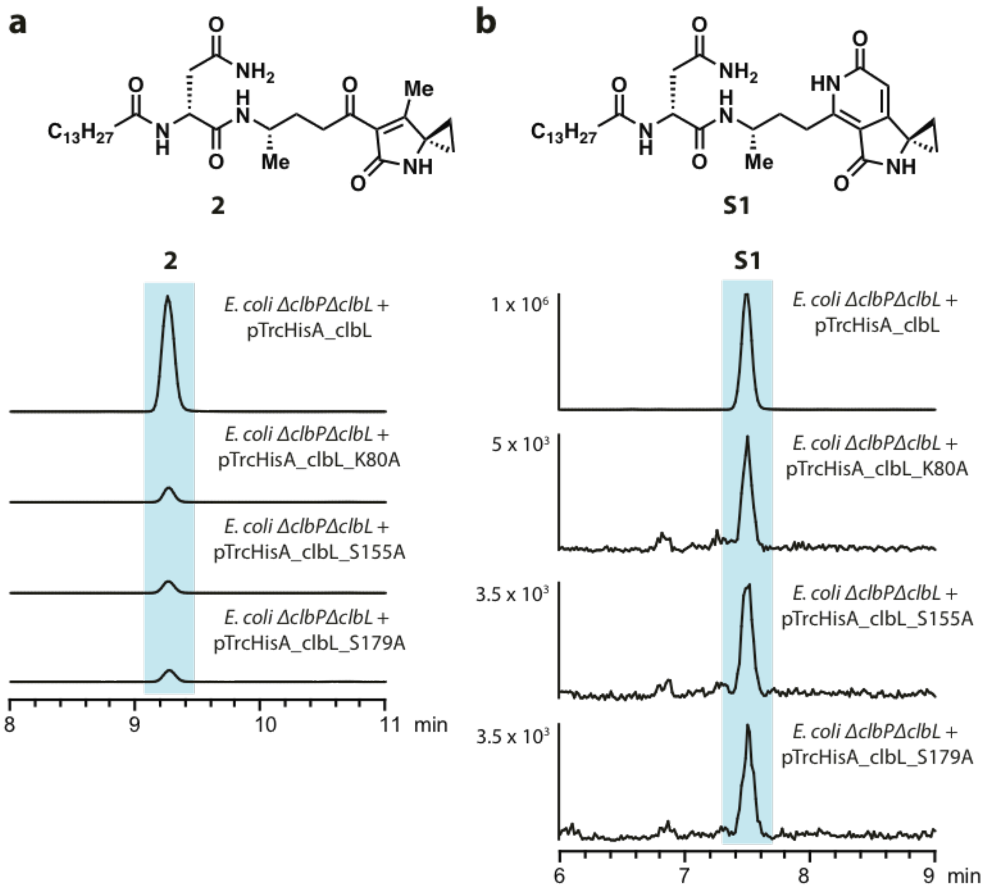


**Supplementary Figure 3.** (**a**-**b**) Extracted ion chromatograms (EICs) of known candidate precolibactin **2** (*m/z* 547.3854) (**a**) and **S1** (*m/z* 572.3806) (**b**) in *E. coli* strains expressing wild-type or mutant ClbL.

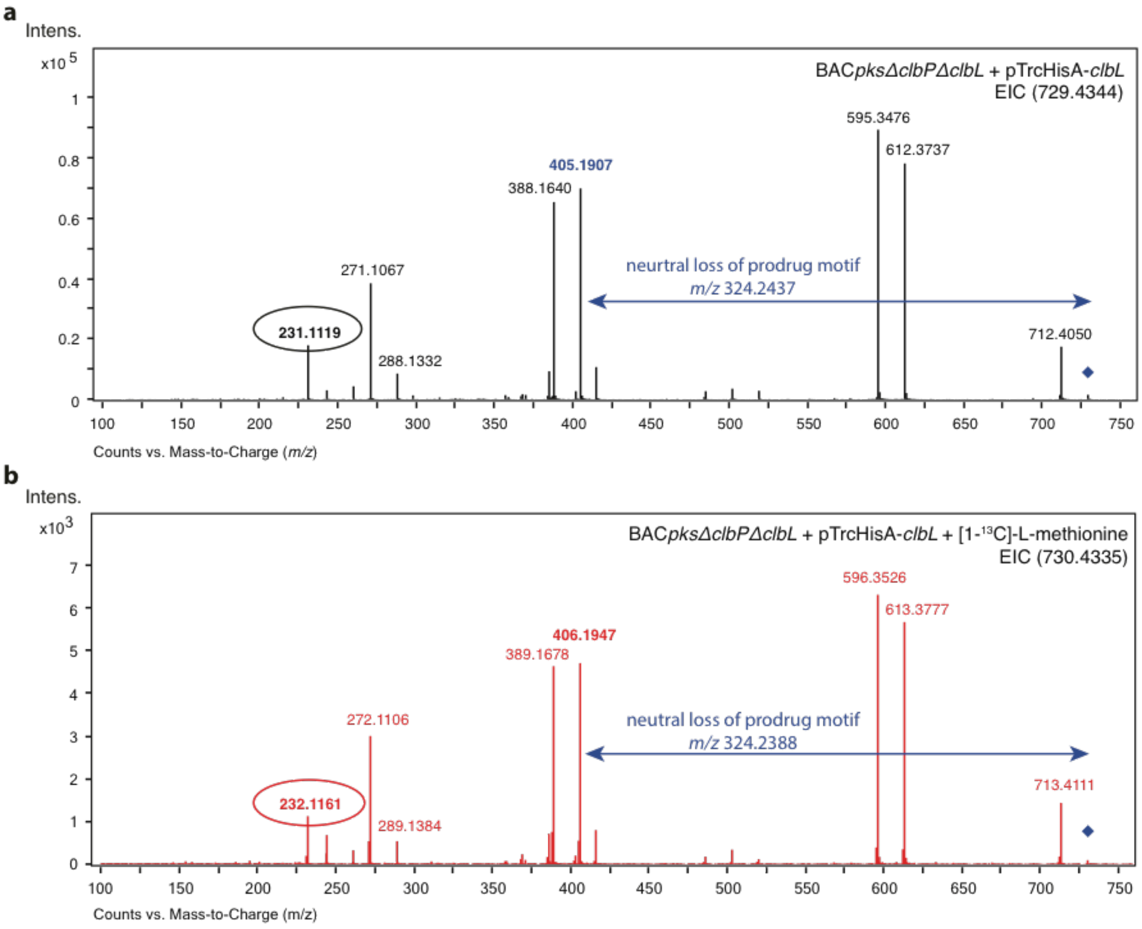


c

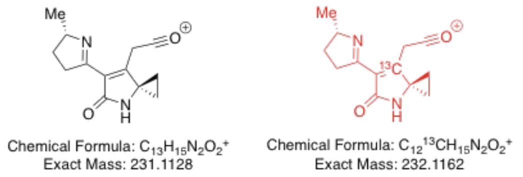


| Species | Theoretical *m/z* | Detected *m/z* | Error (ppm) |
|---|---|---|---|
| $C_{13}H_{15}N_2O_2^+$ | 231.1128 | 231.1119 | 3.9 |
| $C_{12}{}^{13}CH_{15}N_2O_2^+$ | 232.1162 | 232.1161 | 0.4 |

d

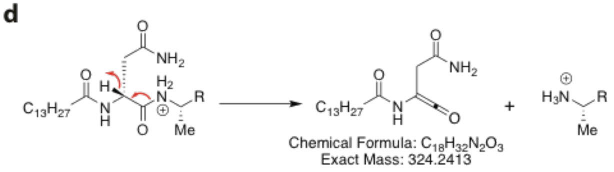


| Species | Theoretical *m/z* | Detected *m/z* | Error (ppm) |
|---|---|---|---|
| $C_{18}H_{32}N_2O_3$ | 324.2413 | 324.2437 | 7.4 |
| $C_{18}H_{32}N_2O_3$ | 324.2413 | 324.2388 | 7.7 |

**Supplementary Figure 4.** (**a**) MS/MS analysis of metabolite **6** in *E. coli* strains expressing ClbL. (**b**) MS/MS analysis of metabolite **6** in *E. coli* strains supplemented with 1 mg/mL [1-$^{13}$C]-L-methionine. (**c**) Predicted structure of the fragment ion *m/z* 231.1128. (**d**) Predicted fragmentation pattern for a neutral loss of *m/z* 324.2413.

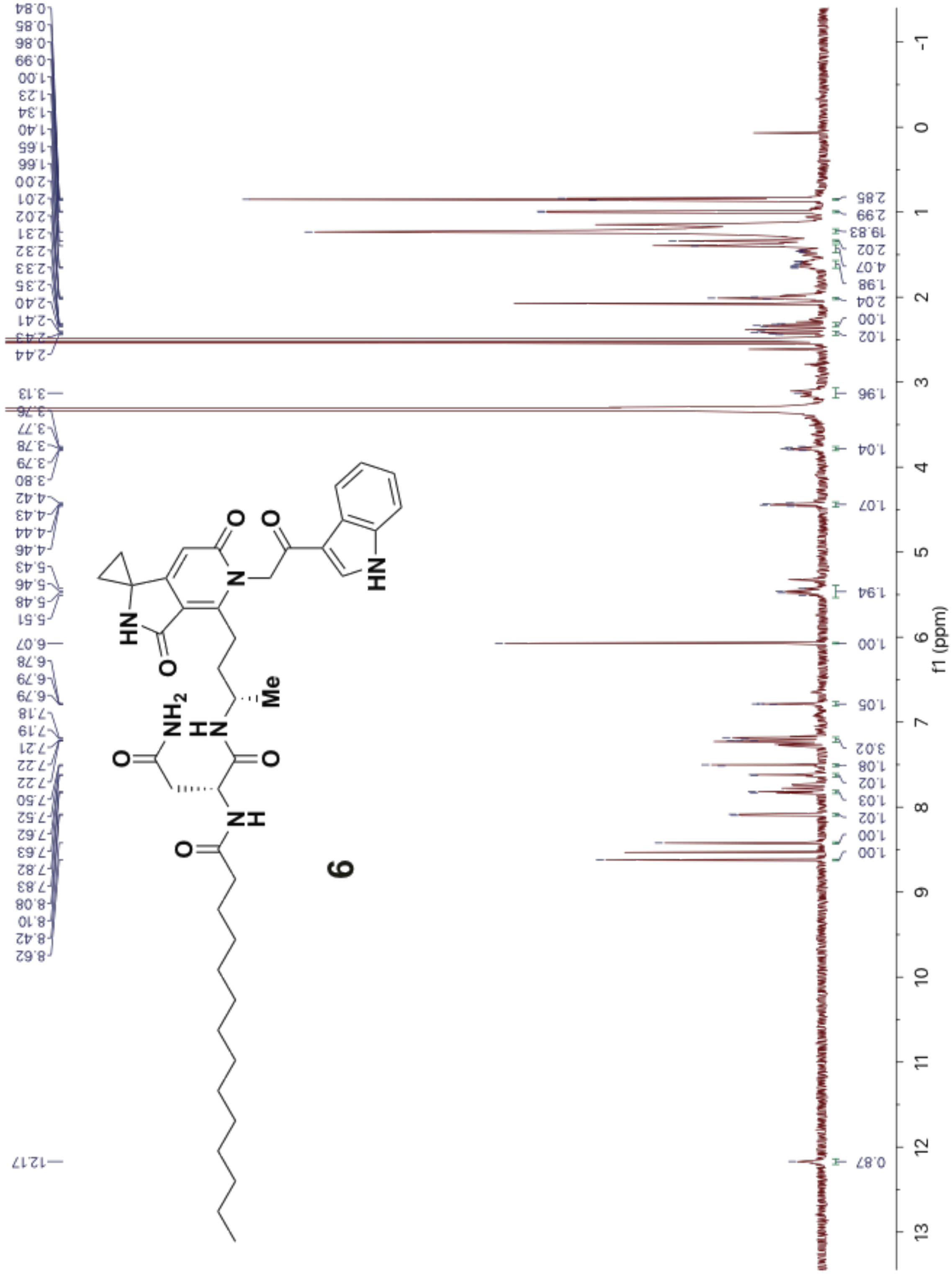


**Supplementary Figure 5.** $^{1}$H-NMR spectrum of candidate precolibactin **6** (recorded in DMSO-$d_6$ at 600 MHz).

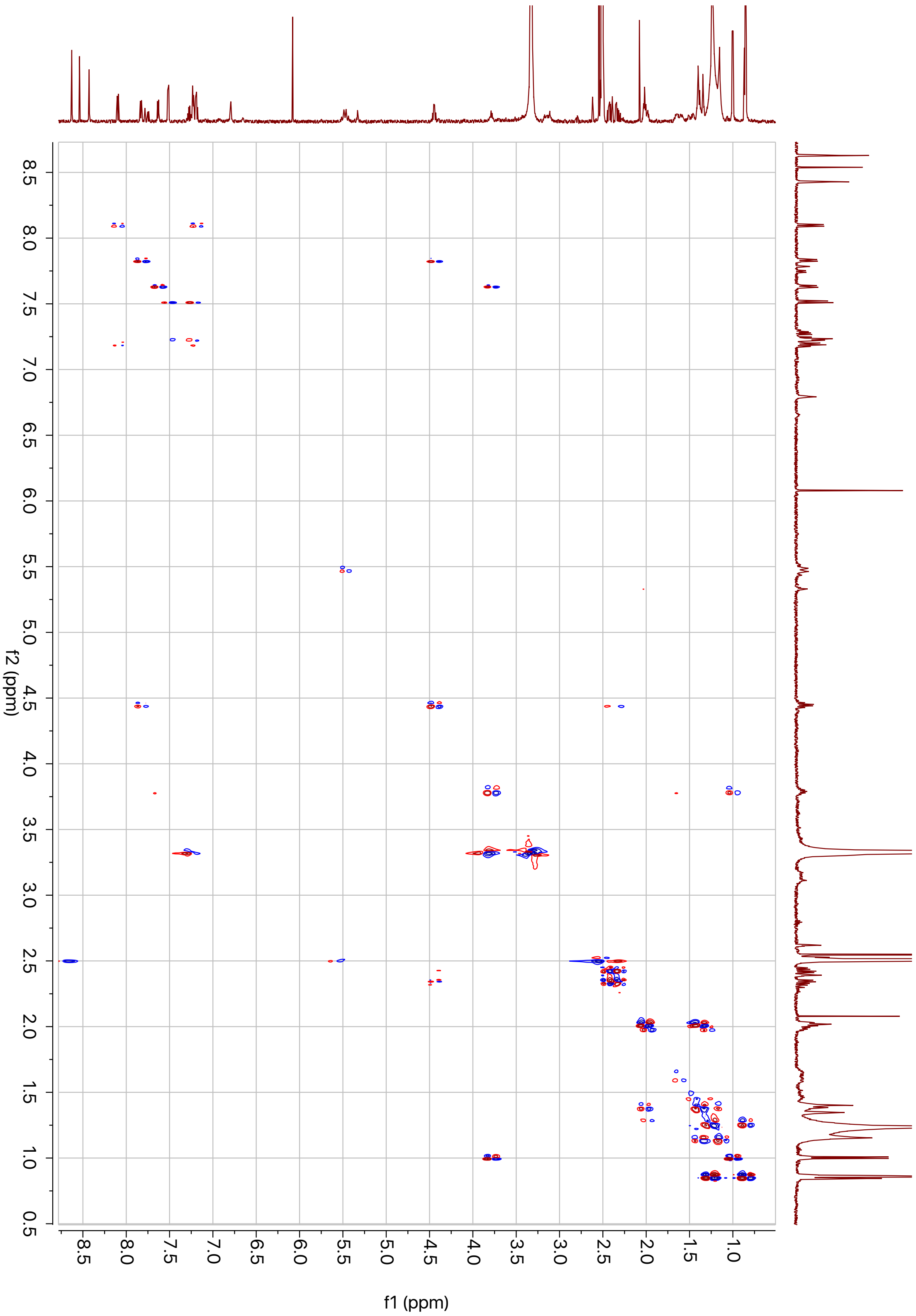


**Supplementary Figure 6.** gCOSY spectrum of candidate precolibactin **6** (recorded in DMSO-$d_6$ at 600 MHz).

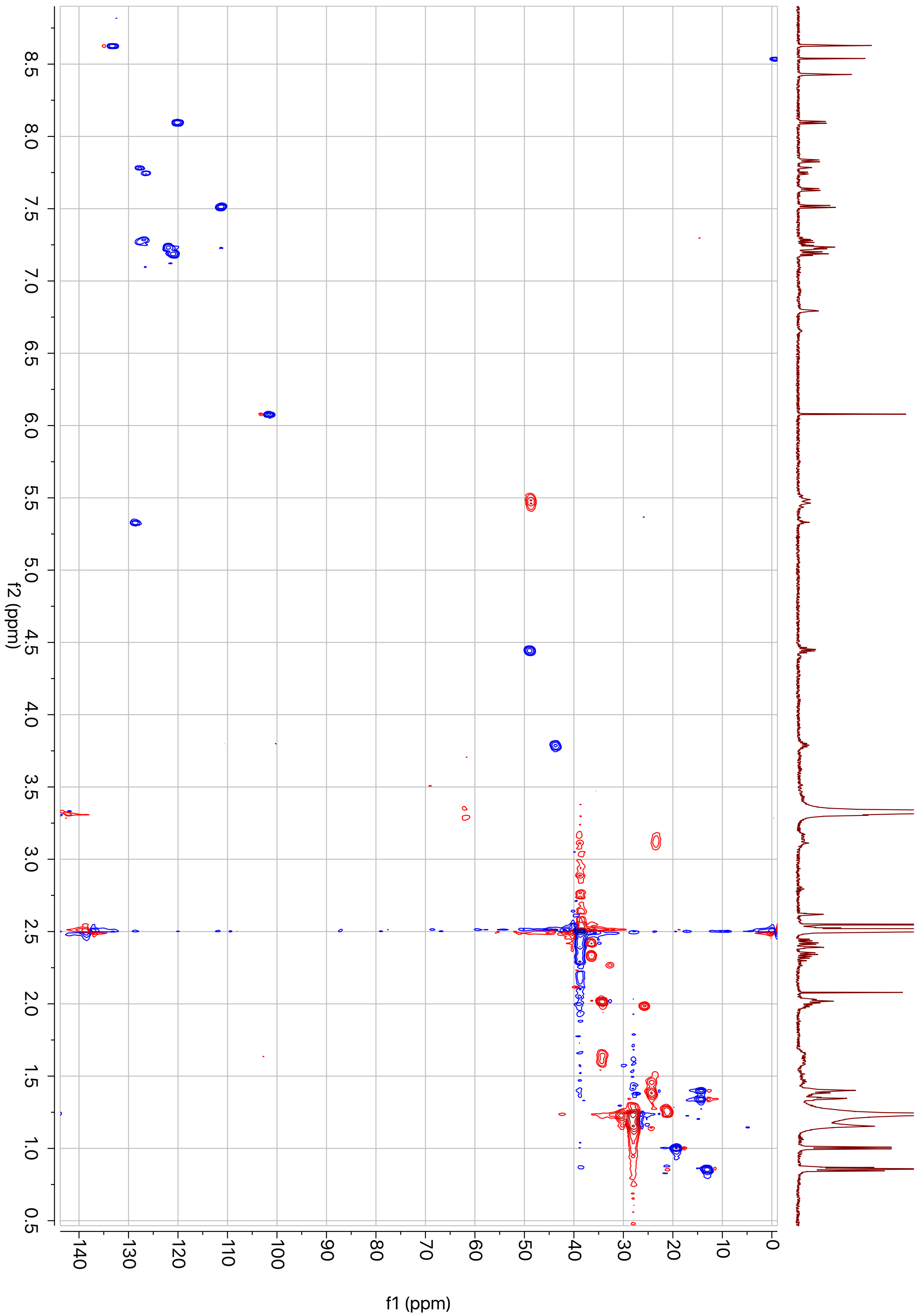


**Supplementary Figure 7.** HSQC spectrum of candidate precolibactin **6** (recorded in DMSO-$d_6$ at 600 MHz).

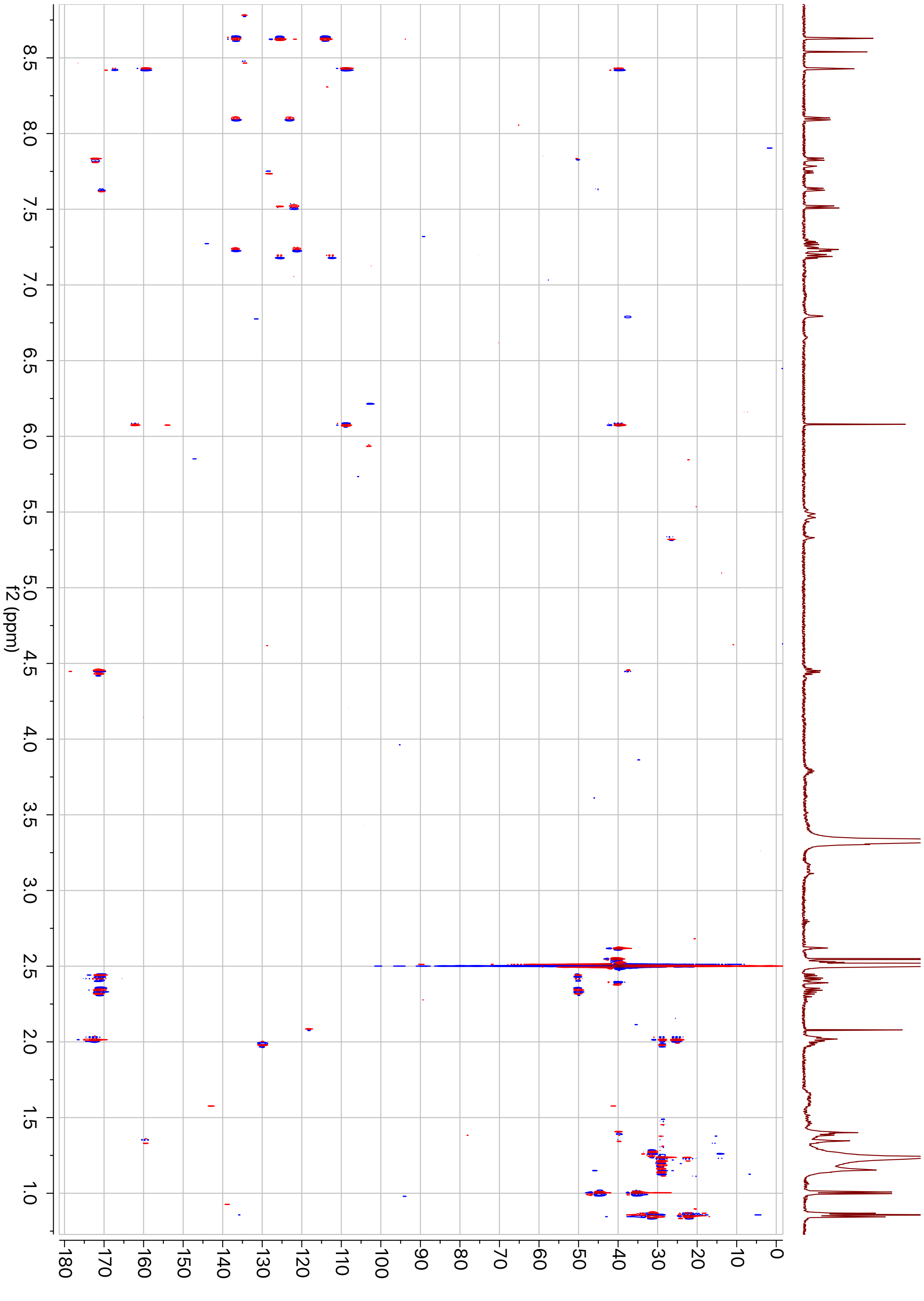


**Supplementary Figure 8.** HMBC spectrum of candidate precolibactin **6** (recorded in DMSO-$d_6$ at 600 MHz).

a

b

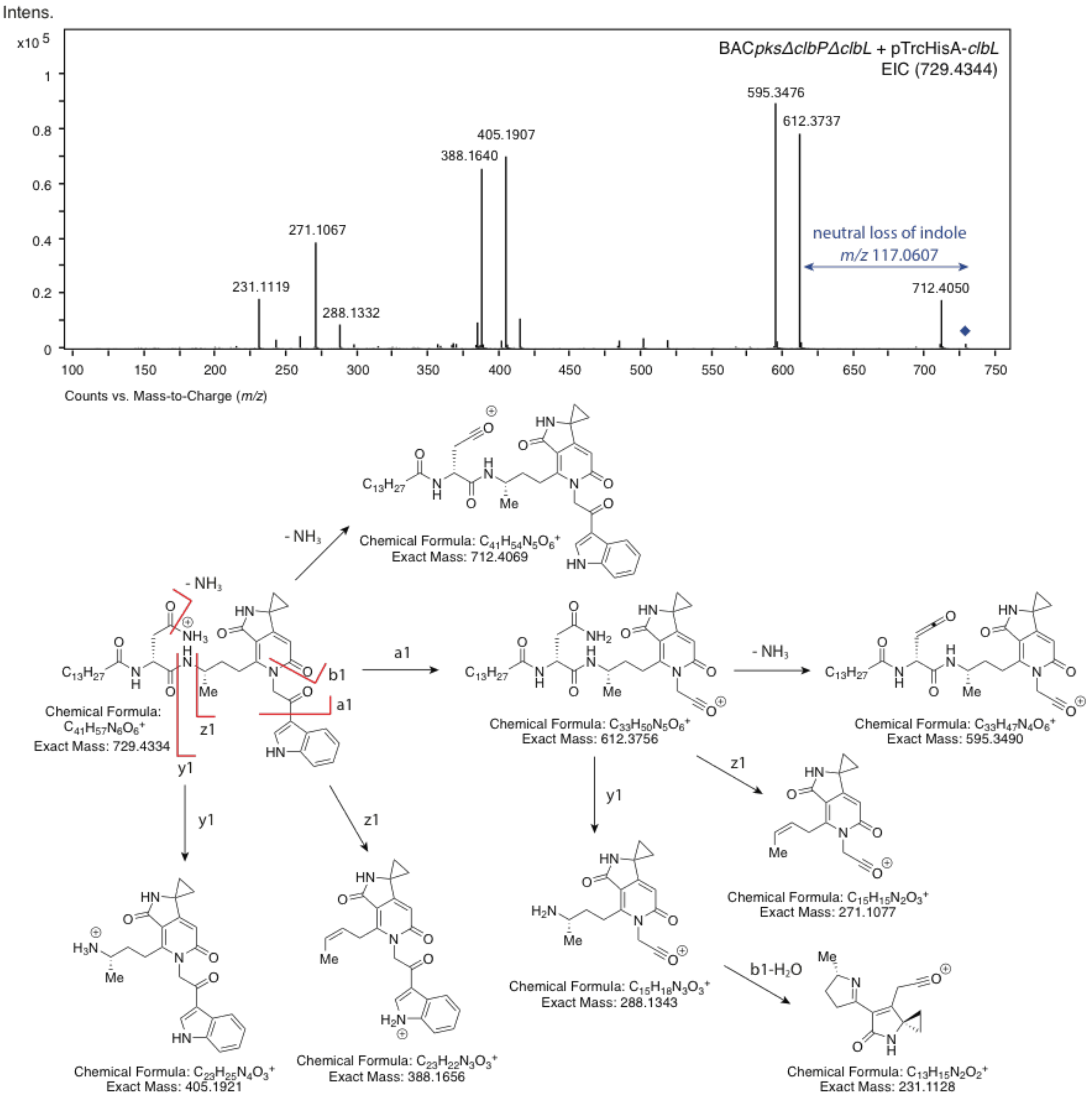


c

| Species | Theoretical *m/z* | Detected *m/z* | Error (ppm) |
|---|---|---|---|
| $C_{41}H_{57}N_6O_6^+$ | 729.4334 | 729.4344 | 1.4 |
| $C_{33}H_{50}N_5O_6^+$ | 612.3756 | 612.3737 | 3.1 |
| $C_{33}H_{47}N_4O_6^+$ | 595.3490 | 595.3476 | 2.4 |
| $C_{23}H_{25}N_4O_3^+$ | 405.1921 | 405.1907 | 3.5 |
| $C_{23}H_{22}N_3O_3^+$ | 388.1656 | 388.1640 | 4.1 |
| $C_{15}H_{18}N_3O_3^+$ | 288.1343 | 288.1332 | 3.8 |
| $C_{15}H_{15}N_2O_3^+$ | 271.1077 | 271.1067 | 3.7 |
| $C_{13}H_{15}N_2O_2^+$ | 231.1128 | 231.1119 | 3.9 |

**Supplementary Figure 9.** (**a**) LC/MS/MS analysis of isolated metabolite **6** (*m/z* 729.4344). (**b**) Predicted major MS/MS fragment ions of **6**. (**c**) Tabulated comparison of theoretical and detected fragment ions of isolated **6**.

**a**

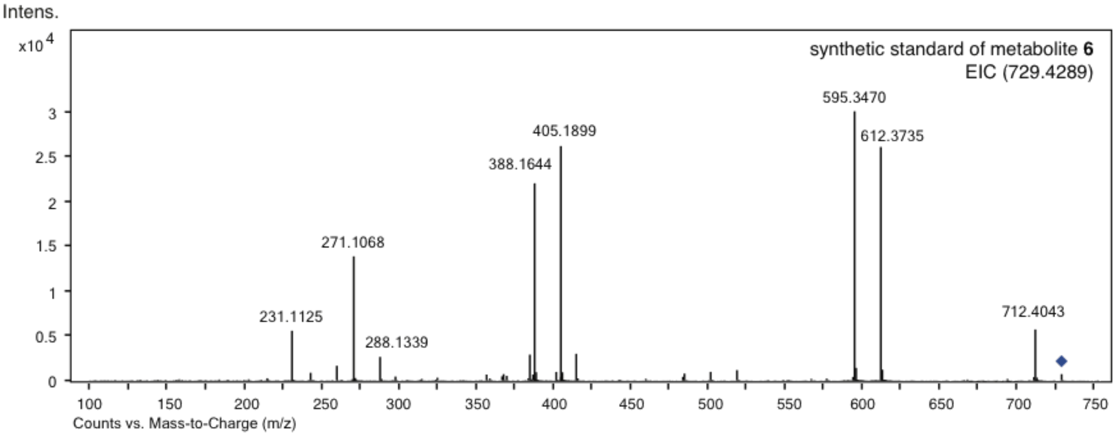


**b**

| Species | Theoretical *m/z* | Detected *m/z* | Error (ppm) |
|---|---|---|---|
| $C_{41}H_{57}N_6O_6^+$ | 729.4334 | 729.4289 | 6.2 |
| $C_{33}H_{50}N_5O_6^+$ | 612.3756 | 612.3735 | 3.4 |
| $C_{33}H_{47}N_4O_6^+$ | 595.3490 | 595.3470 | 3.4 |
| $C_{23}H_{25}N_4O_3^+$ | 405.1921 | 405.1899 | 5.4 |
| $C_{23}H_{22}N_3O_3^+$ | 388.1656 | 388.1644 | 3.1 |
| $C_{15}H_{18}N_3O_3^+$ | 288.1343 | 288.1339 | 1.4 |
| $C_{15}H_{15}N_2O_3^+$ | 271.1077 | 271.1068 | 3.3 |
| $C_{13}H_{15}N_2O_2^+$ | 231.1128 | 231.1125 | 1.3 |

**Supplementary Figure 10.** (**a**) LC/MS/MS analysis of the synthetic standard of **6** (*m/z* 729.4334). (**b**) Tabulated comparison of theoretical and detected fragment ions of the synthetic standard of **6**.

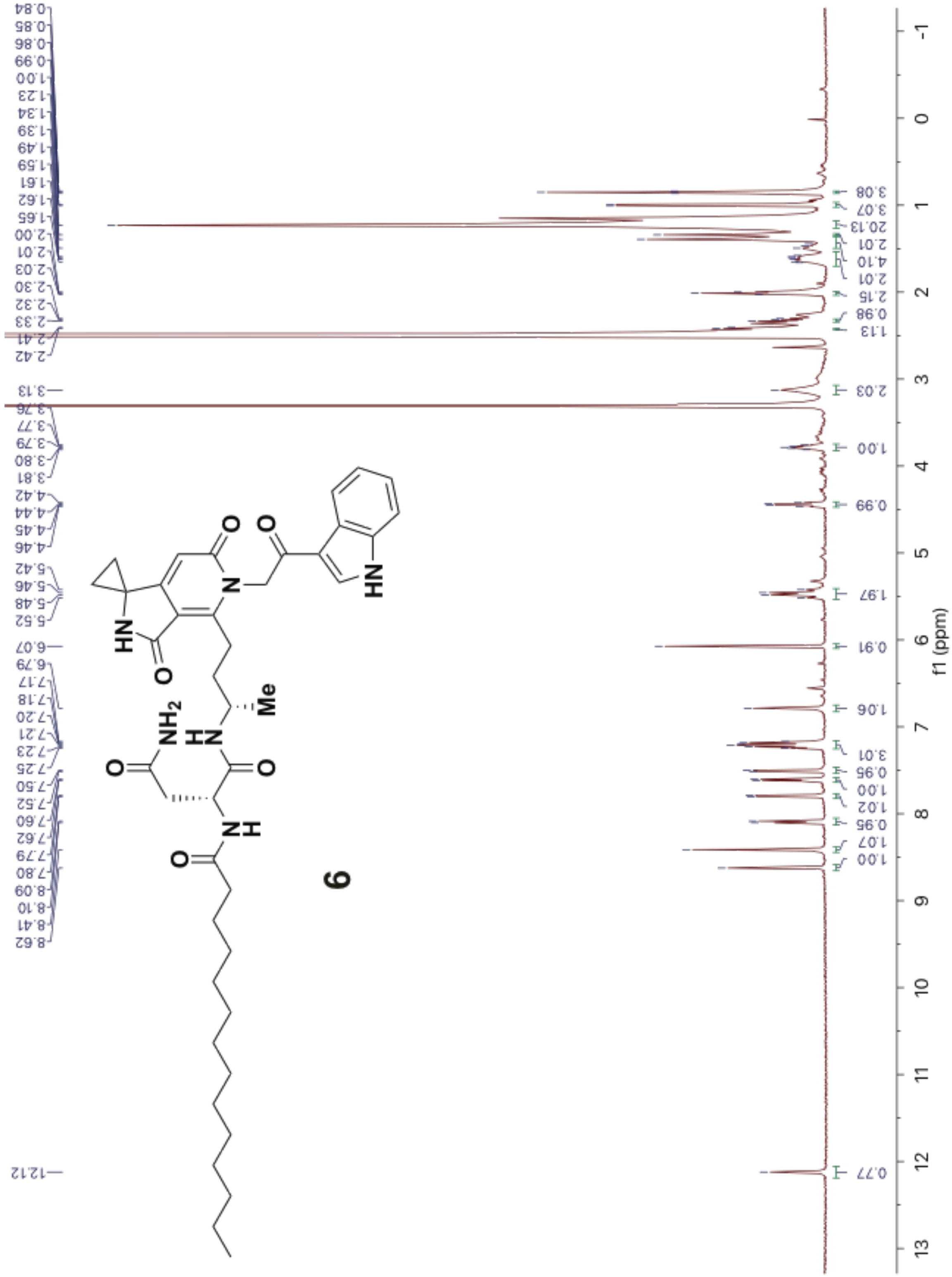

**Supplementary Figure 11.** $^{1}$H-NMR spectrum of the synthetic standard of candidate precolibactin **6** (recorded in DMSO-$d_6$ at 600 MHz).

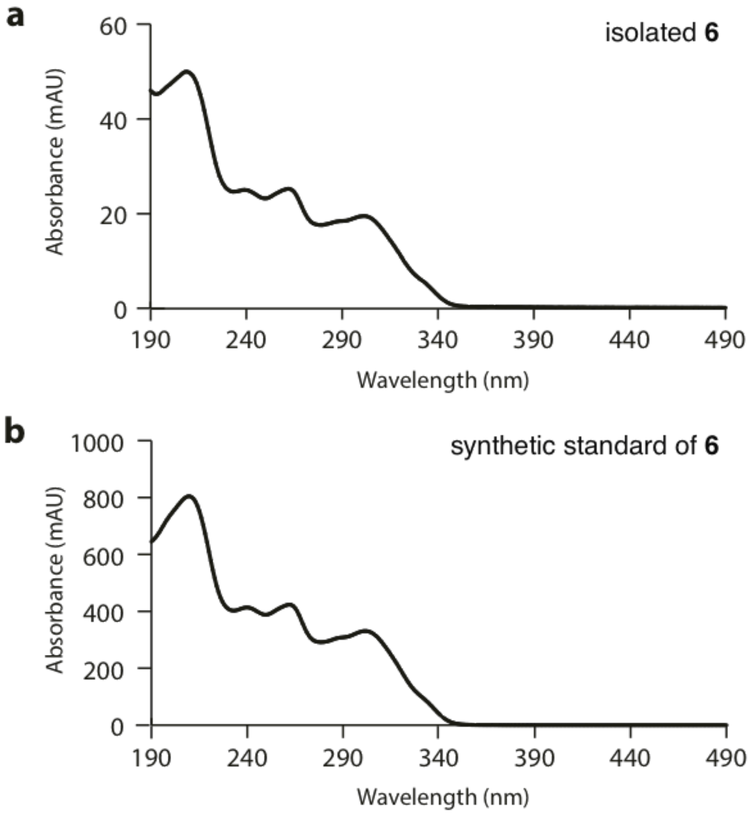


**Supplementary Figure 12.** (**a**-**b**) UV spectrums of candidate precolibactin **6** isolated from bacterial cultures (**a**) and its synthetic standard (**b**).

**a**

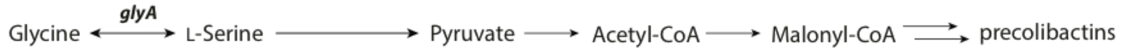


**b**

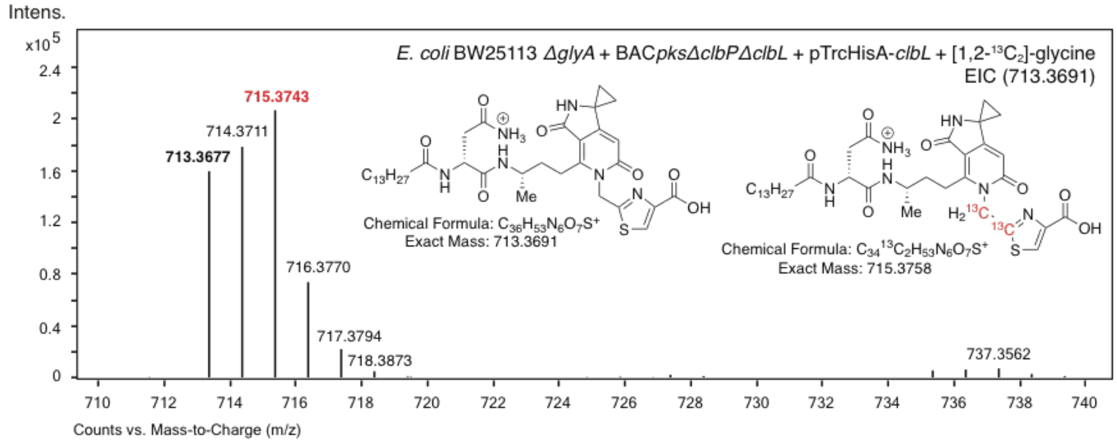


| Species | Theoretical *m/z* | Detected *m/z* | Error (ppm) |
|---|---|---|---|
| $C_{36}H_{53}N_6O_7S^+$ | 713.3691 | 713.3677 | 2.0 |
| $C_{34}{}^{13}C_2H_{53}N_6O_7S^+$ | 715.3758 | 715.3743 | 2.1 |

**c**

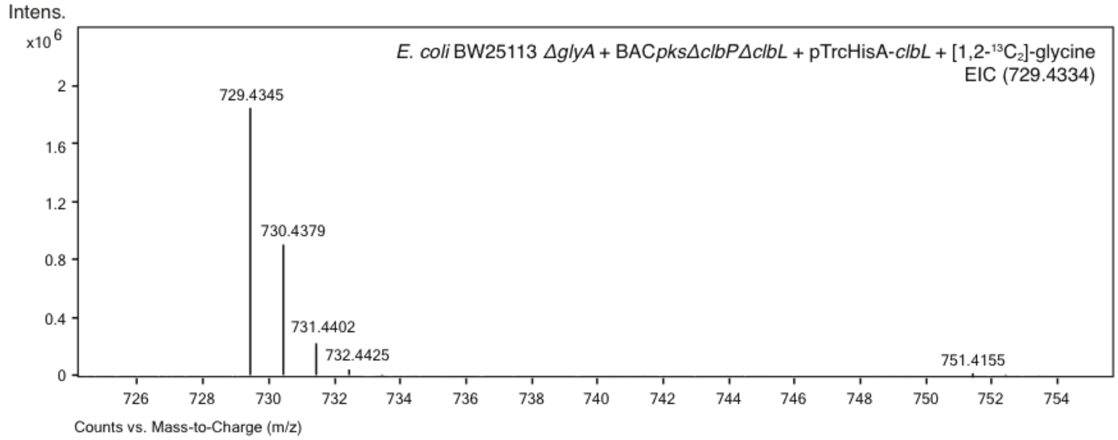


**Supplementary Figure 13.** (**a**) Bacterial conversion of glycine to malonyl-CoA, which could be used to construct the prodrug motif in candidate precolibactins. (**b-c**) EICs of candidate precolibactin **3** (*m/z* 713.3691, **b**) and **6** (*m/z* 729.4334, **c**) in the extracts of a glycine auxotrophic *E. coli* strain expressing BAC*pksΔclbPΔclbL* and pTrcHisA-*clbL*.

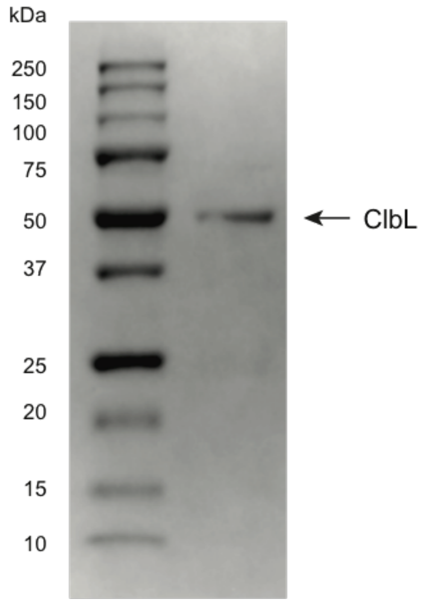


**Supplementary Figure 14.** Coomassie-stained SDS-PAGE gel of purified ClbL.

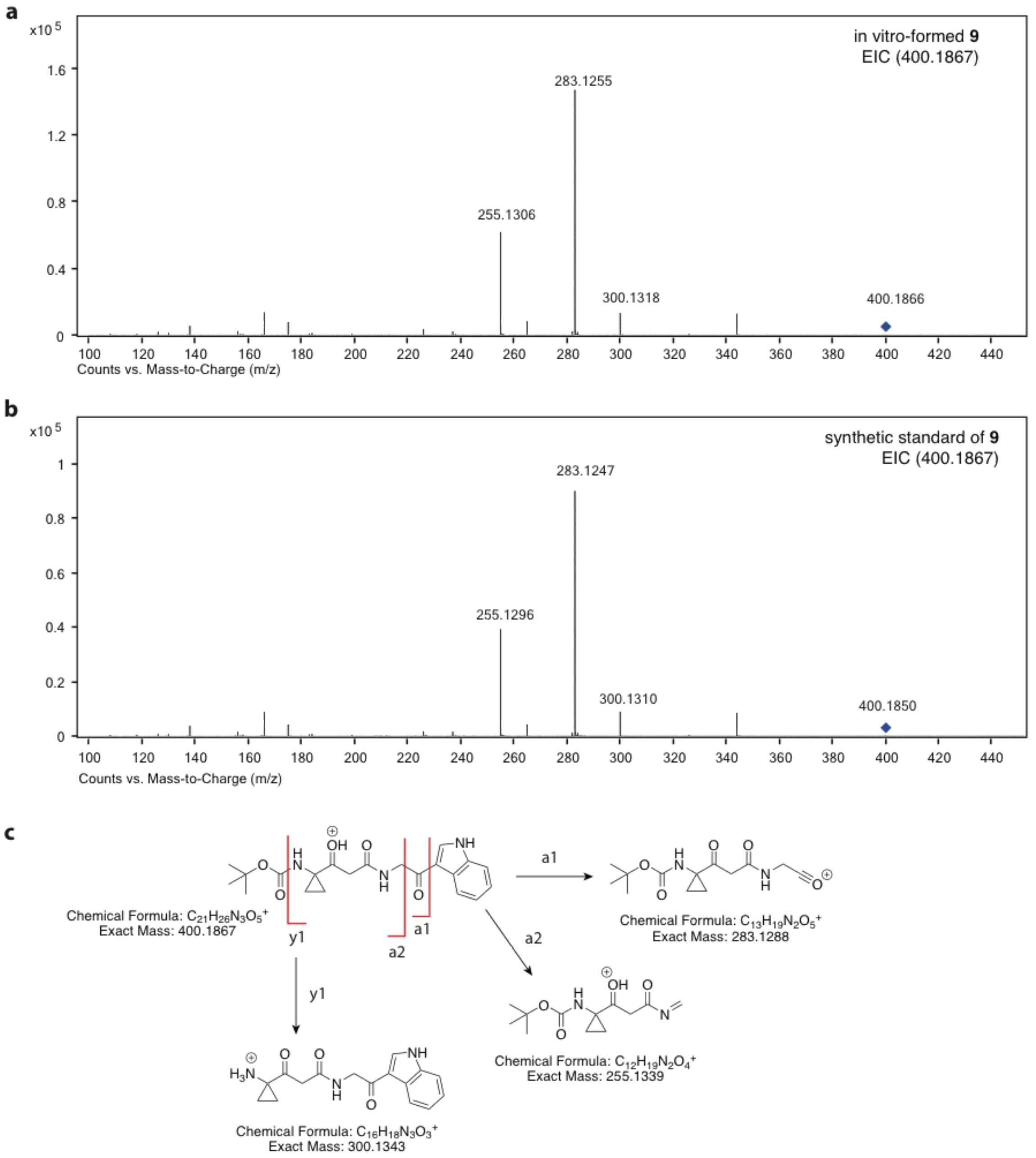


**Supplementary Figure 15.** (**a-b**) LC/MS/MS analysis of in vitro formed product **9** (**a**) and the synthetic standard of **9** (**b**). (**c**) Predicted major MS/MS fragment ions of **9**.

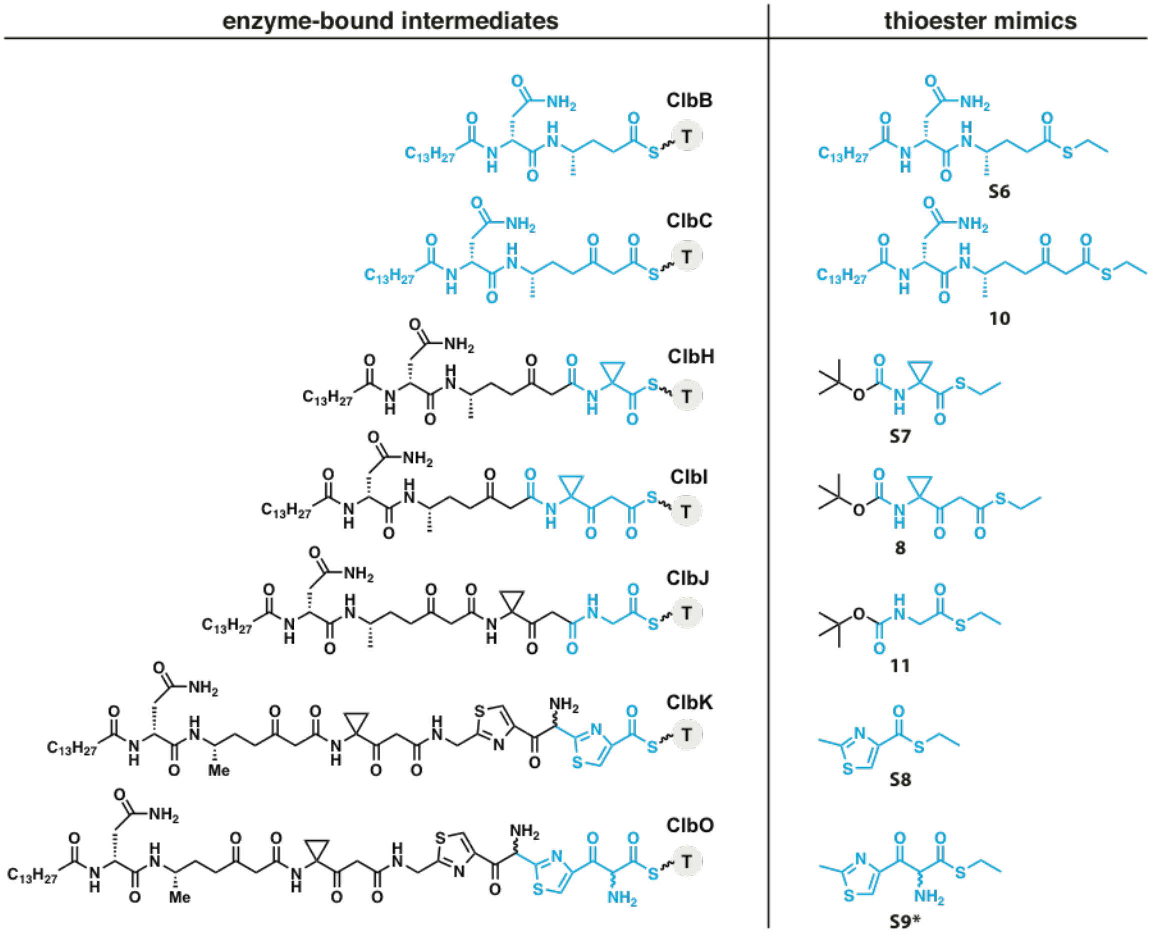


**Supplementary Figure 16.** Thioester mimics synthesized and used in the study. ***S9** was not synthesized due to its intrinsic instability.

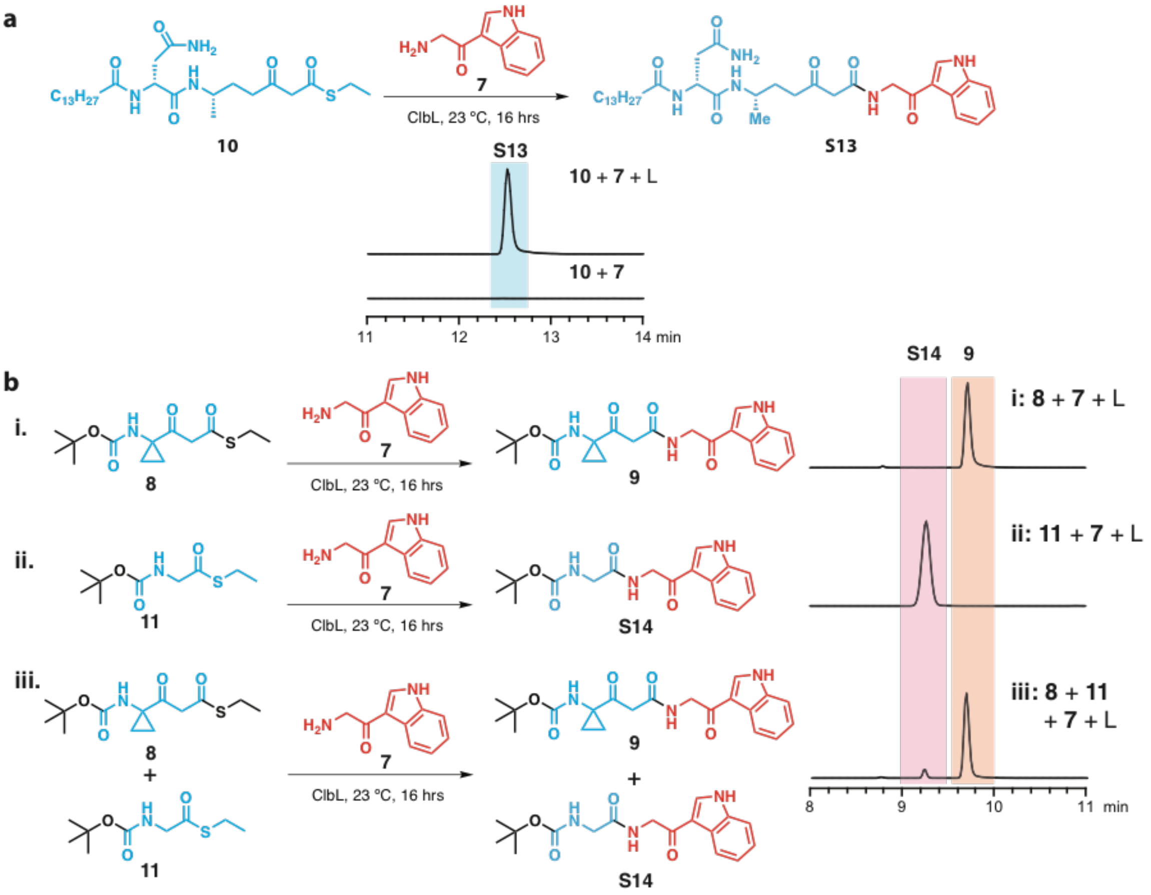


**Supplementary Figure 17.** (**a**) EICs of **S13** (*m/z* 640.4069) formed in an in vitro assay containing ClbL, **10**, and **7**. (**b**) EICs of **9** (*m/z* 400.1867, trace i) and **S14** (*m/z* 332.1605, trace ii) formed in in vitro assays. EICs of **9** and **S14** (trace iii) formed in an in vitro competition assay containing equal molar amounts of **8** and **11**.

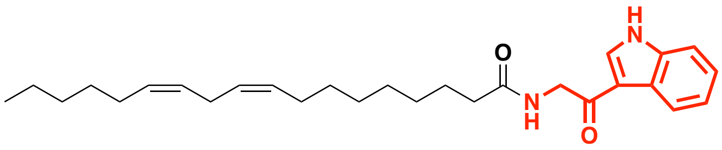

Termitomycamide B

**Supplementary Figure 18.** Chemical structure of Termitomycamide B isolated from *Termitomyces titanicus*.[8] The aminoketone moiety is highlighted in red.

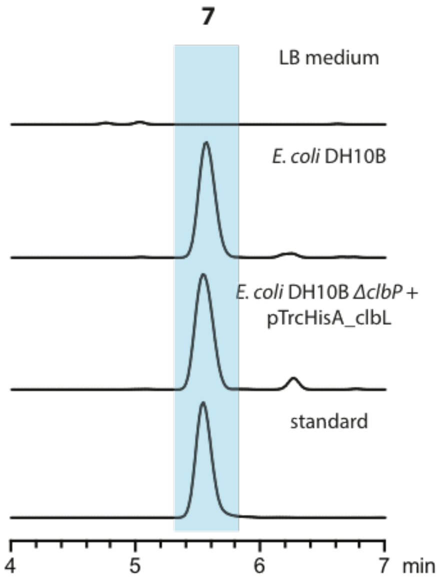


**Supplementary Figure 19.** EICs of **7** (*m/z* 175.0866) in *E. coli* DH10B.

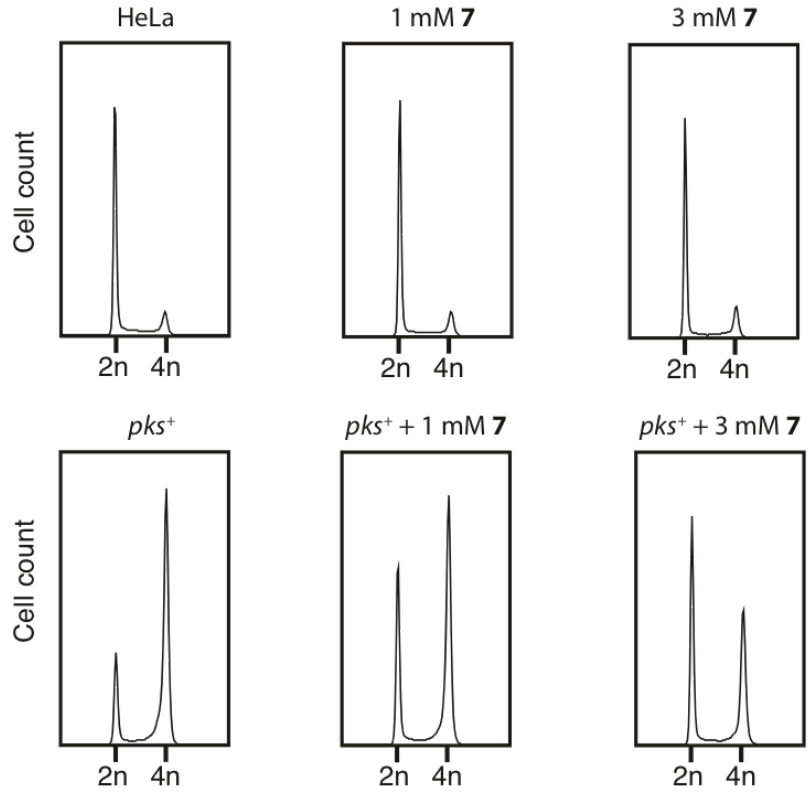


**Supplementary Figure 20.** Cell cycle analysis of HeLa cells treated with *pks*$^+$ *E. coli* and different concentrations of **7**.

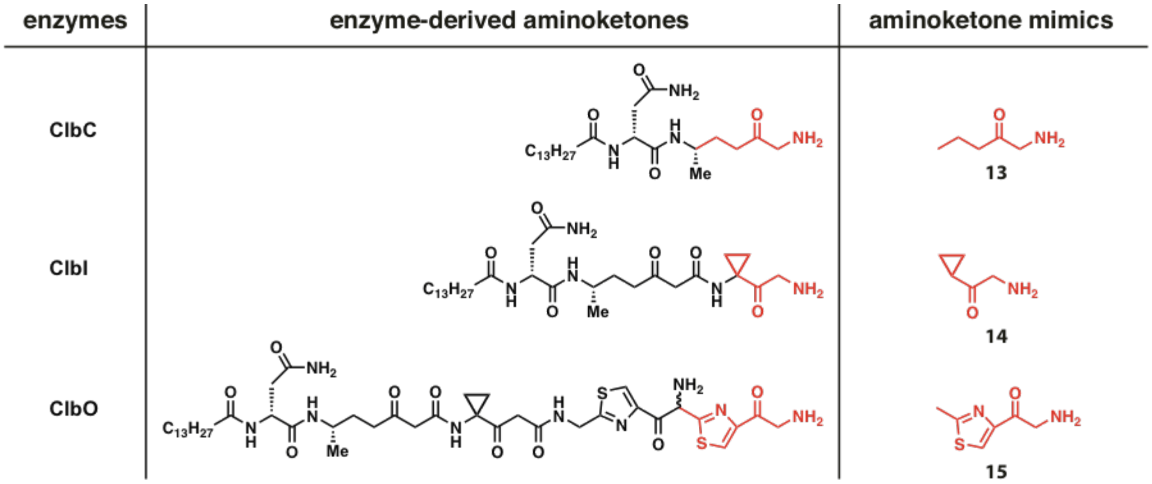


**Supplementary Figure 21.** Aminoketone mimics obtained and used in the study.

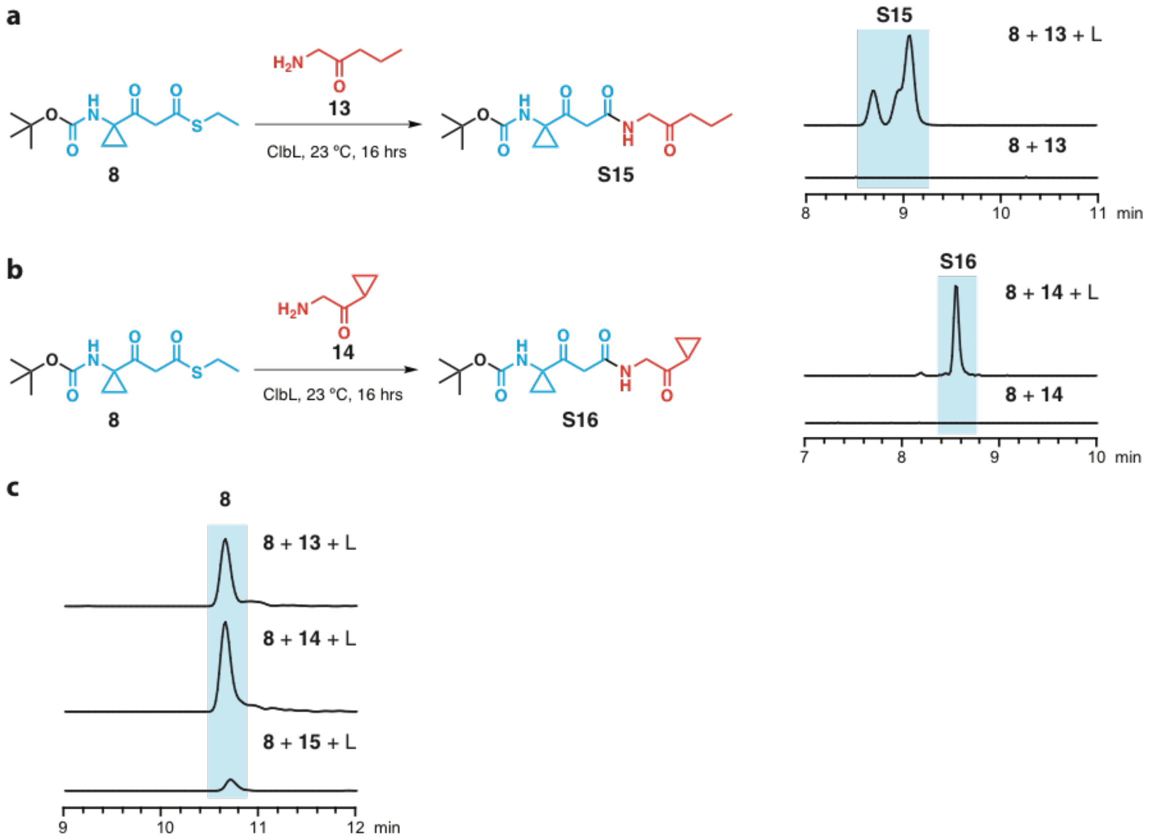


**Supplementary Figure 22.** (**a**) EICs of **S15** (*m/z* 327.1914) formed in an in vitro assay containing ClbL, **8**, and **13**. (**b**) EICs of **S16** (*m/z* 325.1758) formed in an in vitro assay containing ClbL, **8**, and **14**. (**c**) EICs of **8** (*m/z* 288.1264) remained in the assays after 16 h of incubation. Much less **8** remained in the in vitro assay containing **15**, suggesting that ClbL preferentially uses aminoketone **15**.

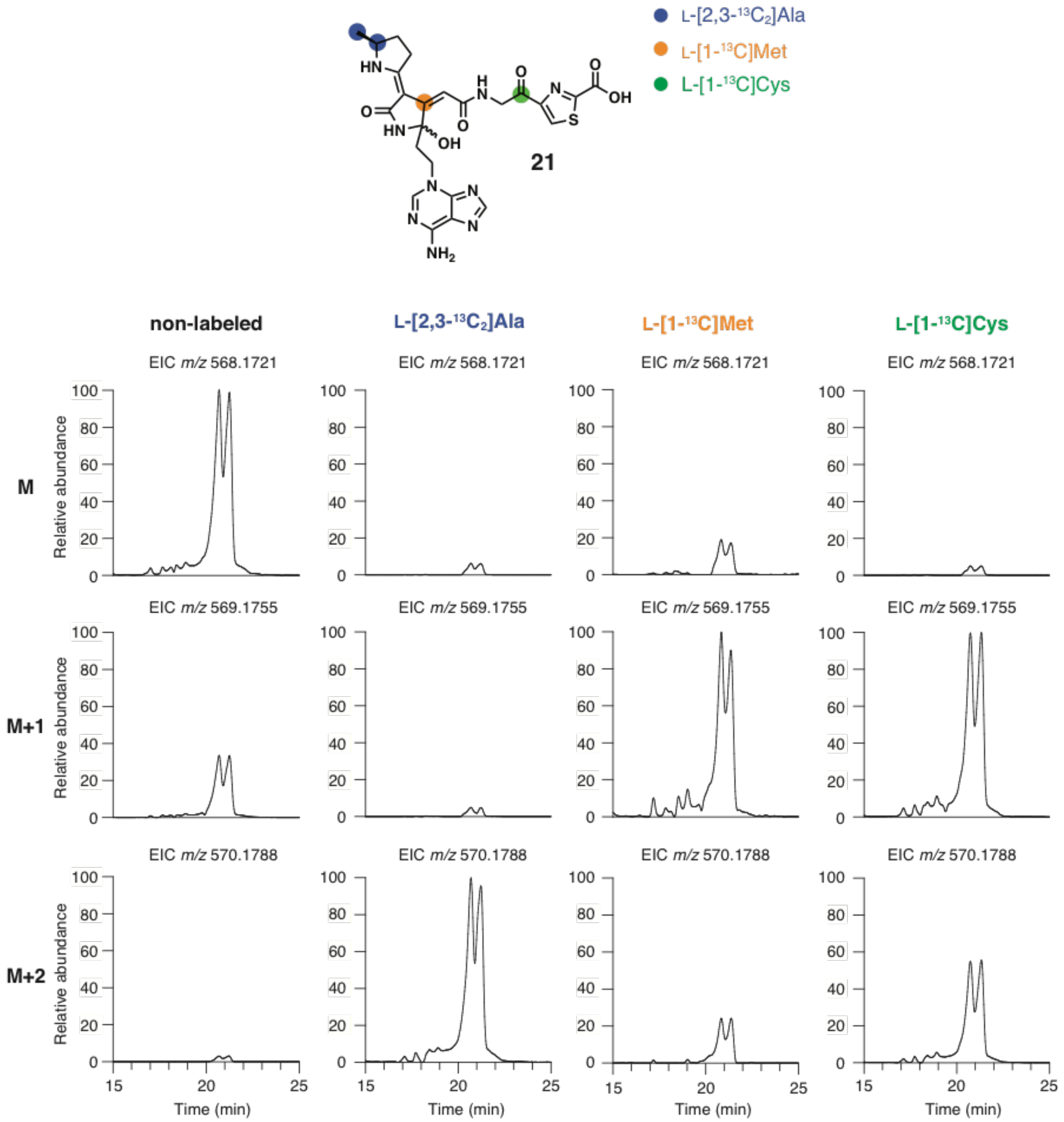


**Supplementary Figure 23.** EICs of *m/z* 568.1721, 569.1755, and 570.1788 for non-labeled, L-[2,3-$^{13}C_2$]Ala, L-[1-$^{13}C$]Met, and L-[1-$^{13}C$]Cys labeled colibactin-derived DNA adduct **21**.

**a**

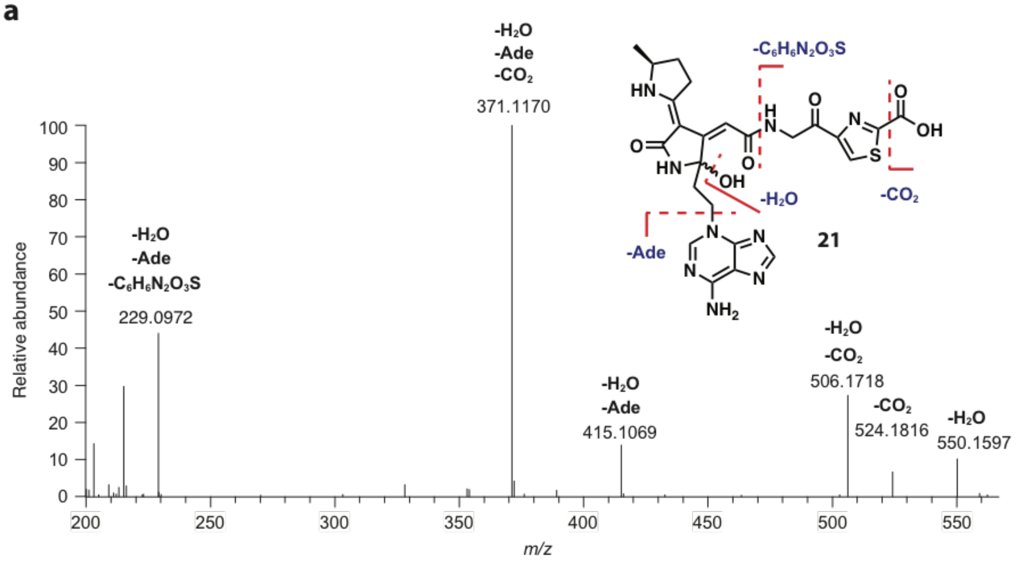


**b**

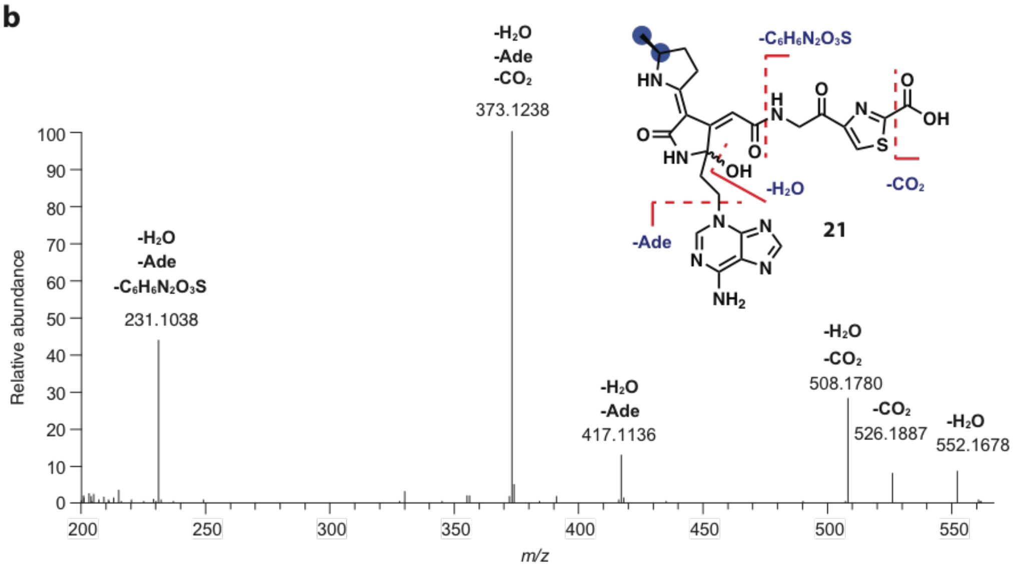


**c**

| Non-labeled feeding | | | | L-[2,3-$^{13}C_2$]Ala feeding | | | |
|---|---|---|---|---|---|---|---|
| Species | Theoretical *m/z* | Detected *m/z* | Error (ppm) | Species | Theoretical *m/z* | Detected *m/z* | Error (ppm) |
| $C_{24}H_{24}N_9O_5S^+$ | 550.1616 | 550.1597 | 3.4 | $C_{22}{}^{13}C_2H_{24}N_9O_5S^+$ | 552.1683 | 552.1678 | 0.9 |
| $C_{23}H_{26}N_9O_4S^+$ | 524.1823 | 524.1816 | 1.3 | $C_{21}{}^{13}C_2H_{26}N_9O_4S^+$ | 526.1890 | 526.1887 | 0.6 |
| $C_{23}H_{24}N_9O_3S^+$ | 506.1717 | 506.1718 | 0.2 | $C_{21}{}^{13}C_2H_{24}N_9O_3S^+$ | 508.1784 | 508.1780 | 0.8 |
| $C_{19}H_{19}N_4O_5S^+$ | 415.1071 | 415.1069 | 0.5 | $C_{17}{}^{13}C_2H_{19}N_4O_5S^+$ | 417.1138 | 417.1136 | 0.5 |
| $C_{18}H_{19}N_4O_3S^+$ | 371.1172 | 371.1170 | 0.5 | $C_{16}{}^{13}C_2H_{19}N_4O_3S^+$ | 373.1239 | 373.1238 | 0.3 |
| $C_{13}H_{13}N_2O_2^+$ | 229.0972 | 229.0972 | 0.0 | $C_{11}{}^{13}C_2H_{13}N_2O_2^+$ | 231.1039 | 231.1038 | 0.4 |

**Supplementary Figure 24.** (**a**) LC/MS/MS analysis of non-labeled colibactin-derived DNA adduct **21** (*m/z* 568.1721). (**b**) LC/MS/MS analysis of L-[2,3-$^{13}C_2$]Ala labeled **21** (*m/z* 570.1788). (**c**) Tabulated comparison of theoretical and detected fragment ions of non-labeled and L-[2,3-$^{13}C_2$]Ala labeled **21**.

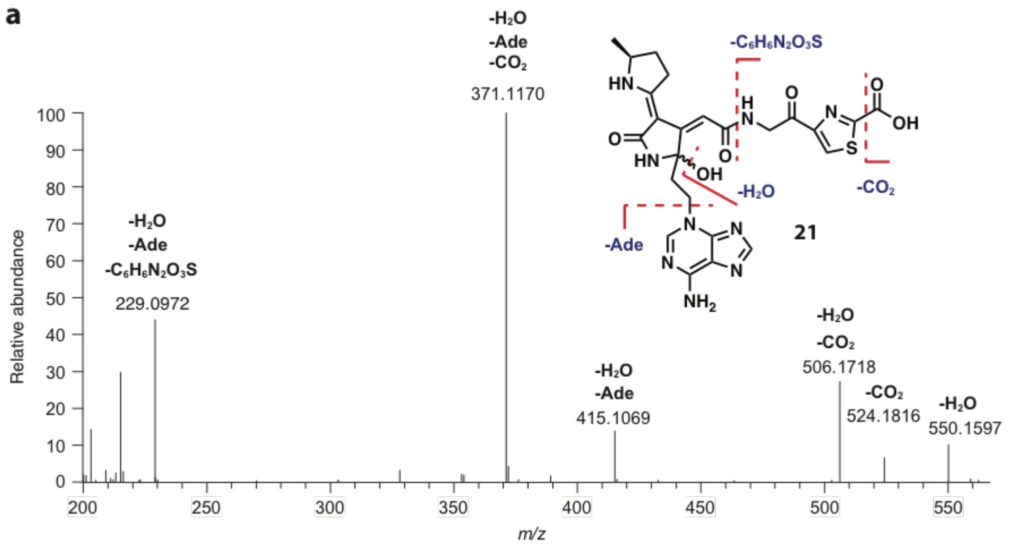


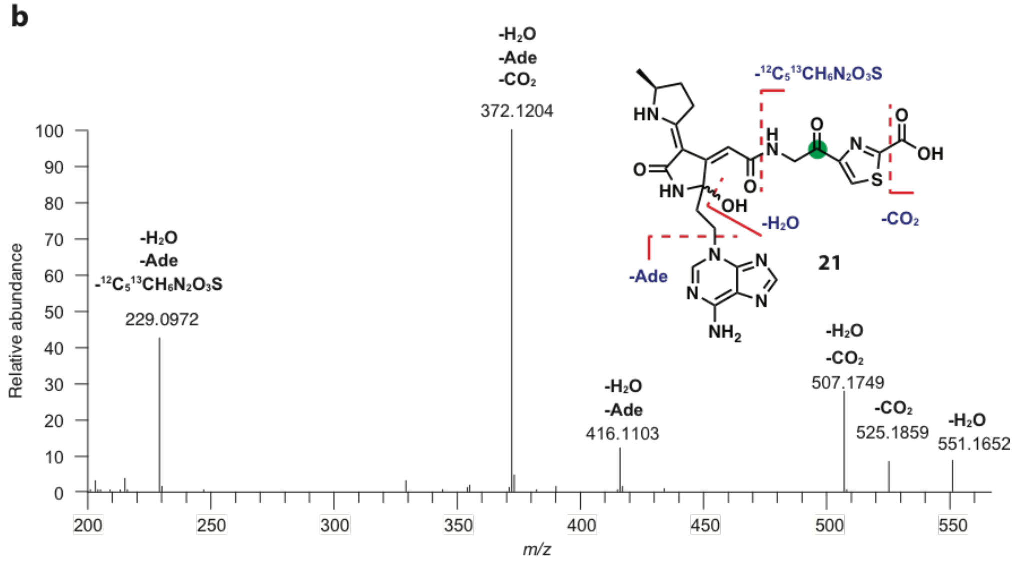


**c**

| Non-labeled feeding | | | | L-[1-$^{13}$C]Cys feeding | | | |
|---|---|---|---|---|---|---|---|
| Species | Theoretical *m/z* | Detected *m/z* | Error (ppm) | Species | Theoretical *m/z* | Detected *m/z* | Error (ppm) |
| $C_{24}H_{24}N_9O_5S^+$ | 550.1616 | 550.1597 | 3.4 | $C_{23}{}^{13}CH_{24}N_9O_5S^+$ | 551.1649 | 551.1652 | 0.5 |
| $C_{23}H_{26}N_9O_4S^+$ | 524.1823 | 524.1816 | 1.3 | $C_{22}{}^{13}CH_{26}N_9O_4S^+$ | 525.1857 | 525.1859 | 0.4 |
| $C_{23}H_{24}N_9O_3S^+$ | 506.1717 | 506.1718 | 0.2 | $C_{22}{}^{13}CH_{24}N_9O_3S^+$ | 507.1751 | 507.1749 | 0.4 |
| $C_{19}H_{19}N_4O_5S^+$ | 415.1071 | 415.1069 | 0.5 | $C_{18}{}^{13}CH_{19}N_4O_5S^+$ | 416.1104 | 416.1103 | 0.2 |
| $C_{18}H_{19}N_4O_3S^+$ | 371.1172 | 371.1170 | 0.5 | $C_{17}{}^{13}CH_{19}N_4O_3S^+$ | 372.1206 | 372.1204 | 0.5 |
| $C_{13}H_{13}N_2O_2^+$ | 229.0972 | 229.0972 | 0.0 | $C_{13}H_{13}N_2O_2^+$ | 229.0972 | 229.0972 | 0.0 |

**Supplementary Figure 25.** (**a**) LC/MS/MS analysis of non-labeled colibactin-derived DNA adduct **21** (*m/z* 568.1721). (**b**) LC/MS/MS analysis of L-[1-$^{13}$C]Cys labeled **21** (*m/z* 569.1755). (**c**) Tabulated comparison of theoretical and detected fragment ions of non-labeled and L-[1-$^{13}$C]Cys labeled **21**.

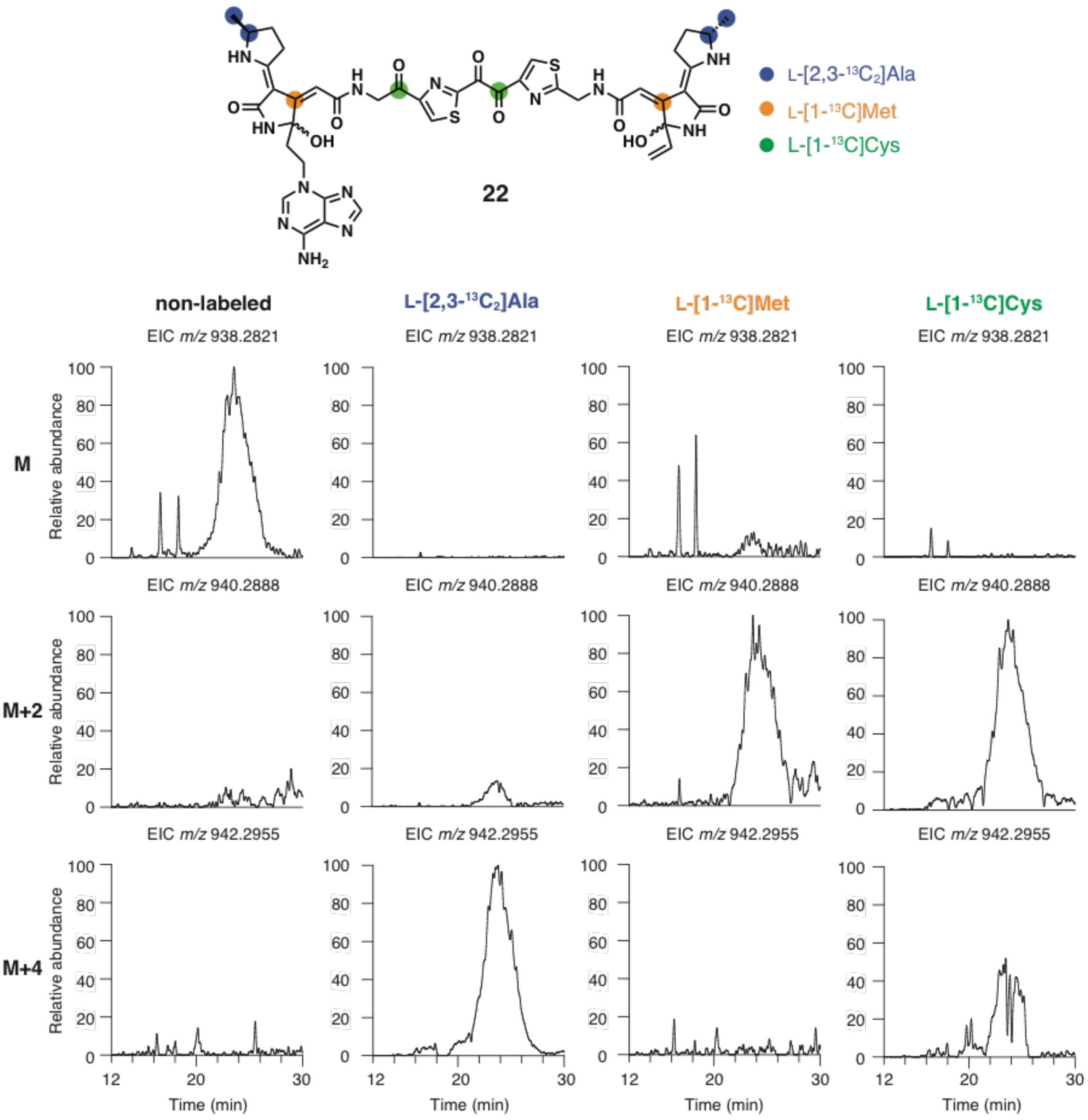


**Supplementary Figure 26.** EICs of $m/z$ 938.2821, 940.2888, and 942.2955 for non-labeled, L-[2,3-$^{13}C_2$]Ala, L-[1-$^{13}$C]Met, and L-[1-$^{13}$C]Cys labeled colibactin-derived DNA adduct **22**.

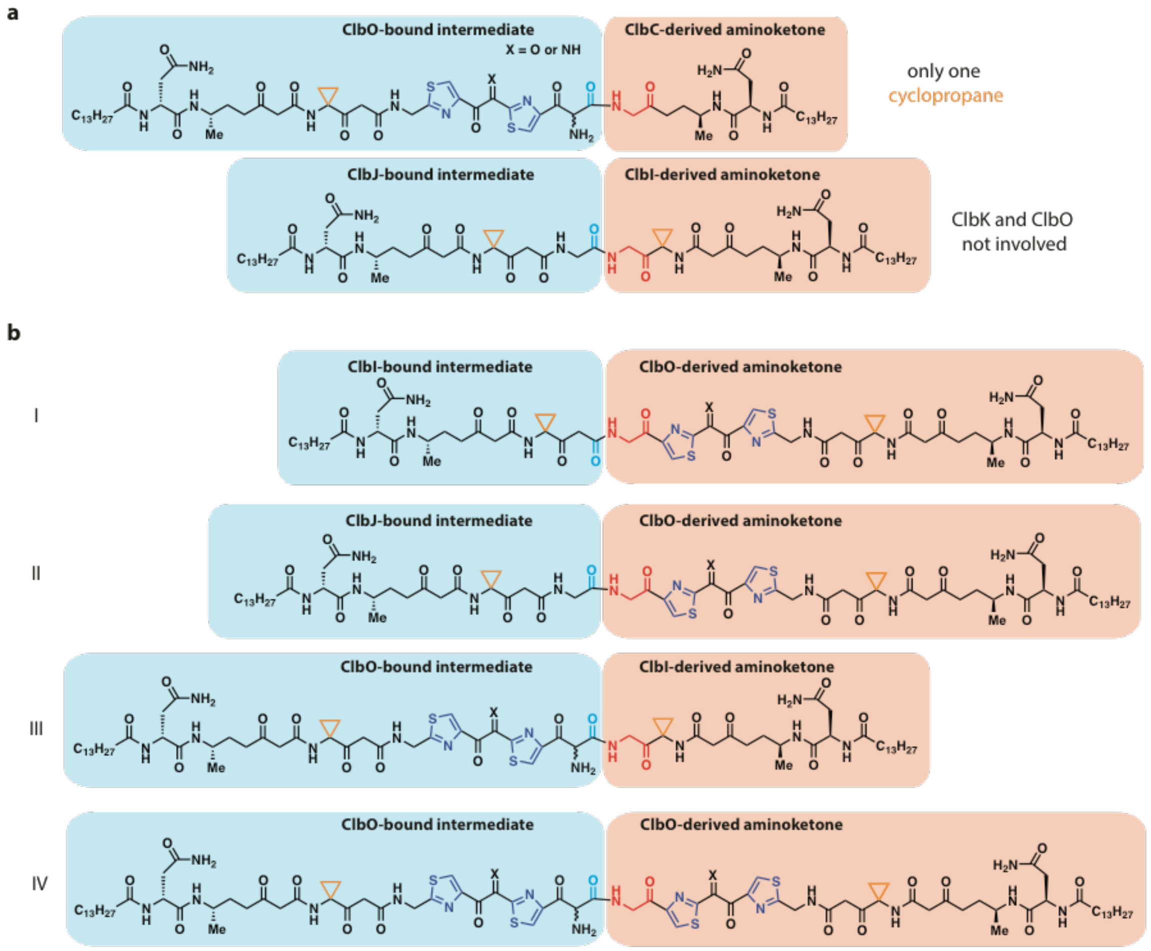


**Supplementary Figure 27.** (**a**) Examples of combinations that violate the two criteria for the structure of the active colibactin genotoxin. (**b**) Four possible structures of precolibactin compatible with these criteria.

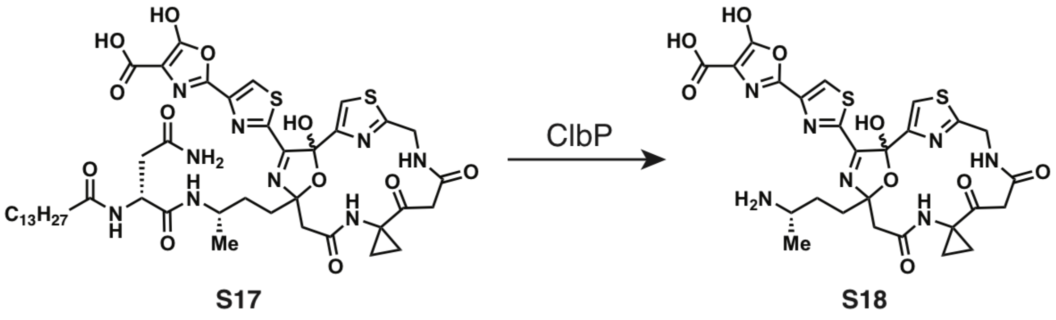


**Supplementary Figure 28.** Proposed precolibactin **S17** and colibactin **S18** by Qian and Zhang.

### 4. Synthetic procedures

General Procedure A: preparation of thioesters

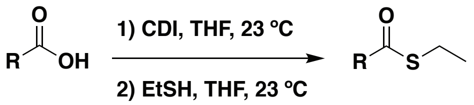


To an oven-dried round bottom flask equipped with a stir bar was added the appropriate carboxylic acid (1.0 equiv) and anhydrous THF (0.5 M) under dry $N_2$ at 23 ºC. 1,1'-Carbonyldiimidazole (CDI, 1.5 equiv) was added and the reaction mixture was stirred at 23 ºC until TLC indicated the reaction was complete (approximately 4 h). Once the reaction was complete, ethanethiol (2.0 equiv) was added and the reaction mixture was stirred at 23 ºC overnight. The reaction mixture was then diluted with 5 reaction volumes of ethyl acetate (EtOAc) and 1 reaction volume of $H_2O$. The layers were separated and the aqueous layer was extracted 2 more times with EtOAc. The combined organic layers were washed with brine, dried over $Na_2SO_4$, filtered, and concentrated *in vacuo*. The resulting residue was purified by flash column chromatography (0% EtOAc/Hexane to 50% EtOAc/Hexane) on silica gel.

Preparation of β-keto amide **9**

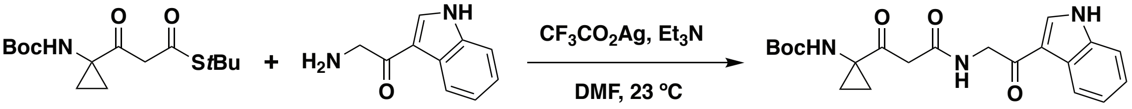


S2 7 9

To an oven-dried round bottom flask equipped with a stir bar was added known β-keto thioester **S2** (104 mg, 0.33 mmol, 1.2 equiv)[9] and anhydrous DMF (1.4 mL, 0.2 M) followed by 2-amino-1-(1*H*-indol-3-yl)ethan-1-one hydrochloride **7** (47 mg, 0.27 mmol, 1 equiv) and triethylamine (0.15 mL, 1.08 mmol, 4 equiv) under dry $N_2$. To the resulting solution was added silver trifluoroacetate (72 mg, 0.33 mmol, 1.2 equiv) in three equal amounts every 20 min (total of 1 h). The reaction mixture was then stirred at 23 ºC for an additional 1 h (total reaction time of 2 h). The reaction mixture was diluted with 5% aqueous citric acid solution (10 mL) and the precipitate isolated by filtration. The resulting crude solid was dissolved in DMSO (5 mL) and purified by reverse-phase preparative HPLC at a flow rate of 8 mL/min using 0.1% formic acid in water as mobile phase A and 0.1% formic acid in acetonitrile as mobile phase B. The following gradient was applied: 0-1 min, 40% isocratic; 1-15 min, 40-70% B; 15-18 min, 70-100% B; 18-20 min, 100% B isocratic; 20-22 min, 100-40% B; 22-26 min, 40% isocratic. The product containing fractions were pooled and concentrated to afford **9** (58 mg, 53% yield) as a white solid. $^1H$ NMR (500 MHz, DMSO-$d_6$): δ (ppm) = 12.04 (s, 1H), 8.43 (s, 1H), 8.34 (t, $J$ = 5.1 Hz, 1H), 8.16 (d, $J$ = 7.6 Hz, 1H), 7.75 (s, 1H), 7.49 (d, $J$ = 8.1 Hz, 1H), 7.25 - 7.17 (m, 2H), 4.49 (d, $J$ = 5.2 Hz, 2H), 3.59 (s, 2H), 1.42 (s, 9H), 1.40 - 1.35 (m, 2H), 1.09 - 1.05 (m, 2H). $^{13}C$ NMR (126 MHz, DMSO-$d_6$): δ (ppm) = 205.0, 189.9, 166.3, 156.0, 136.4, 133.8, 125.4, 122.9, 121.8, 121.1, 113.9, 112.2, 78.6, 45.9, 41.2, 28.2, 19.4. HRMS (ESI): calcd for $C_{21}H_{26}N_3O_5^+$ $[M+H]^+$, 400.1867; found, 400.1876.

Preparation of synthetic standard of **6**

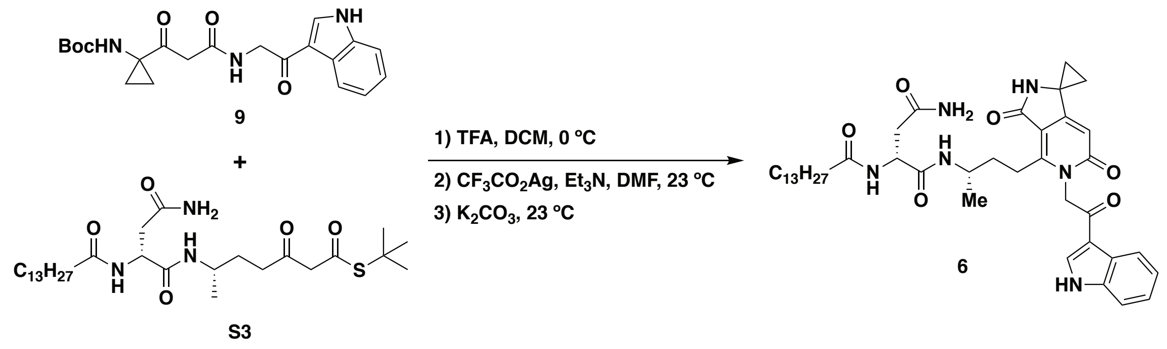


To an oven-dried round bottom flask equipped with a stir bar was added Boc-protected β-keto amide **9** (12 mg, 0.030 mmol, 1 equiv) and dry $CH_2Cl_2$ (0.2 mL, 0.15 M). The solution was cooled to 0 ºC and trifluoroacetic acid (0.2 mL, 0.15 M) was added dropwise. The reaction mixture was stirred at 0 ºC until TLC indicated the reaction was complete (approximately 2 h). Once the reaction was complete, dry toluene (1 mL) was added to the reaction mixture. The resulting solution was then concentrated *in vacuo* to give a light brown oil. The crude TFA salt was used in the next step without further purification.

Dry DMF (1.0 mL, 0.03 M) was added to the crude TFA salt followed by the known β-keto thioester **S3** (14 mg, 0.025 mmol, 0.8 equiv),[9] triethylamine (0.017 mL, 0.12 mmol, 4 equiv), and silver trifluoroacetate (12 mg, 0.054 mmol, 1.8 equiv). The reaction mixture was stirred at 23 ºC for 2 h. After 2 h, the reaction mixture was diluted with 5% aqueous citric acid solution (10 mL) and the precipitate isolated by filtration. The resulting crude solid was dissolved in DMSO (3 mL) and taken to the next step without further purification.

To a small vial equipped with a stir bar was added the coupled product in DMSO (2 mL) followed by MeOH (0.5 mL, 0.04 M) and potassium carbonate (10 mg, 0.06 mmol, 3.5 equiv) at 23 ºC. The reaction mixture was stirred 16 h at 23 ºC. After 16 h, the reaction mixture was transferred to an Eppendorf tube and centrifuged at 23 ºC (13, 200 rpm × 2 min). The resulting supernatant was purified by reverse-phase semi-preparative HPLC using a Hypersil Gold™ aQ C18 polar endcapped column (250 × 10 mm, 5 μm particle size, Thermo Fisher Scientific). The following gradient was applied: 0-15 min, 65-68% B; 15-20 min, 68-75% B; 20-20.5 min, 75-100% B; 20.5-23 min, 100% B isocratic; 23-24 min, 100-65% B; 24-30 min, 65% B isocratic (solvent A: water + 0.1% formic acid; solvent B: acetonitrile + 0.1% formic acid; flow rate: 3 mL/min). The product containing fractions were pooled and concentrated to afford **6** (4.0 mg, 22% yield over three steps) as a white solid. $^1H$ NMR (600 MHz, DMSO-$d_6$): δ (ppm) = 12.12 (br s, 1H), 8.62 (s, 1H), 8.41 (s, 1H), 8.10 (d, $J$ = 7.7 Hz, 1H), 7.80 (d, $J$ = 7.9 Hz, 1H), 7.61 (d, $J$ = 7.9 Hz, 1H), 7.51 (d, $J$ = 8.0 Hz, 1H), 7.25 - 7.16 (m, 3H), 6.79 (s, 1H), 6.07 (s, 1H), 5.53 - 5.41 (m, 2H), 4.44 (q, $J$ = 7.1 Hz, 1H), 3.82 - 3.75 (m, 1H), 3.18 - 3.07 (m, 2H), 2.44 - 2.40 (m, 1H), 2.35 - 2.30 (m, 1H), 2.01 (t, $J$ = 7.1 Hz, 2H), 1.70 - 1.54 (m, 2H), 1.50 - 1.38 (m, 4H), 1.36 - 1.32 (m, 2H), 1.27 - 1.18 (m, 20H), 1.00 (d, $J$ = 6.4 Hz, 3H), 0.85 (t, $J$ = 6.7 Hz, 3H). $^{13}C$ NMR data were not collected due to poor solubility of **6** in DMSO. HRMS (ESI): calcd for $C_{41}H_{57}N_6O_6^+$ $[M+H]^+$, 729.4334; found, 729.4289.

Preparation of β-keto thioester **8**

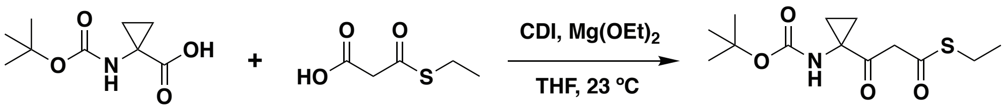


To an oven-dried round bottom flask equipped with a stir bar was added *N*-(*tert*-butoxycarbonyl)-1-amino-1-cyclopropane carboxylic acid **S4** (114 mg, 0.57 mmol, 1.0 equiv) and anhydrous THF (2.8 mL, 0.2 M) under dry $N_2$ at 23 ºC. CDI (138 mg, 0.85 mmol, 1.5 equiv) was added and the reaction mixture was stirred for 6 h at 23 ºC. In another oven-dried round bottom flask equipped with a stir bar, magnesium ethoxide (65 mg, 0.57 mmol, 1 equiv) was added to a solution of 3-(ethylthio)-3-oxopropanoic acid (**S5**, 168 mg, 1.13 mmol, 2 equiv) in THF (2.3 mL, 0.25 M). The reaction mixture was stirred for 6 h at 23 ºC, and then concentrated *in vacuo*. The resulting magnesium salt of **S5** was dissolved in anhydrous THF (0.5 mL) and transferred to the flask containing the acylimidazole derivative of **S4** via syringe. The reaction mixture was stirred overnight at 23 ºC. EtOAc (20 mL) and $H_2O$ (20 mL) were added to the reaction mixture and the layers were then separated. The aqueous layer was extracted with EtOAc (2 × 20 mL). The combined organic layers were washed with saturated aqueous NaCl solution (40 mL), dried with $Na_2SO_4$, filtered, and concentrated *in vacuo*. The resulting residue was purified by flash column chromatography (0% EtOAc/Hexane to 20% EtOAc/Hexane) on silica gel to afford compound **8** as a colorless oil (78 mg, 48% yield). $^1$H NMR (600 MHz, $CDCl_3$): δ (ppm) = 5.34 (s, 1H), 3.87 (s, 2H), 2.91 (q, *J* = 7.5 Hz, 2H), 1.61 (q, *J* = 4.6 Hz, 2H), 1.45 (s, 9H), 1.26 (t, *J* = 7.5 Hz, 3H), 1.20 – 1.15 (m, 2H). $^{13}$C NMR (126 MHz, $CDCl_3$): δ (ppm) = 203.2, 193.0, 155.7, 80.5, 54.3, 41.6, 28.3, 23.9, 21.7, 14.5. HRMS (ESI): calcd for $C_{13}H_{22}NO_4S$ $[M+H]^+$, 288.1264; found, 288.1277.

Preparation of thioester **S6**

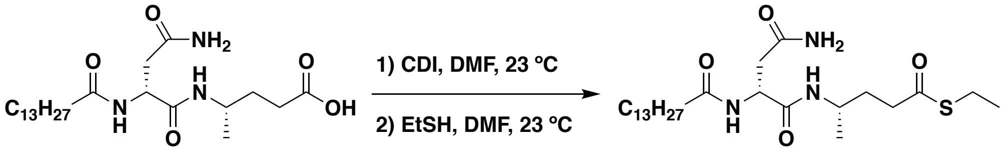


To an oven-dried round bottom flask equipped with a stir bar was added known carboxylic acid **S10** (70 mg, 0.16 mmol, 1.0 equiv)[9] and anhydrous DMF (0.8 mL, 0.2 M) under dry $N_2$ at 23 ºC. CDI (39 mg, 0.24 mmol, 1.5 equiv) was added and the reaction mixture was stirred for 6 h at 23 ºC. After 6 h, ethanethiol (0.024 mL, 0.32 mmol, 2.0 equiv) was added and the reaction mixture was stirred at 23 ºC overnight. The reaction mixture was then diluted with 1M aqueous HCl (10 mL) and the resulting precipitate isolated by filtration. The crude solid was then dissolved in DMSO (2 mL) and purified by reverse-phase preparative HPLC using a Hypersil Gold™ aQ C18 polar endcapped column (250 × 20 mm, 5 μm particle size, Thermo Fisher Scientific). The following gradient was applied: 0-1 min, 50% B isocratic; 1-20 min, 50-100% B; 20-23 min, 100% B isocratic; 23-24 min, 100-50% B; 24-30 min, 50% B isocratic (solvent A: water + 0.1% formic acid; solvent B:

acetonitrile + 0.1% formic acid; flow rate: 8 mL/min). The product containing fractions were pooled and concentrated to afford **S6** (32 mg, 42% yield) as a white solid. $^{1}$H NMR (600 MHz, DMSO-$d_6$): δ (ppm) = 7.90 (d, *J* = 7.9 Hz, 1H), 7.52 (d, *J* = 8.4 Hz, 1H), 7.23 (br s, 1H), 6.82 (br s, 1H), 4.44 (q, *J* = 7.3 Hz, 1H), 3.78 – 3.68 (m, 1H), 2.80 (q, *J* = 7.4 Hz, 2H), 2.50 - 2.44 (m, 2H), 2.44 (dd, *J* = 15.1, 6.1Hz, 1H), 2.32 (dd, *J* = 15.1, 7.6 Hz, 1H), 2.08 (t, *J* = 7.3 Hz, 2H), 1.71 - 1.57 (m, 1H), 1.50 - 1.42 (m, 2H), 1.23 (s, 20H), 1.15 (t, *J* = 7.4 Hz, 3H), 1.00 (d, *J* = 6.6 Hz, 3H), 0.85 (t, *J* = 6.7 Hz, 3H). $^{13}$C NMR (126 MHz, DMSO-$d_6$)*: δ (ppm) = 198.6, 172.1, 171.4, 170.4, 49.8, 43.7, 40.4, 37.2, 35.2, 31.4, 31.3, 29.04, 29.00, 28.94, 28.86, 28.70, 28.62, 25.2, 22.5, 22.1, 20.4, 14.7, 13.9. HRMS (ESI): calcd for $C_{25}H_{48}N_3O_4S$ $[M+H]^+$, 486.3360; found, 486.3358. $[\alpha]_D^{23}$ = + 9.4° (*c* = 1.0, DMSO-$d_6$).

*23 distinct $^{13}$C signals are observed for this substrate (25 are expected), which is likely due to the close overlap of peaks from the myristoyl group in the 29.1 - 28.5 ppm range.

Preparation of β-keto thioester **10**

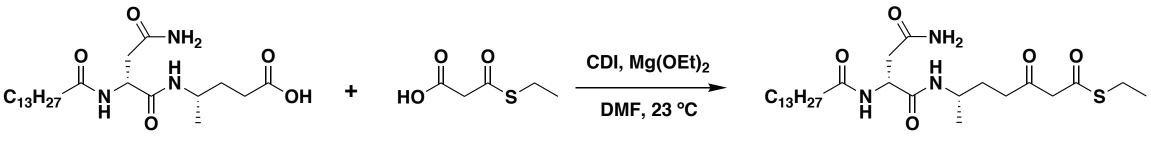


To an oven-dried round bottom flask equipped with a stir bar was added the known carboxylic acid **S10** (44 mg, 0.10 mmol, 1.0 equiv)[9] and anhydrous DMF (1.0 mL, 0.1 M) under dry $N_2$ at 23 ºC. CDI (24 mg, 0.15 mmol, 1.5 equiv) was added and the reaction mixture was stirred for 6 h at 23 ºC. In another oven-dried round bottom flask equipped with a stir bar, magnesium ethoxide (11 mg, 0.10 mmol, 1 equiv) was added to a solution of 3-(ethylthio)-3-oxopropanoic acid (**S5**, 30 mg, 0.20 mmol, 2 equiv) in THF (1.0 mL, 0.1 M). The reaction mixture was stirred for 6 h at 23 ºC, and then concentrated *in vacuo*. The resulting magnesium salt of **S5** was dissolved in anhydrous DMF (0.5 mL) and transferred to the flask containing the acylimidazole derivative of **S10** via syringe. The reaction mixture was stirred overnight at 23 ºC. The reaction mixture was then diluted with 1M aqueous HCl (10 mL) and the resulting precipitate isolated by filtration. The crude solid was then dissolved in DMSO (2 mL) and purified by reverse-phase preparative HPLC using a Hypersil Gold™ aQ C18 polar endcapped column (250 × 20 mm, 5 µm particle size, Thermo Fisher Scientific). The following gradient was applied: 0-1 min, 50% B isocratic; 1-20 min, 50-100% B; 20-23 min, 100% B isocratic; 23-24 min, 100-50% B; 24-30 min, 50% B isocratic (solvent A: water + 0.1% formic acid; solvent B: acetonitrile + 0.1% formic acid; flow rate: 8 mL/min). The product containing fractions were pooled and concentrated to afford **10** (25 mg, 47% yield) as a white solid. $^{1}$H NMR (600 MHz, DMSO-$d_6$): δ (ppm) = 7.87 (d, *J* = 7.7 Hz, 1H), 7.49 (d, *J* = 8.4 Hz, 1H), 7.25 (br s, 1H), 6.84 (br s, 1H), 4.42 (q, *J* = 7.0 Hz, 1H), 3.85 - 3.74 (m, 2H), 3.73 - 3.62 (m, 1H), 2.83 (q, *J* = 7.3 Hz, 2H), 2.50 - 2.42 (m, 2H), 2.42 (dd, *J* = 15.1, 5.7 Hz, 1H), 2.34 (dd, *J* = 15.1, 7.6 Hz, 1H), 2.09 (t, *J* = 6.6 Hz, 2H), 1.65 - 1.55 (m, 1H), 1.53 - 1.41 (m, 3H), 1.23 (s, 20H), 1.17 (t, *J* = 7.4 Hz, 3H), 0.99 (d, *J* = 6.5 Hz, 3H), 0.85 (t, *J* = 6.5 Hz, 3H). $^{13}$C NMR (126 MHz, DMSO-$d_6$)*: δ (ppm) = 202.5, 192.3, 172.1, 171.4, 170.5, 56.9, 49.9, 43.5, 37.3, 35.2, 31.3,

29.6, 29.06, 29.01, 28.96, 28.87, 28.71, 28.68, 25.2, 23.1, 22.1, 20.5, 14.5, 13.9. HRMS (ESI): calcd for $C_{27}H_{50}N_3O_5S$ $[M+H]^+$, 528.3466; found, 528.3469. $[\alpha]_D^{23}$ = + 10.8° (*c* = 1.0, DMSO-$d_6$).

*24 distinct $^{13}$C signals are observed for this substrate (27 are expected), which is likely due to the close overlap of peaks from the myristoyl group in the 29.1 - 28.5 ppm range.

Preparation of thioester **S7**

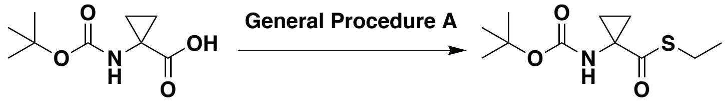


**S4** **S7**

Thioester **S7** was prepared according to General Procedure A using *N*-(*tert*-butoxycarbonyl)-1-amino-1-cyclopropane carboxylic acid (**S4**, 100 mg, 0.50 mmol, 1 equiv). The resulting residue was purified by flash column chromatography (0% EtOAc/Hexane to 30% EtOAc/Hexane) on silica gel to afford compound **S7** as a colorless oil (78 mg, 64% yield). $^1$H NMR (500 MHz, $CDCl_3$): δ (ppm) = 5.33 (s, 1H), 2.83 (q, *J* = 7.4 Hz, 2H), 1.62 - 1.57 (m, 2H), 1.45 (s, 9H), 1.23 - 1.16 (m, 5H). $^{13}$C NMR (126 MHz, $CDCl_3$): δ (ppm) = 201.3, 155.3, 80.5, 42.0, 28.5, 23.8, 20.6, 14.5. HRMS (ESI): calcd for $C_{11}H_{19}NO_3SNa^+$ $[M+Na]^+$, 268.0983; found, 268.0975.

Preparation of thioester **11**

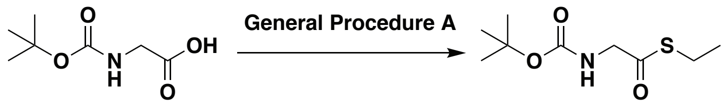


**S11** **11**

Thioester **11** was prepared according to General Procedure A using *N*-(*tert*-butoxycarbonyl)-glycine (**S11**, 200 mg, 1.14 mmol, 1 equiv). The resulting residue was purified by flash column chromatography (0% EtOAc/Hexane to 20% EtOAc/Hexane) on silica gel to afford compound **11** as a colorless oil (178 mg, 71% yield). $^1$H NMR (500 MHz, $CDCl_3$): δ (ppm) = 5.18 (br s, 1H), 4.01 (d, *J* = 2.7 Hz, 2H), 2.88 (q, *J* = 7.4 Hz, 2H), 1.43 (s, 9H), 1.23 (t, *J* = 7.4 Hz, 3H). $^{13}$C NMR (126 MHz, $CDCl_3$): δ (ppm) = 198.3, 155.6, 80.3, 50.3, 28.3, 23.0, 14.5. HRMS (ESI): calcd for $C_9H_{17}NNaO_3S^+$ $[M+Na]^+$, 242.0821; found, 242.0826.

Preparation of thioester **S8**

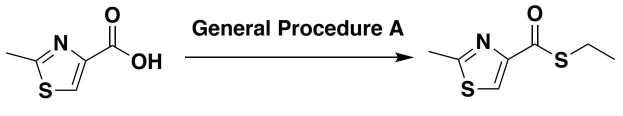


**S12** **S8**

Thioester **S8** was prepared according to General Procedure A using 2-methylthiazole-4-carboxylic acid (**S12**, 100 mg, 0.70 mmol, 1 equiv). The resulting residue was purified by flash column chromatography (0% EtOAc/Hexane to 30% EtOAc/Hexane) on silica gel to afford compound **S8** as a light yellow oil (75 mg, 57 % yield). $^{1}$H NMR (500 MHz, $CDCl_3$): δ (ppm) = 7.95 (s, 1H), 3.05 (q, *J* = 7.4 Hz, 2H), 2.77 (s, 3H), 1.35 (t, *J* = 7.5 Hz, 3H). $^{13}$C NMR (126 MHz, $CDCl_3$): δ (ppm) = 186.1, 166.6, 152.7, 122.6, 23.1, 19.4, 14.6. HRMS (ESI): calcd for $C_7H_{10}NOS_2^+$ $[M+H]^+$, 188.0198; found, 188.0204.

### 5. NMR spectra

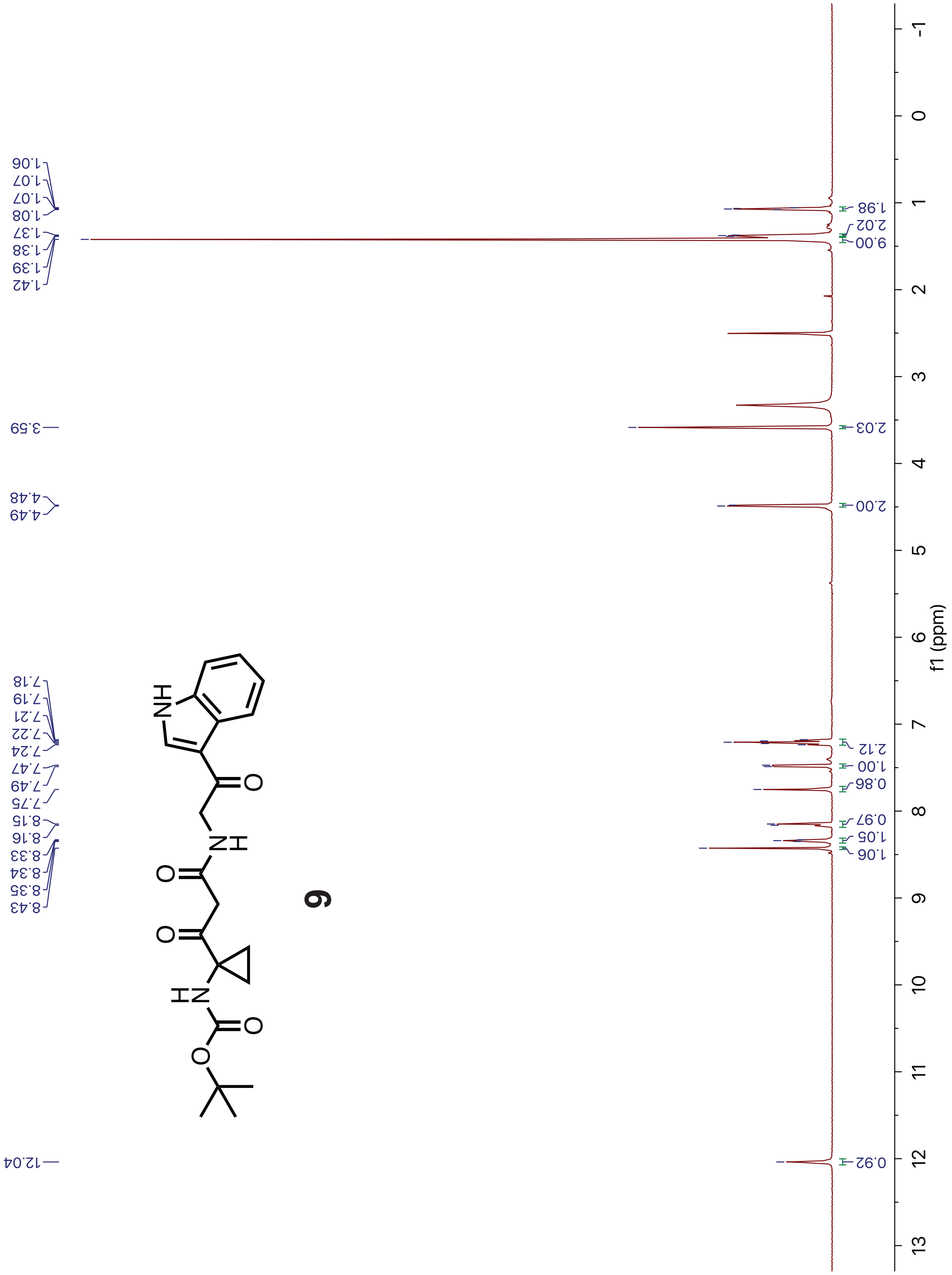


$^{1}$H spectrum of **9** (recorded in DMSO-$d_6$ at 500 MHz).

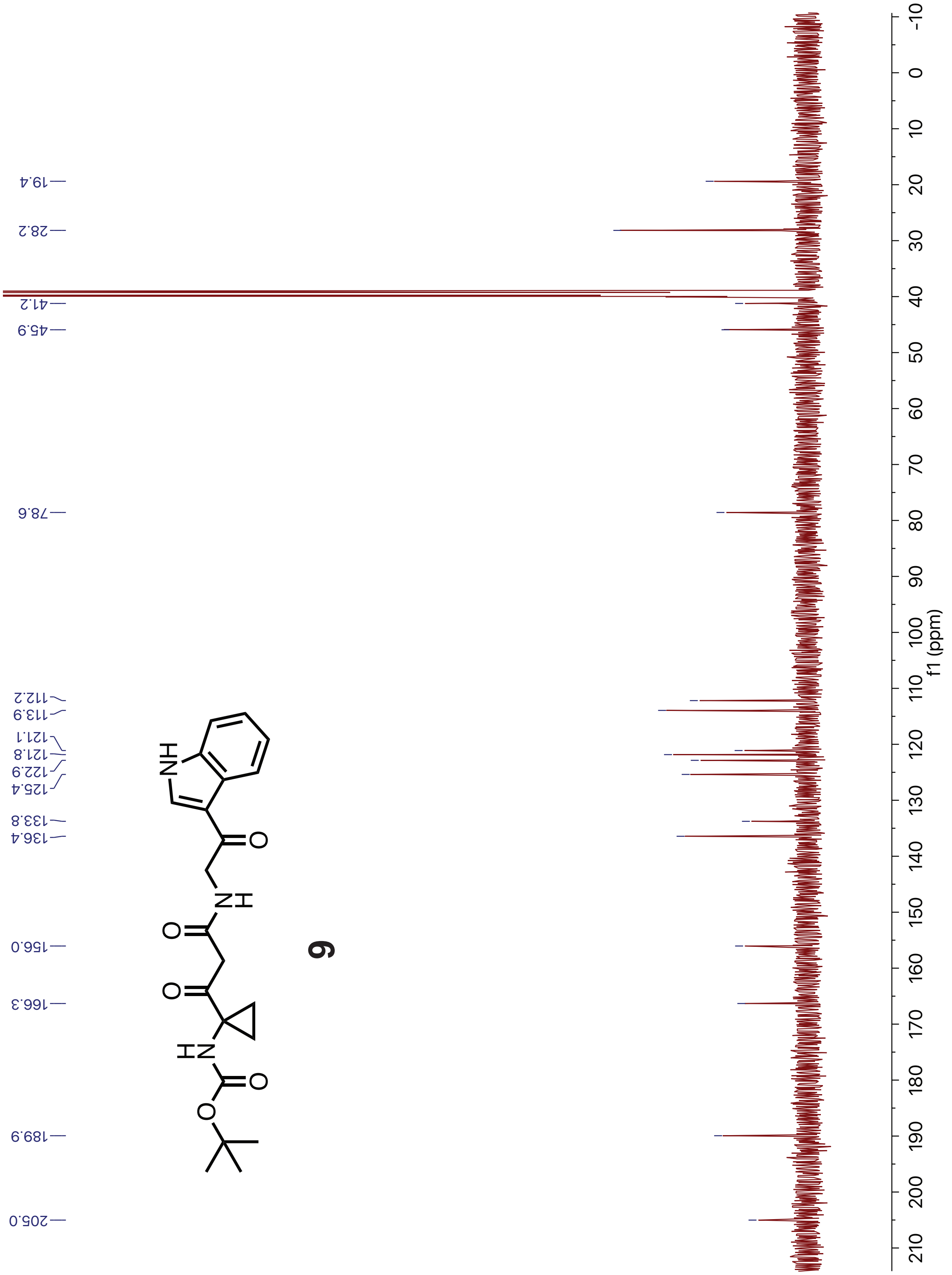


$^{13}$C spectrum of **9** (recorded in DMSO-$d_6$ at 126 MHz).

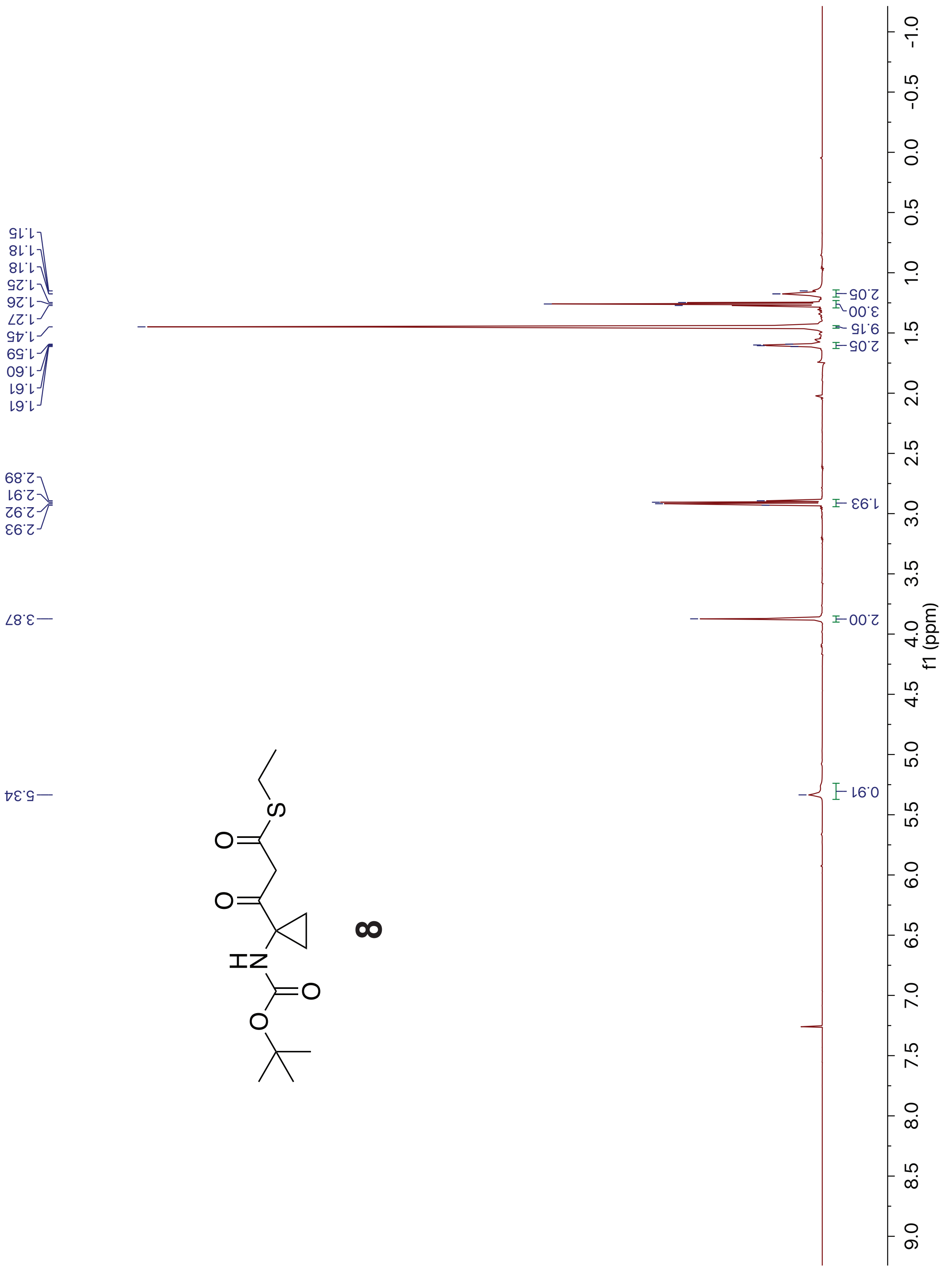


[1]H spectrum of **8** (recorded in $CDCl_3$ at 600 MHz).

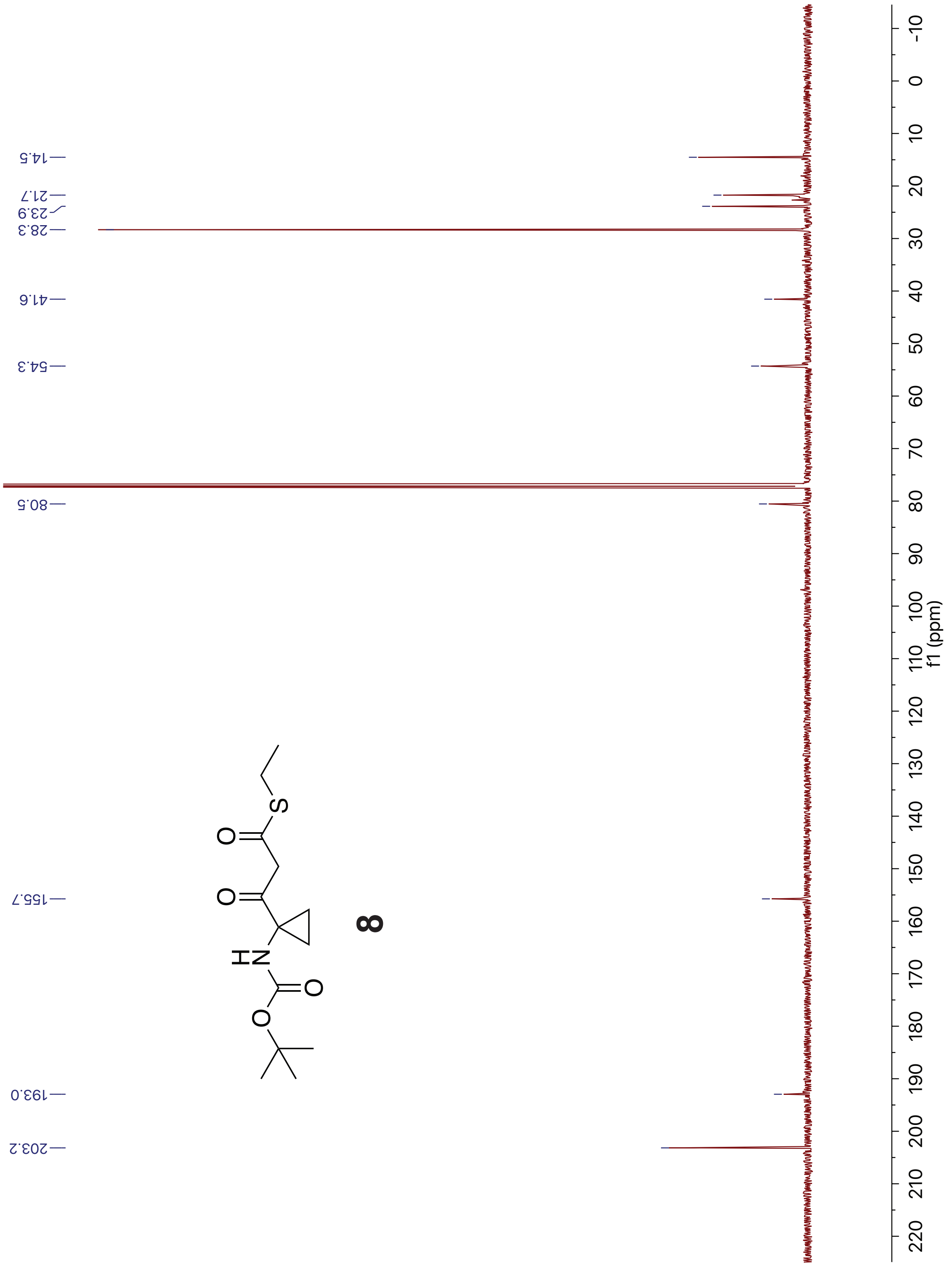

$^{13}C$ spectrum of **8** (recorded in $CDCl_3$ at 126 MHz).

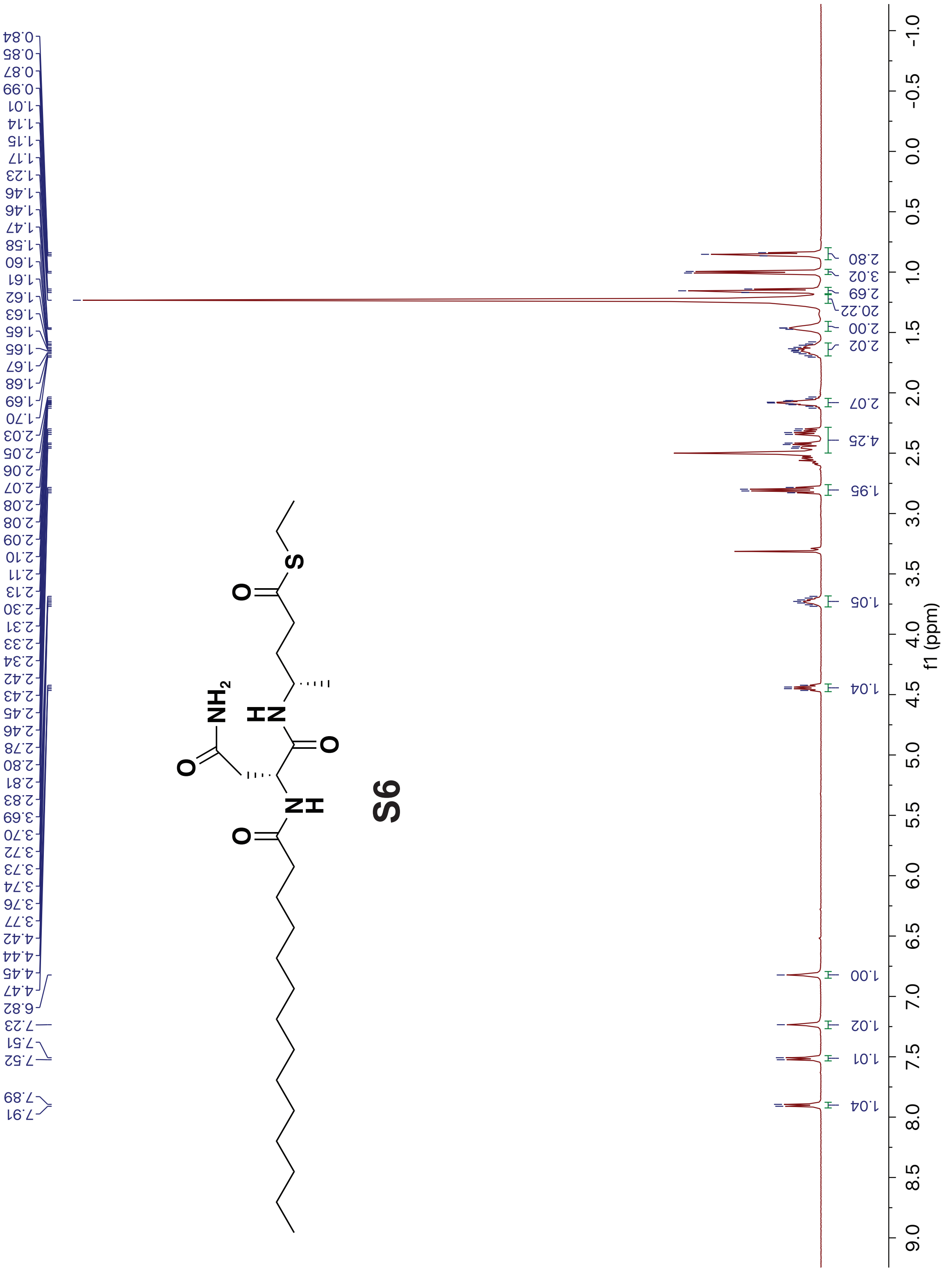


[1]H spectrum of **S6** (recorded in DMSO-$d_6$ at 500 MHz).

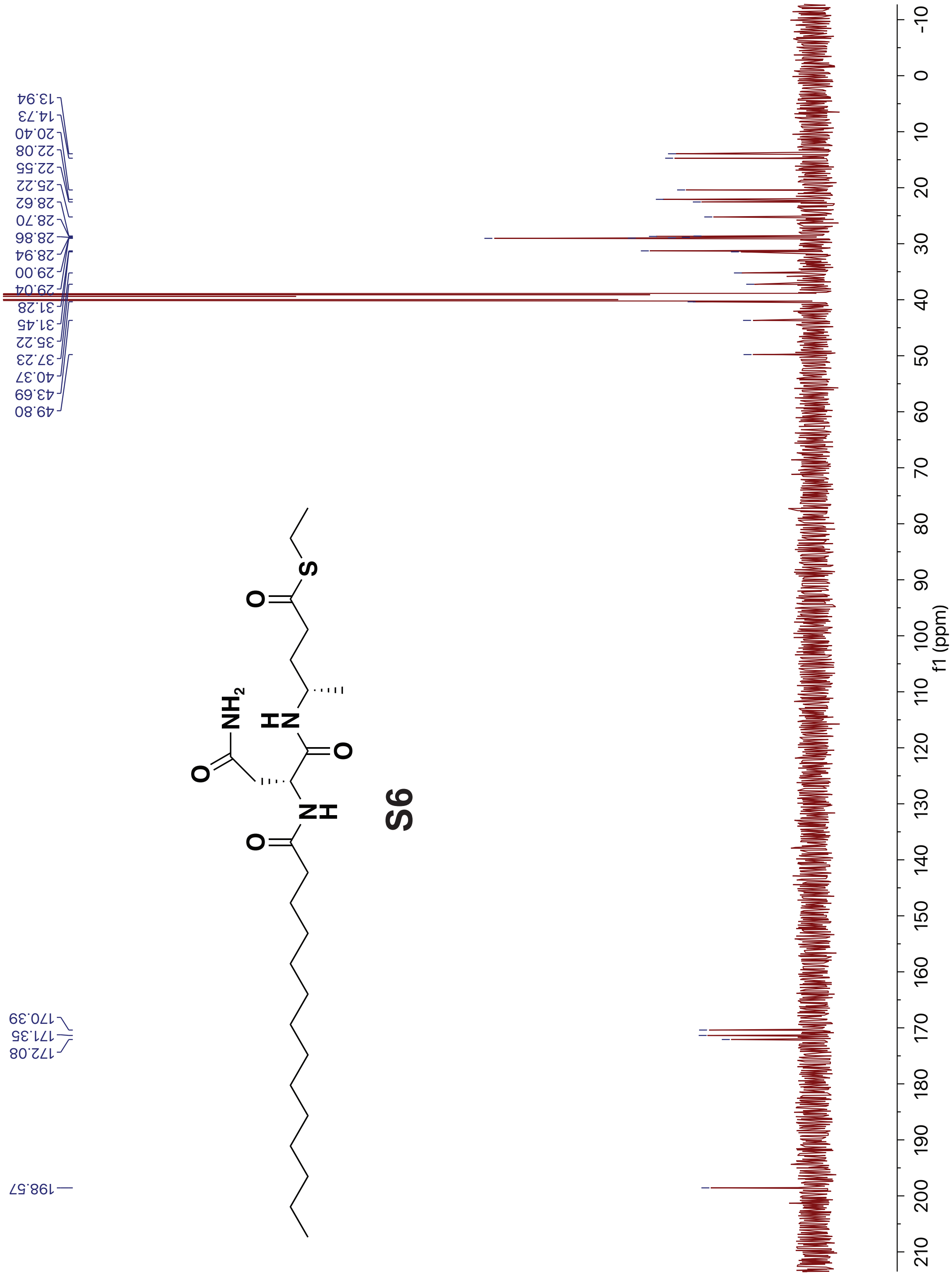


$^{13}$C spectrum of **S6** (recorded in DMSO-$d_6$ at 126 MHz).

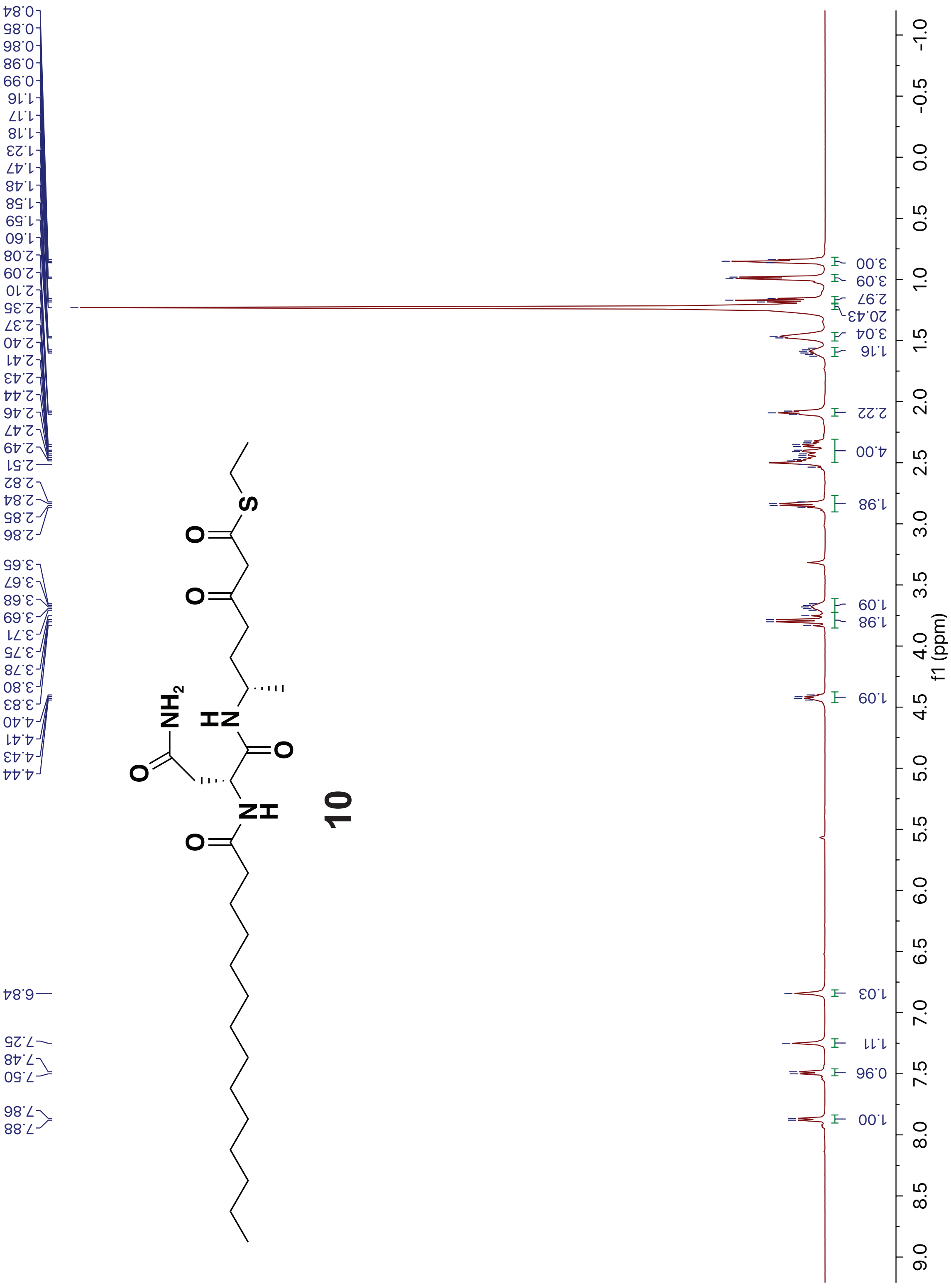


[1]H spectrum of **10** (recorded in DMSO-$d_6$ at 500 MHz).

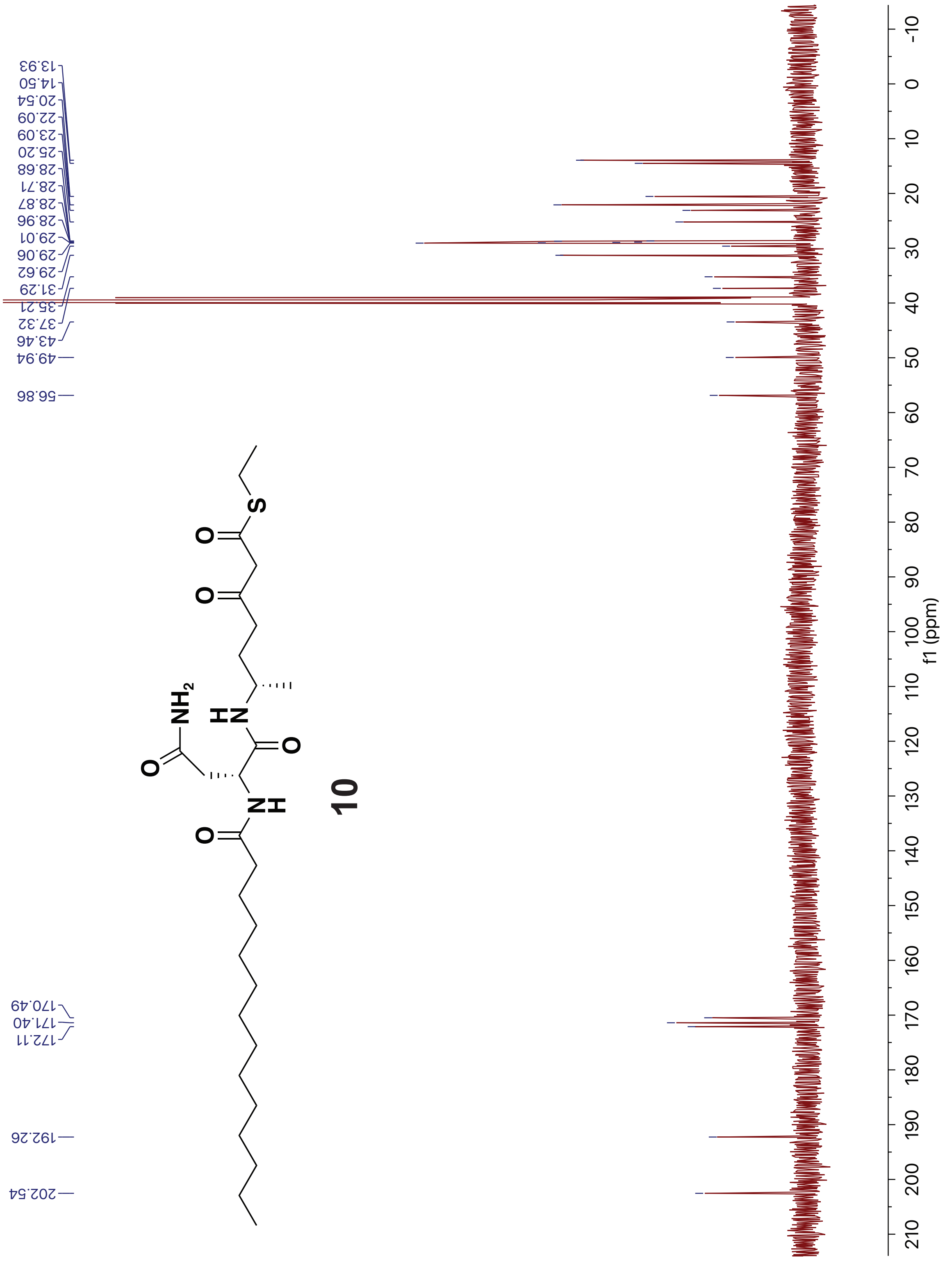


$^{13}$C spectrum of **10** (recorded in DMSO-$d_6$ at 126 MHz).

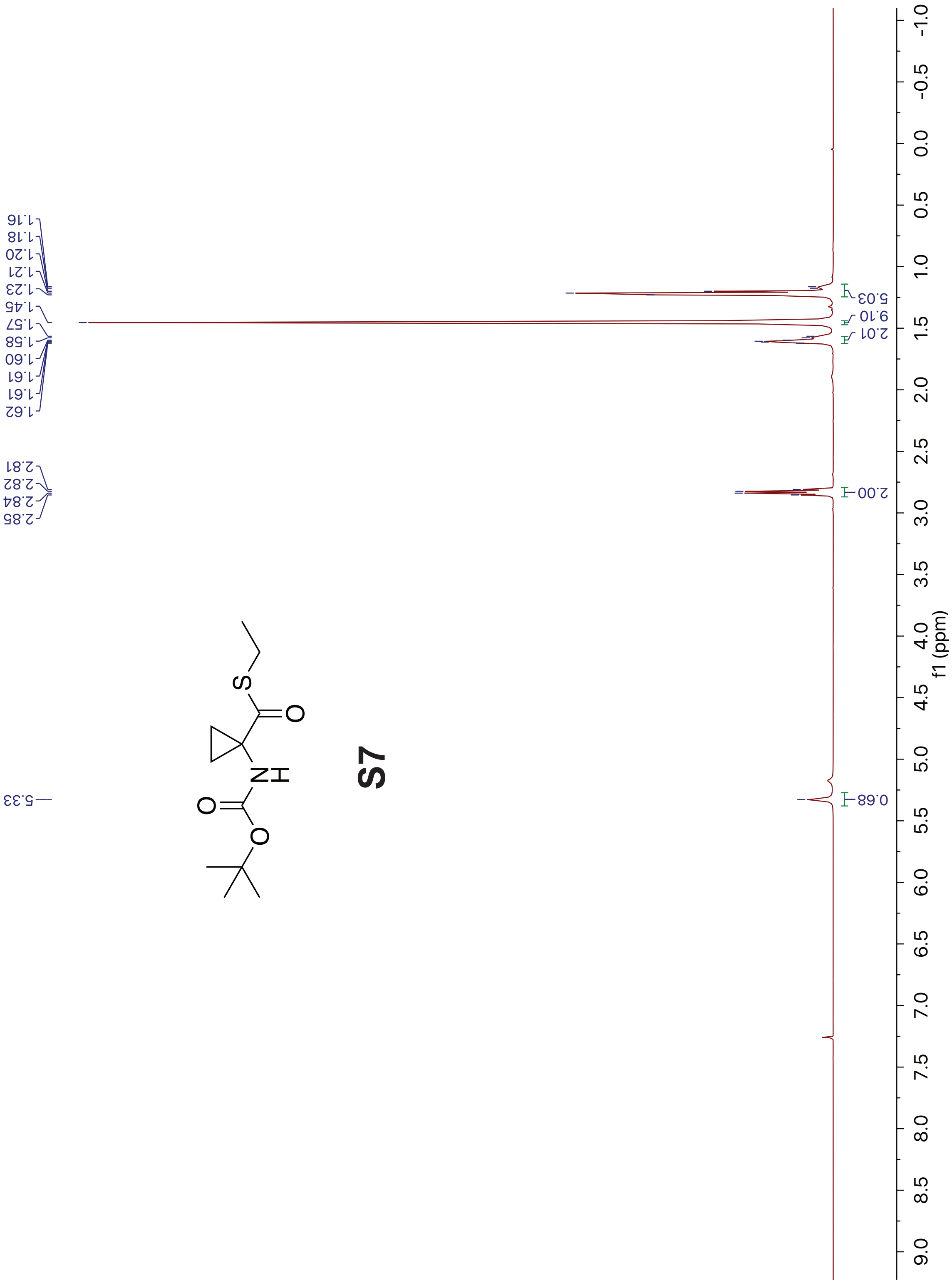


[1]H spectrum of **S7** (recorded in $CDCl_3$ at 500 MHz).

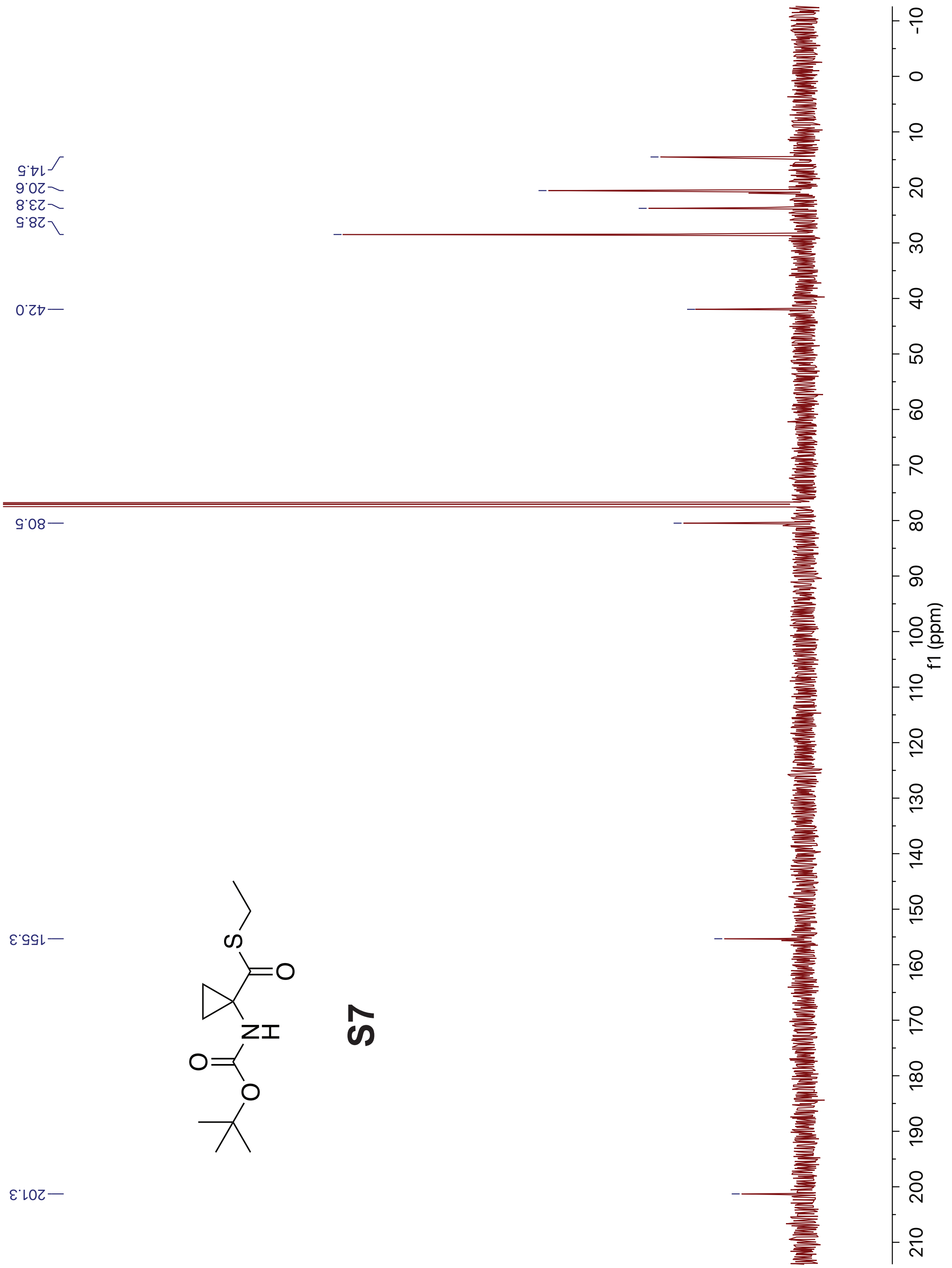


$^{13}$C spectrum of **S7** (recorded in $CDCl_3$ at 126 MHz).

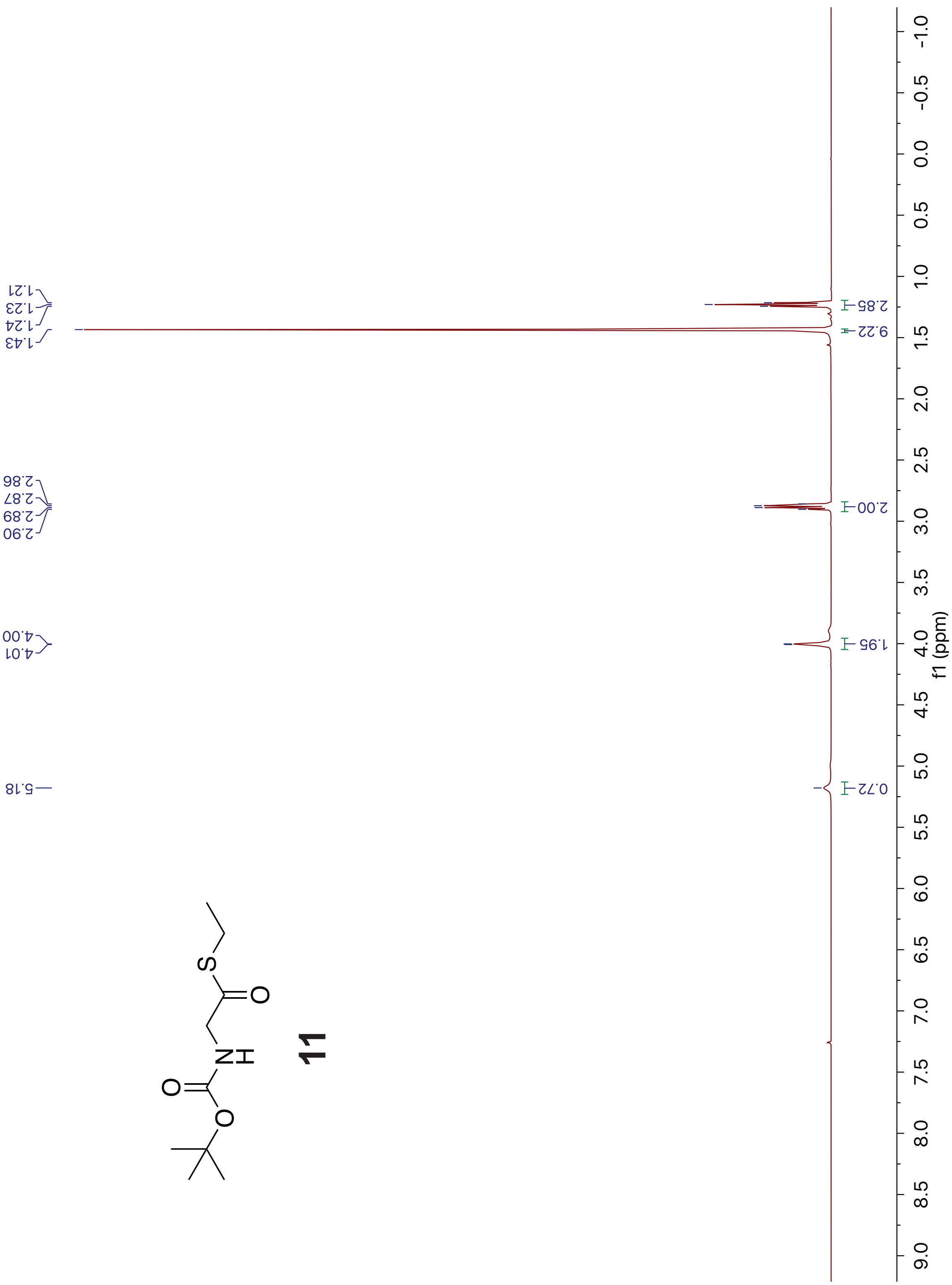


[1]H spectrum of **11** (recorded in $CDCl_3$ at 500 MHz).

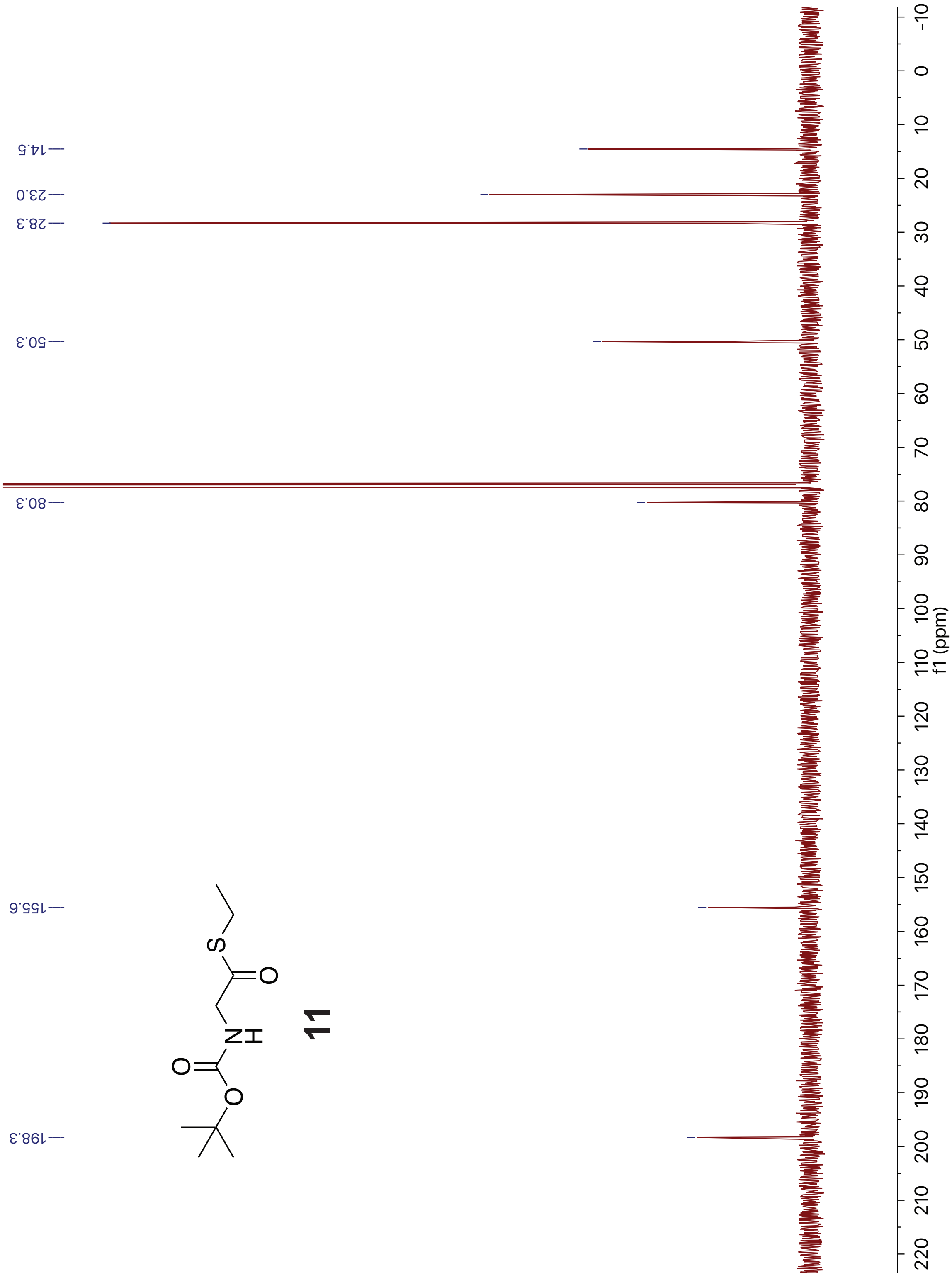


$^{13}C$ spectrum of **11** (recorded in $CDCl_3$ at 126 MHz).

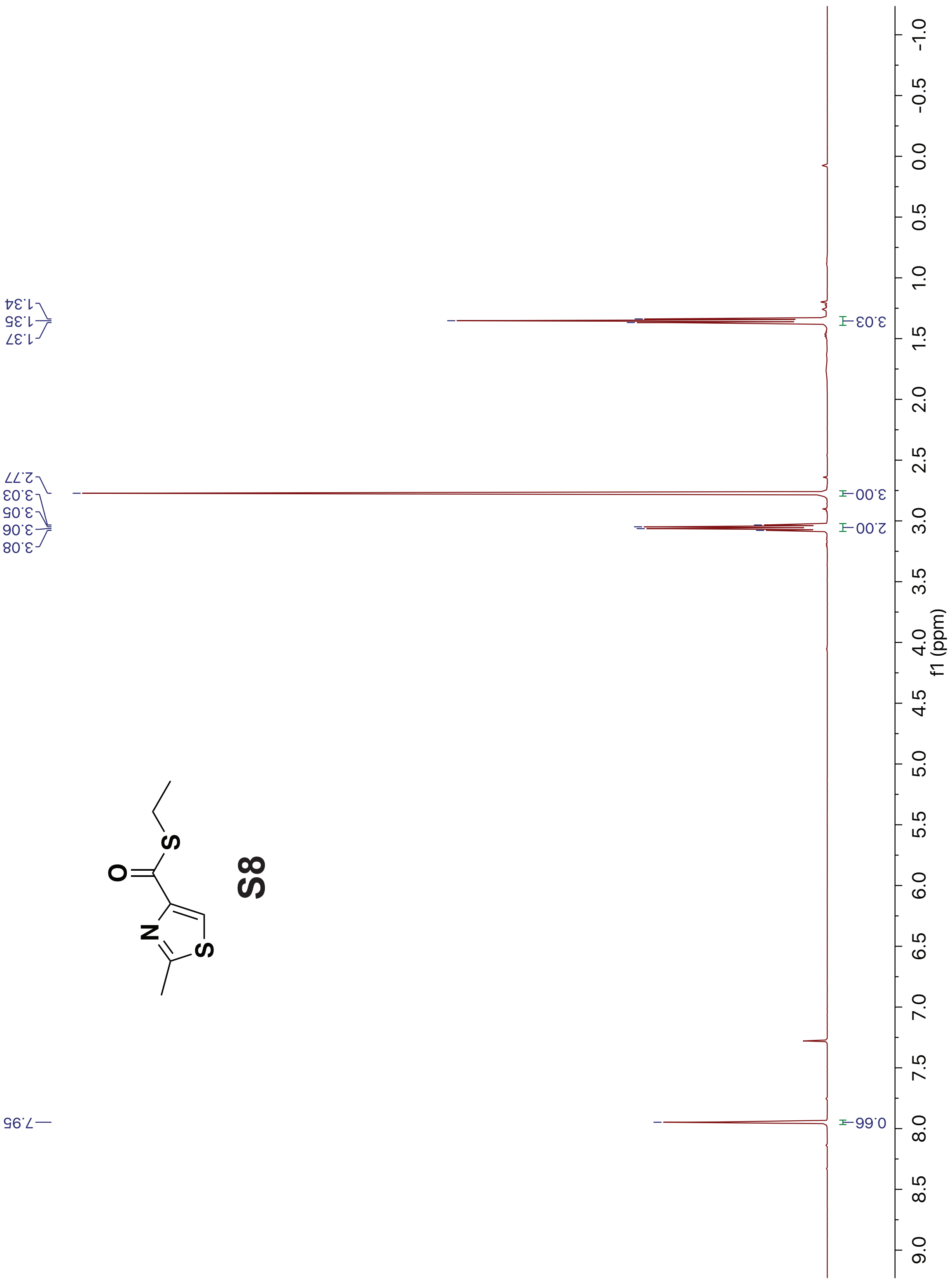


[1]H spectrum of **S8** (recorded in $CDCl_3$ at 500 MHz).

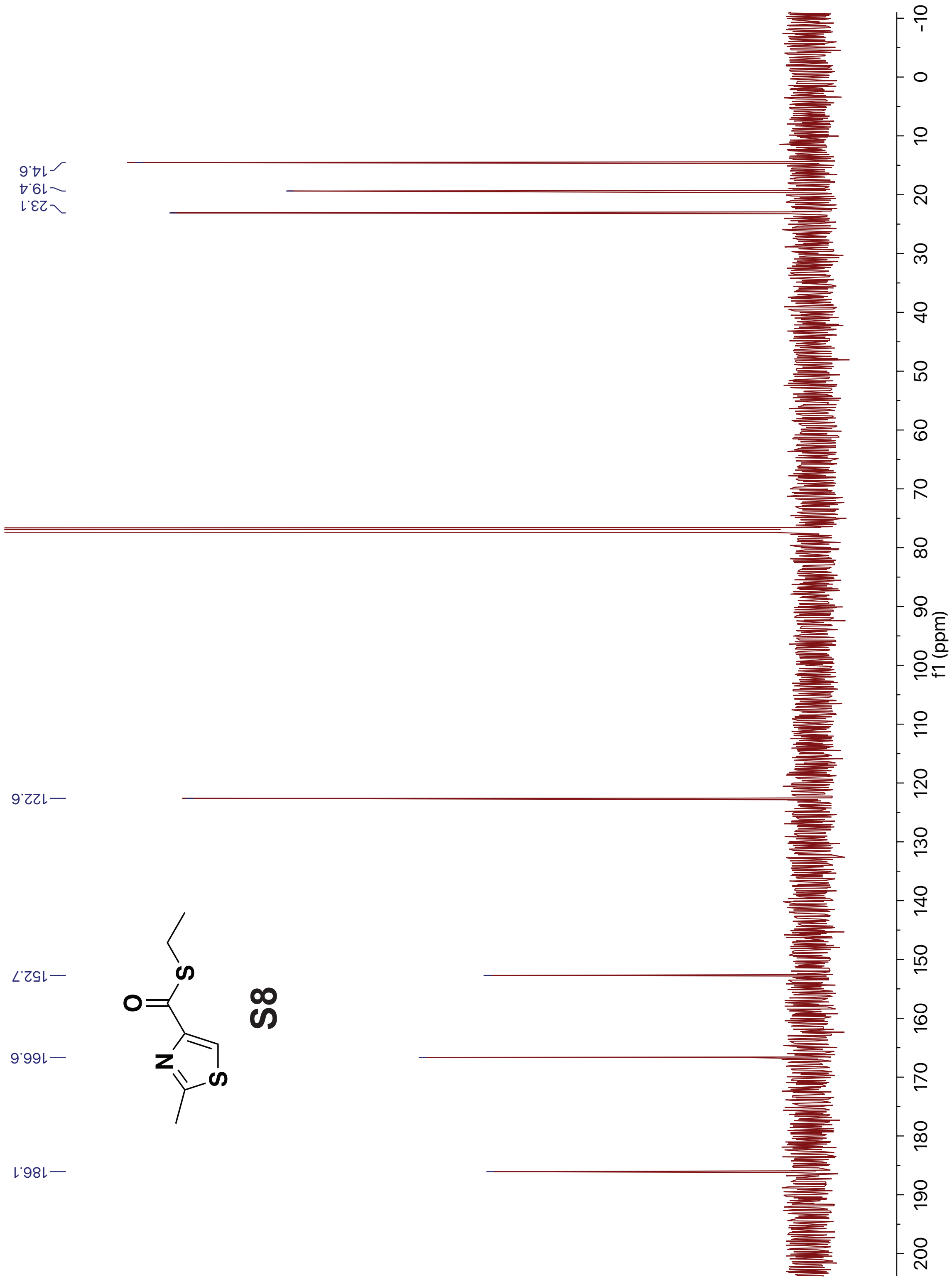


[13]C spectrum of **S8** (recorded in $CDCl_3$ at 126 MHz).
